# Discovery of Small Molecules and a Druggable Groove That Regulate DNA Binding and Release of the AP1 Transcription Factor ΔFOSB

**DOI:** 10.1101/2025.10.21.683675

**Authors:** Sean McNeme, Yun Young Yim, Ashwani Kumar, Yi Li, Brandon Hughes, Corey Peyton St. Romain, Galina Aglyamova, Jianping Chen, Nghi D. Nguyen, Shanghua Fan, Gabriel S. Stephens, Wen-Ning Zhao, Samantha Kruzshak, Molly Estill, Corrine Brener, Solange Tofani, Anil Kumar, Earnest P. Chen, Nadeen Takatka, Alfred J. Robison, Haiying Chen, Reid T. Powell, Stephen J. Haggarty, Clifford Stephan, Eric J. Nestler, Jeannie Chin, Mischa Machius, Jia Zhou, Gabby Rudenko

**Author notes:** Joint First Authors (Sean McNeme, Yun Young Yim, Ashwani Kumar, Yi Li). Corrine Brener: Hackensack Meridian School of Medicine, 340 Kingsland Street, Nutley, New Jersey, 07110, USA; Jianping Chen: Shanghai Institute of Materia Medica, Chinese Academy of Sciences, Pudong, Shanghai, 201203, China; Samantha Kruzshak: Department of Chemical and Biological Engineering, Tufts University, Medford, Massachusetts, 02155, USA; Ashwani Kumar: Department of Physiology & Biophysics, Case Western Reserve University, Cleveland, Ohio, 44106, USA; Yi Li: Shanghai Institute of Materia Medica, Chinese Academy of Sciences, Pudong, Shanghai 201203, China; Galina Aglyamova: Department of Biology, University of Houston, Houston, Texas, 77204, USA; Gabriel S. Stephens: Center for Dementia Research, Nathan Kline Institute for Psychiatric Research, Orangeburg, New York, 10962, USA; Solange Tofani: University of Hawaiʻi at Mānoa, Honolulu, Hawaiʻi, 96822, USA; Wen-Ning Zhao: K2B Therapeutics, Cambridge, Massachusetts, 02142, USA.

## Abstract

ΔFOSB, a member of the AP1 family of transcription factors, mediates long-term neuroadaptations underlying drug addiction, seizure-related cognitive decline, dyskinesias, and several other chronic conditions. AP1 transcription factors are notoriously difficult to modulate pharmacologically due to the absence of well-defined binding pockets. Here, we identify a novel site on ΔFOSB, located outside the DNA-binding cleft, that accommodates small molecules. We show that sulfonic acid-containing compounds bind to this site via an induced-fit mechanism, reorienting side chains critical for DNA binding, and that they may hinder the ΔFOSB bZIP α-helix from binding to the major groove of DNA. *In vivo*, direct administration of one such compound, JPC0661, into the brain reduces ΔFOSB occupancy at genomic AP1 consensus sites by approximately 60% as determined by CUT&RUN-sequencing. These findings suggest that DNA binding and release by AP1 transcription factors can be controlled via small molecules that dock into a novel site that falls outside of the DNA-binding cleft. Minimal sequence conservation across 29 bZIP domain-containing transcription factors in this druggable groove suggests that it can be exploited to develop AP1-subunit-selective compounds. Our studies thus reveal a novel strategy to design small-molecule inhibitors of ΔFOSB and other members of the bZIP transcription factor family.

## Introduction

The AP1 transcription factor, ΔFOSB, is a promising drug target for several neurological and psychiatric disorders, including drug addiction, seizure-induced cognitive decline, and dyskinesias. The ΔFOSB protein accumulates in response to chronic (but not acute) stimuli and insults in specific regions of the brain (1). Accumulation of ΔFOSB in these brain regions alters the expression of specific sets of genes, inducing a range of long-term neural and behavioral adaptations. The ΔFOSB protein is unusually stable with a half-life of ∼7 days in the brain, while that of FOSB and all other FOS family proteins is several hours at most (2–4).

ΔFOSB protein levels rise in the striatum following chronic exposure to drugs of abuse and lead to increased reward-seeking and reward-reinforcing behaviors (1, 5). ΔFOSB protein levels also dramatically increase in the hippocampus in individuals with Alzheimer’s disease (AD), mild cognitive impairment (a prodromal form of AD), several forms of epilepsy (6–8), as well as in the hippocampus of mouse models of epilepsy or AD neuropathology that express elevated amounts of amyloid precursor protein (APP)-derived amyloid-β peptides (6, 7). Similarly, high levels of ΔFOSB protein are induced in the striatum following dyskinesias (abnormal involuntary movements), such as seen in individuals with Parkinson’s disease treated with dopaminergic agonists (9, 10).

Importantly, inhibiting ΔFOSB within the relevant brain regions of animal models for drug addiction, Alzheimer’s disease, or Parkinson’s disease, through genetic or viral means, reverses neural and behavioral maladaptations seen in these models. For instance, decreasing ΔFOSB levels specifically in the nucleus accumbens reduces drug-addiction behaviors (5), whereas a broader reduction in the striatum diminishes L-DOPA-induced dyskinesias observed in Parkinson’s disease (10). Also, inhibiting ΔFOSB activity by adenoviral administration of a defective partner, ΔJUND, improves spatial memory and other cognitive measures in an AD mouse model (6, 7) and alleviates L-DOPA-induced dyskinesias in a non-human primate Parkinson’s disease model (11).

Despite these encouraging findings, extrapolating results from genetic and viral manipulations to therapeutic applications remains challenging. Such manipulations are typically applied locally to specific brain regions or even specific cell types in those regions, and the interpretation of genetic manipulations is further complicated by possible compensatory adaptations occurring during development. Adding to the complexity, the molecular composition of ΔFOSB-containing complexes *in vivo* is likely variable, with ΔFOSB recruiting a diverse array of binding partners that differ across specific brain regions, cell types, and conditions (1). Therefore, the exact physiological roles of ΔFOSB remain unclear, and harnessing its therapeutic potential as a drug target is challenging. Pharmacological probes that target ΔFOSB *in vivo*, either by upregulating or downregulating its action, would be enormously useful to interrogate ΔFOSB function, its roles in different disease pathologies, and its utility as a therapeutic target.

AP1 transcription factors, such as ΔFOSB, have typically been considered ‘undruggable’ because they lack obvious molecular features that can be targeted with small molecules, and they are conformationally and compositionally highly dynamic. Key structure-function relationships have been revealed for ΔFOSB in recent years. The ΔFOSB protein (237 amino acids) is composed of an intrinsically disordered N-terminal region (∼150 amino acids) and a C-terminal basic region/leucine zipper (bZIP) that forms a DNA-binding site upon assembly with another AP1 bZIP partner (12–14). In the brain, a predominant binding partner of ΔFOSB is JUND (15, 16). Following dimerization of the ΔFOSB and JUND bZIP domains, which is mediated by the helical leucine-zipper region, the helical basic regions insert into the major groove of double-stranded DNA, recognizing the AP1 DNA palindromic consensus sequence (AP1 site; 5’-TGA C/G TCA-3’) (12, 13). Cysteine residues found N-terminal to the DNA-binding motifs (ΔFOSB Cys172 and JUND Cys285) form a putative redox switch that is conserved in the closely related AP1 transcription factor FOS/JUN (13, 17). In the DNA-bound form, the redox residues are kept far apart (14 Å), but in the absence of DNA and under oxidizing conditions, they can form a disulfide bond that kinks ΔFOSB (but not JUND), malforming the DNA-binding site (13). ΔFOSB homomers can also form, at least *in vitro*, via a non-canonical arrangement. They selectively recognize AP1-consensus DNA sites but appear not to be under redox control, and their *in vivo* role is not known (14).

The unusual stability of ΔFOSB stems in part from the lack of the C-terminal 101 amino acids found in FOSB, which carry a transactivation domain as well as degron sequences (5). Strikingly, ΔFOSB can increase or decrease the expression of various target genes, although the molecular bases of these opposing actions are not well understood (1, 18). Recent *in vivo* studies of medium spiny neurons in mouse nucleus accumbens revealed that, while most (93%) ΔFOSB is bound to AP1-sites as expected, the majority of these sites (85%) are located outside traditional gene promoter regions, instead mapping to putative enhancer sites along intergenic and intronic regions of the mouse genome (18). It was also observed that 50% of the ΔFOSB-binding sites that did map to gene promoters did not bear the histone marks for a gene promoter undergoing transcriptional activation, suggesting that these might be ΔFOSB-containing complexes involved in the repression of gene transcription (18). Thus, despite significant progress in elucidating the structure-function relationships of ΔFOSB, few novel vulnerabilities have emerged that seem suitable for targeting this transcription factor with small molecules, except for the redox switch (19).

Here, we demonstrate that ΔFOSB, a prototypical AP1 transcription factor, can be pharmacologically regulated via small-molecule compounds binding to a novel druggable groove. We used high-throughput screening (HTS) to identify compounds that disrupt ΔFOSB/JUND heterodimeric and ΔFOSB homomeric complexes bound to DNA. We identified the compound 5-(3-amino-5-oxo-4,5-dihydro-1*H*-pyrazol-1-yl)-2-phenoxybenzenesulfonic acid, JPC0661, as a direct inhibitor of ΔFOSB/JUND and ΔFOSB DNA-binding biochemically, as an inhibitor of ΔFOSB-mediated transcription in cell-based assays, and as an inhibitor of ΔFOSB binding to AP1 consensus DNA-binding sites *in vivo* in an APP mouse model of AD neuropathology (Line J20). The crystal structure of the ΔFOSB/JUND bZIP domain in complex with JPC0661_comm_ revealed that the compound binds to a surface outside of the direct DNA-binding cleft, leading to a rearrangement of key DNA-binding residues within ΔFOSB. Importantly, the residues forming the broader compound-binding groove are not conserved among human AP1 transcription factors, suggesting that this novel site can be exploited to develop compounds that selectively target distinct AP1 transcription factors.

## Results

### Identifying Novel Inhibitors of ΔFOSB

To identify compounds that disrupt the binding of ΔFOSB/JUND heterodimers (‘ΔFOSB/JUND’) and ΔFOSB homomers (‘ΔFOSB’) to DNA, we screened a total of 44,480 compounds from the Chembridge NT638 Diversity Set 1 (30,080 compounds) and Maybridge (14,400 compounds) libraries using a fluorescence polarization (FP) assay adapted for high-throughput screening with a TAMRA-labeled oligonucleotide carrying an AP1 consensus site (‘AP1-oligo’, a.k.a. ‘TMR-*cdk5*’). Hits were defined as compounds causing disruption of the protein:DNA complex for either ΔFOSB/JUND or ΔFOSB proteins, using broad cut-offs for the signal (75-125%). These compounds were retested to confirm their hit status, resulting in the identification of 123 compounds. After removing potentially problematic chemotypes using the PAINS database (20), 47 candidate scaffolds were manually chosen based on their chemical structures. The corresponding compounds were purchased, and FP-based dose-response curves (FP-DRC) were recorded as a confirmation step. Of these, two compounds, JPC0661_comm_ and HTS10307, were chosen for follow-up (**Fig. 1A**). JPC0661_comm_ is known commercially as JFD00458. We use the term ‘JPC0661_comm_’ to indicate when the commercially sourced compound was used for studies and the term ‘JPC0661’ to refer to the compound in general and to the compound that we resynthesized in-house for purity validation, as well as further cell-based studies, and *in vivo* studies. JPC0661_comm_ and HTS10307 inhibited the binding of ΔFOSB/JUND and ΔFOSB to DNA with IC_50_ values in the micromolar range (IC_50_ ∼9 μM and IC_50_ ∼52 μM for JPC0661_comm_; IC_50_ ∼3 μM and IC_50_ ∼7 μM for HTS10307, respectively) (**Fig. 1B**). Of note, both compounds were less active in the presence of 1 mM TCEP, a reductant, decreasing the activity of JPC0661_comm_ and HTS10307 against ΔFOSB/JUND four to five-fold but less than two-fold against ΔFOSB (**Fig. 1C**). We confirmed the ability of JPC0661_comm_ and HTS10307 to disrupt the binding of ΔFOSB/JUND and ΔFOSB to the AP1-oligonucleotide via electrophoretic mobility assays (EMSAs) (**Fig. 1D**). In these EMSAs, the presence of TCEP also greatly diminished the activity of the compounds as tested for ΔFOSB/JUND (**Fig. 1D**), again discussed below.

**Figure 1.**
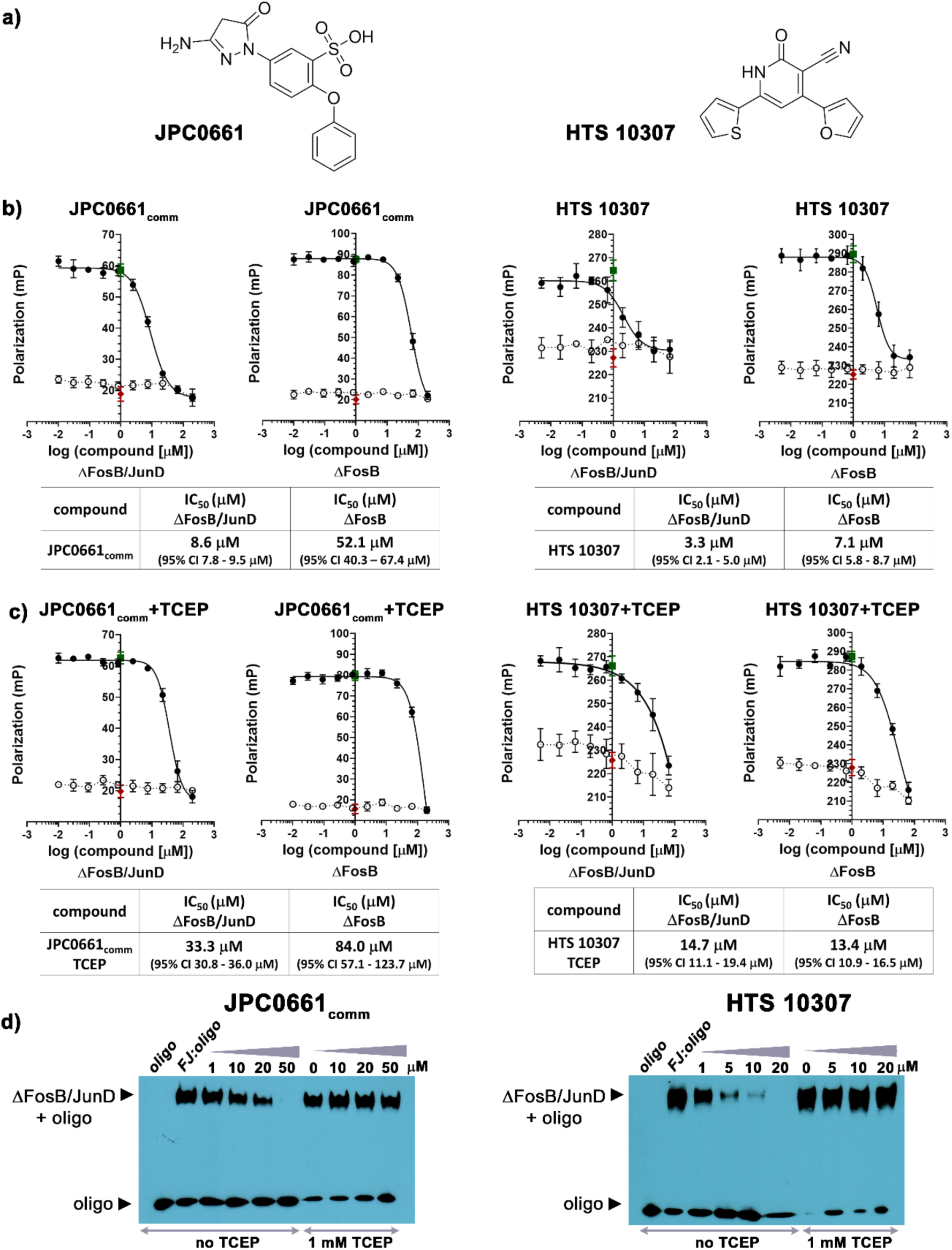
Compounds targeting ΔFOSB/JUND and ΔFOSB. **A)** Chemical structures of JPC0661 and HTS10307; **B)** JPC0661_comm_ and HTS10307 disrupt the binding of a TAMRA-labeled AP1-oligonucleotide (TMR-*cdk5*) in FP-DRC assays; **C)** Addition of 1 mM TCEP significantly reduces the activity of JPC0661_comm_ and HTS10307 in FP-DRC assays for ΔFOSB/JUND, though much less for ΔFOSB. For **B)** and **C)**, 280 nM ΔFOSB/JUND full-length protein (•; n=4), 320 nM ΔFOSB full-length protein (•; n=4) or no protein (o; n=4) was incubated with 25 nM TMR-*cdk5* oligonucleotide (oligo) and increasing amounts of compound (0-200 μM). Controls for ‘100%-inhibition’ (TMR-*cdk5* oligo alone; ♦; n=16) and ‘0%-inhibition’ (ΔFOSB/JUND+TMR-*cdk5* oligo; ▪; n=16) are indicated. Data points show the mean of n replicates, and error bars indicate the standard deviation (SD). **D)** The activity of JPC0661_comm_ and HTS10307 was validated orthogonally using electrophoretic mobility shift assays (EMSAs) by incubating ΔFOSB/JUND with a biotinylated-*cdk5* oligo (BIO-*cdk5*) and increasing amounts of compound (1-50 μM) with or without 1 mM TCEP. BIO-*cdk5* alone (‘oligo’) and the starting amount of ΔFOSB/JUND:BIO-*cdk5* complex in the absence of compound (‘FJ:oligo’) are shown as well.

Before evaluating the activity of our compounds in cell-based assays and *in vivo*, we assessed their toxicity and metabolic stability (**Fig. 2**). JPC0661_comm_ and HTS10307 were well tolerated in human neural progenitor cells (NPC), with no discernable decrease in cell viability after 24 h (0-20 μM) and only minimal losses after 72 h (<25 % at the highest dose of 20 μM) (**Fig. 2A**). In mouse liver microsomal stability assays, JPC0661_comm_ was remarkably stable (t_1/2_ = 51.2 min), however, HTS10307 was rapidly metabolized (t_1/2_ = 9.6 min) (**Fig. 2B**). Given its promising pharmacokinetic properties, we therefore selected JPC0661 as our lead compound.

**Figure 2.**
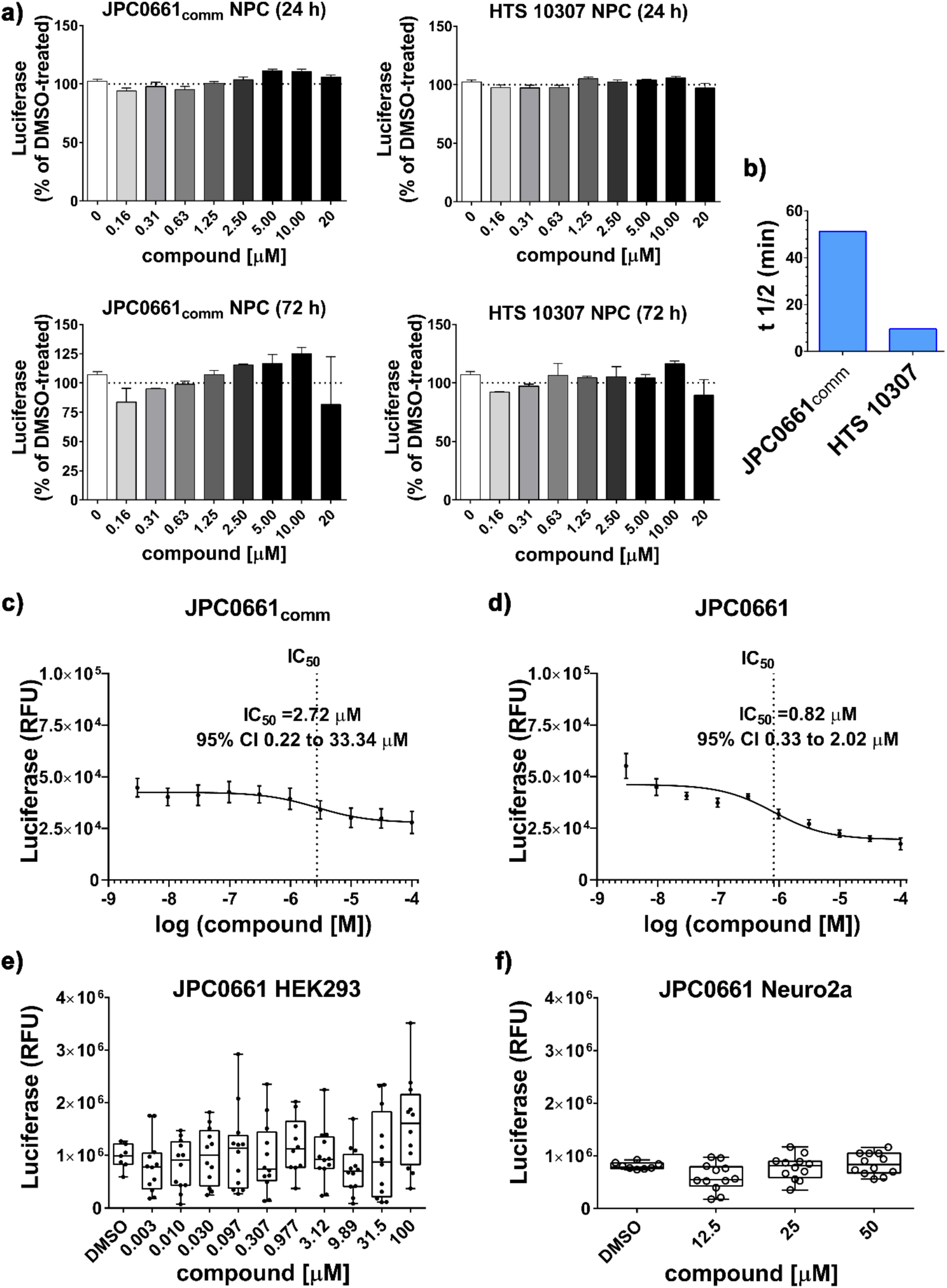
Cellular activity of JPC0661. **A)** Cell viability of JPC0661_comm_ and HTS10307 assessed in the range of 0-20 μM after 24 h (top) and 72 h (bottom) in neuronal progenitor cells (NPC) using the CellTiter-Glo viability assay. Cell viability measures are normalized to the negative control containing no compound but 0.6% DMSO. Dotted lines indicate 100% cell viability. The mean of n replicates is shown for 24 h (n=3) and 72 h (n=2) with error bars indicating the SEM. **B)** Stability of JPC0661_comm_ and HTS10307 in mouse liver microsomes. **C)** and **D)** Effects of JPC0661_comm_ (0–100 μM; **C**) and JPC0661 (0–100 μM; **D**) on AP1-driven luciferase activity in AP1-luc HEK293 cells. The dose-dependent activation of the AP1-driven luciferase reporter was measured based on changes in the luciferase signal and expressed as relative fluorescence units (RFU). Each compound was tested twice (n = 4 wells per experiment; total n = 6–8 wells for JPC0661_comm_ and n = 7–8 wells for JPC0661 per concentration), with results normalized to the luciferase signals of blank wells from corresponding experiments (n = 8 wells). Nonlinear regression using a three-parameter model was applied to fit the luciferase signal and calculate IC_50_ values, which are reported with a 95% confidence interval (CI), and data points are presented as the mean ± SEM. **E)** Effect of serially diluted JPC0661 (0.003-100 μM) on the viability of AP1-luc HEK293 cells using the Celltiter-Go viability assay (2 h). Cell viability was normalized to the DMSO control, which contained 0.5% DMSO but no compound (n = 7 wells). Data points are presented as the mean ± SEM, with a total n = 10–12 wells per concentration. F) Effect of JPC0661 (at doses 12.5, 25, and 50 μM) on the viability of Neuro 2A cells (72 h) using the Celltiter-Go viability assay to assess potential toxicity in mouse neuronal cells. Cell viability was normalized to the DMSO control, which contained 0.5% DMSO but no compound (n = 8 wells). Data points are presented as the mean ± SEM, with a total n = 12 wells per concentration.

To assess the ability of JPC0661 to regulate ΔFOSB-driven gene expression, we used HEK293 cells containing a stably integrated luciferase reporter under the control of an AP1-consensus sequence (AP1-luc HEK293 cells) (**Fig. 2C** and **2D**). ΔFOSB protein accumulates in these cells when they are first serum-starved (0.5% serum, 24 h) and then given high-serum conditions (20% serum, 24 h) (19). JPC0661_comm_ inhibited AP1 transcription factor-mediated expression of the luciferase reporter gene with IC_50_ ∼2.7 μM (**Fig. 2C**). To validate the identity of the compound and prepare large amounts for *in vivo* studies, we next synthesized an in-house version of JPC0661_comm_, yielding the compound JPC0661, which was chemically indistinguishable and similarly active in our FP-assay (**Supplementary Fig. S1**) as well as in our cell-based reporter assay with an IC_50_ ∼0.8 μM (**Fig. 2D**). JPC0661 was well tolerated in AP1-luc HEK293 cells (24 h exposure) and Neuro2A cells (72 h exposure) with no noticeable decrease in viability even at high doses up to 100 μM in CellTiter-Glo assays (**Fig. 2E** and **2F**).

Thus, we have identified a novel inhibitor of ΔFOSB, JPC0661, that disrupts DNA-binding directly, inhibits the ability of ΔFOSB to promote gene expression in cultured cells, and has favorable metabolic stability as well as low toxicity.

### JPC0661 Disrupts ΔFOSB Binding to DNA *in vivo*

To determine whether JPC0661 disrupts ΔFOSB binding to genomic DNA also *in vivo* in the brain, we performed CUT&RUN-sequencing, a high-resolution chromatin profiling method that maps genome-wide protein-DNA interactions. We profiled ΔFOSB binding in the dorsal hippocampus of APP transgenic mice following intracranial infusion of JPC0661 (“Treatment”) or vehicle (“Control”). APP mice overexpress mutant human APP and produce high levels of Aβ, which we have shown previously leads to robust induction of ΔFOSB in this brain region (7). JPC0661 or vehicle was delivered into the dorsal hippocampus via osmotic minipumps for three days, after which time the dorsal hippocampus was isolated and prepared for CUT&RUN-sequencing that was performed as described previously (18), with minor modifications. We detected 26,287 ΔFOSB-bound peaks in vehicle-treated mice, with JPC0661-treated animals exhibiting an approximately 60% reduction in total binding sites (10,132 peaks; **Fig. 3A**). A similar reduction was observed in the number of unique ΔFOSB-bound genes (**Supplementary Fig. S2A**). In addition to the peak number, average peak width was significantly decreased following JPC0661 treatment (**Fig. 3B**), indicating loss of binding stability and chromatin occupancy. These results show that JPC0661 directly disrupts ΔFOSB–DNA interactions *in vivo*, potentially altering downstream transcriptional regulation.

**Figure 3.**
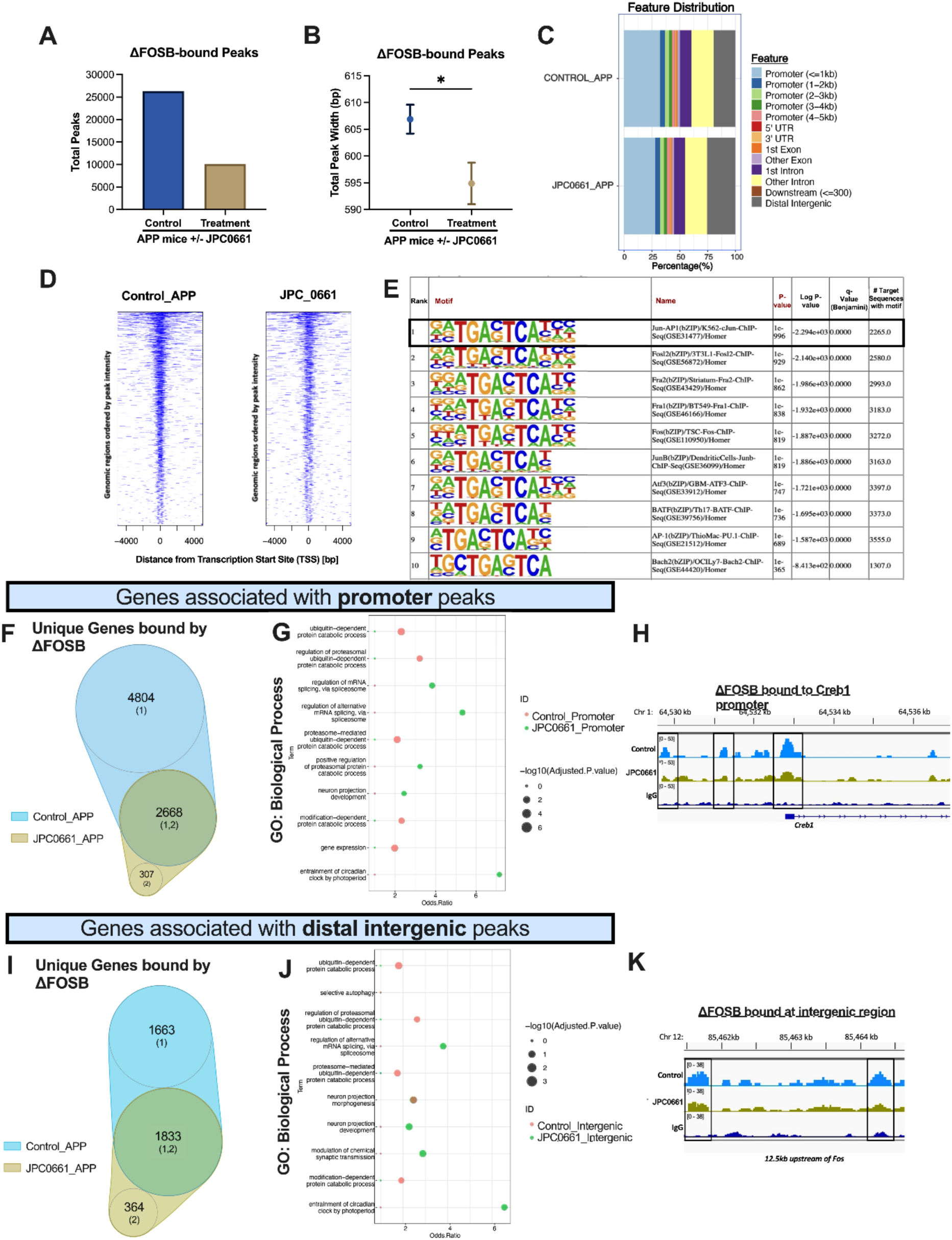
Effect of JPC0661 on ΔFOSB binding in dorsal hippocampus *in vivo*. APP mice were infused via osmotic minipumps into the dorsal hippocampus for 3 days with vehicle (‘Control’) or JPC0661 (‘Treatment’). Subsequently, genome-wide ΔFOSB binding was quantified with CUT&RUN-sequencing performed on nuclei isolated from unilateral dorsal hippocampus tissue of individual vehicle- or JPC0661-treated mice. **A)** Total number of ΔFOSB-bound peaks in the dorsal hippocampus of male and female APP mice receiving vehicle vs. JPC0661. **B)** Width of ΔFOSB-bound peaks from samples in **A)**. Data for **A)** and **B)** below are shown as mean ± SEM (n = 5 per group); *p < 0.05. No sex differences were observed for **A)** and **B)**. **C)** Genomic distribution of ΔFOSB peaks in the dorsal hippocampus under vehicle and JPC0661 conditions, showing the percentage of peaks at the promoter, intronic, or intergenic regions. **D)** Heatmaps showing ΔFOSB binding intensity centered around the transcription start sites (TSSs) in APP mice under vehicle (left) and JPC0661 (right) conditions, showing reduced number and amplitudes in the latter condition. **E)** Motif analysis showing that the vast majority of ΔFOSB peaks are bound to sequences that contain AP1 consensus sequences (TGA C/G TCA). **F)** Number of unique genes with ΔFOSB peaks bound to gene promoter regions in APP mice treated with vehicle or JPC0661, analyzed via Venn diagrams, indicating that the predominant effect of JPC0661 is to deplete peaks seen also under vehicle-treated conditions. **G)** Gene Ontology (GO) enrichment analysis of genes associated with ΔFOSB promoter-binding peaks in vehicle-treated (Control_Promoter, red) and JPC0661-treated (JPC0661_Promoter, green) conditions. Dot size represents the significance (-log_10_ of the adjusted p-value), and “Odds.Ratio” indicates pathway enrichment. **H)** Example of ΔFOSB binding at the promoter region of *Creb1*. Shown are samples from the dorsal hippocampus of APP mice treated with vehicle (top track in sky blue) or treated with JPC0661 (middle track in gold-green) using an anti-ΔFOSB-antibody for CUT&RUN-sequencing vs. using an IgG-antibody as a control for non-specific binding (bottom track in dark blue; subtracted background). **I)** Number of unique genes with ΔFOSB peaks bound to the distal intergenic regions in APP mice treated with vehicle or JPC0661, analyzed via Venn diagrams, indicating that the predominant effect of JPC0661 is to deplete peaks seen under vehicle-treated conditions. **J)** GO terms for the unique genes bound by ΔFOSB at intergenic regions, as described also in **G)**. **K)** Example of ΔFOSB binding at a distal intergenic/putative enhancer region of *Fos*, as described in **H)**.

We next analyzed the genomic distribution of ΔFOSB peaks across annotated features (**Fig. 3C**). Binding was broadly distributed across promoters, introns, and distal intergenic regions in both groups, with a subtle shift following JPC0661 treatment: a modest decline in promoter-proximal peaks (≤1 kb) and a relative enrichment in distal intergenic peaks (**Fig. 3D**). We performed motif enrichment analysis on the ΔFOSB-bound sequences and confirmed that most binding peaks were associated with an AP-1 consensus sequence (TGA C/G TCA), which provided important validation, specificity, and quality of these CUT&RUN data. Indeed, the top 10 ranked binding motifs identified in ΔFOSB-bound peaks (using samples from vehicle-treated mice for CUT&RUN sequencing) corresponded to consensus AP-1 binding sites (**Fig. 3E**).

We then analyzed the unique genes bound by ΔFOSB by comparing the ΔFOSB binding peaks detected under vehicle and JPC0661 conditions, respectively, in Venn diagrams assessing promoter regions (**Fig. 3F**) and intergenic regions (**Fig. 3I**). ΔFOSB was bound to a total of 7,472 unique genes in vehicle-treated animals, but this was reduced to 2,975 genes after JPC0661 treatment, representing an approximately 60% reduction (**Fig. 3F**). Most of the ΔFOSB-binding peaks that remained stably present in JPC0661-treated mice (about 90%) were found in vehicle-treated mice as well, indicating that the compound induced an overall global loss of DNA-binding *in vivo* but not an altered pattern of binding to novel DNA sites. Gene ontology (GO) analysis showed that the 4,804 genes depleted of ΔFOSB binding after JPC0661 treatment were associated with proteasome-mediated degradation, while the retained genes were enriched in circadian and neuronal pathways (**Fig. 3G**). Overall, these findings indicate that JPC0661 treatment might preferentially reduce ΔFOSB binding to certain families of genes *in vivo*, although this possibility requires further study. Investigation of individual gene promoters, e.g., the *Creb1* promoter, provided clear evidence of decreased ΔFOSB binding caused by JPC0661 treatment (**Fig. 3H**).

We then extended this analysis to distal intergenic-associated peaks, identifying the nearest genes associated with ΔFOSB binding. Control animals exhibited ΔFOSB binding at 3,496 unique intergenic loci, which decreased to 1,833 following treatment (about 48% reduction; **Fig. 3I**). Again, the vast majority of the 2,197 unique genes assigned to ΔFOSB binding peaks that remained stable after JPC0661 treatment were also detected in vehicle-treated mice (1,833 genes), i.e., 83% (**Fig. 3I**). GO analysis of intergenic region-associated genes revealed enrichment for proteasome-related degradation and neuronal development pathways in vehicle-treated mice. By contrast, genes retaining ΔFOSB binding after treatment were enriched for alternative splicing, synaptic transmission, and circadian clock entrainment (**Fig. 3J**). The 1,663 unique genes whose associated intergenic regions were seemingly depleted of ΔFOSB binding after JPC0661 treatment were enriched in pathways related to proteasome-mediated degradation, modification-dependent protein catabolic processes, and regulation of proteasomal ubiquitin-dependent protein catabolic processes (**Fig. 3J**). The *Fos* locus provided an example of a JPC0661-induced decrease in ΔFOSB binding at a putative intergenic enhancer region (12.5 kb upstream) associated with this gene (**Fig. 3K**). Genome browser tracks of several other previously established *in-vivo* target genes for ΔFOSB, e.g., *Grin2a, Sirt2, H2az1, Grin2b*, and *Homer1*, demonstrated similar reductions in ΔFOSB occupancy in response to JPC0661 treatment (**Supplementary Fig. S2B**).

Together, these data indicate that *in vivo* JPC0661 administration causes a robust, broad reduction in ΔFOSB binding across most this transcription factor’s normal genomic locations, consistent with our several lines of evidence that the compound decreases ΔFOSB binding to AP-1 DNA sequences, as well as inhibition of ΔFOSB function.

### 3D Structure of ΔFOSB/JUND bZIP+JPC0661_comm_

To probe how JPC0661 could be inhibiting the binding of ΔFOSB/JUND to DNA, we determined the 3D X-ray crystal structure of the DNA-binding region of ΔFOSB/JUND, the bZIP domain, in complex with JPC0661comm to a resolution of 1.73 Å (Fig. 4; Table 1). We used the bZIP domain of ΔFOSB (E^153^-H^219^), as it contains the DNA-binding region, but lacks the N-terminally disordered region to facilitate crystallization; and its bZIP counterpart in JUND (Q^266^-V^332^; human numbering scheme).

**Figure 4.**
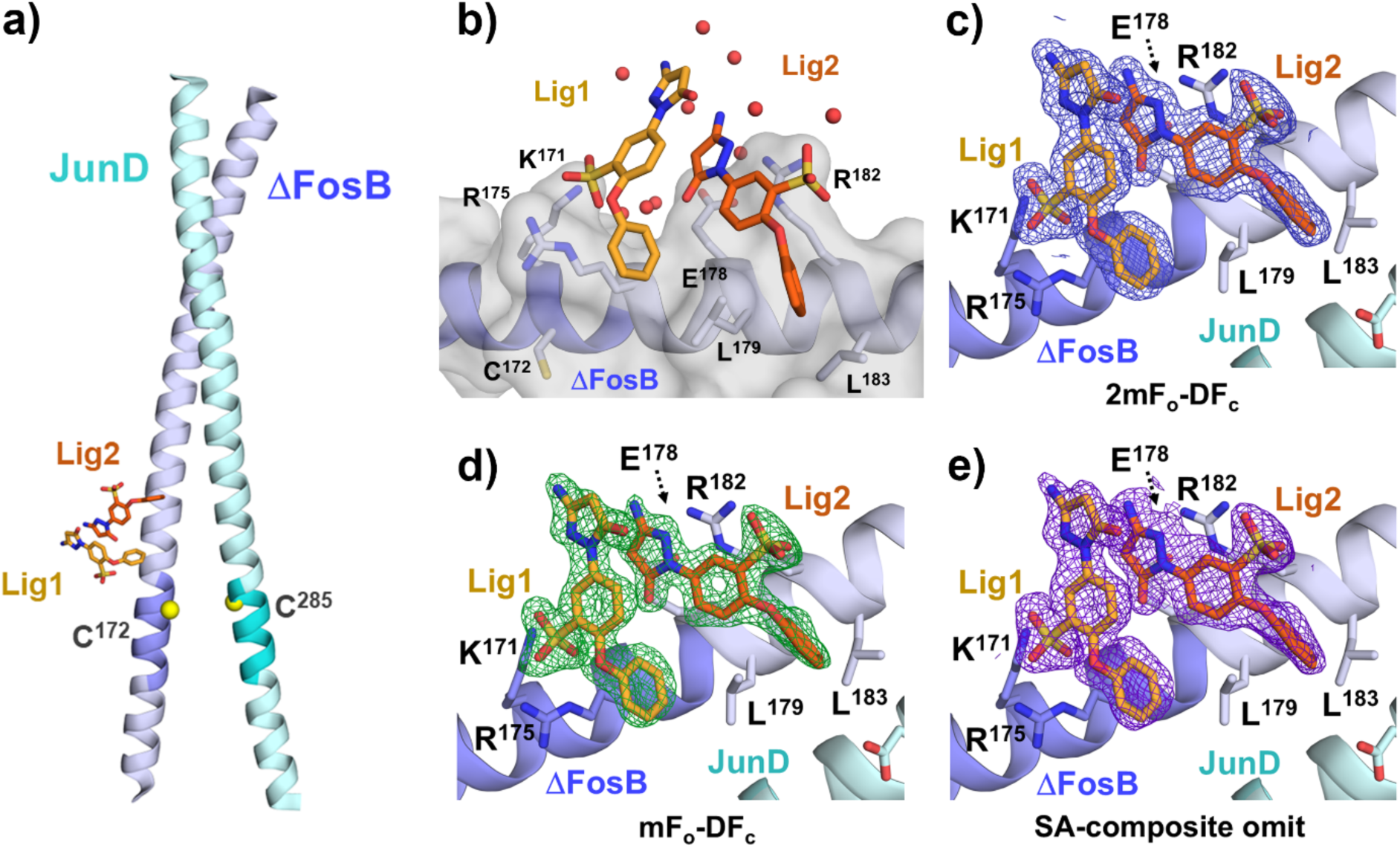
Crystal structure of ΔFOSB/JUND bZIP in complex with JPC0661_comm_. **A)** X-ray crystal structure of the ΔFOSB/JUND bZIP+JPC0661_comm_ showing the location of two bound compound molecules. The redox switch residues ΔFOSB Cys172 and JUND Cys285 are indicated by yellow spheres. ΔFOSB is shown in lilac, and JUND in light cyan; the DNA-binding motifs (ΔFOSB N165-Arg173 and JUND N278-R286) are shown in dark lilac and dark cyan, respectively. JPC0661 molecules (Lig1 and Lig2) are shown with carbon atoms in light and dark orange, respectively, oxygen in red, nitrogen in blue, and sulfur in yellow. **B)** Close-up of the JPC0661-binding site showing the solvent-accessible surface (grey) as well as key residues interacting with the JPC0661 molecules. Color scheme as in **A)**. Water molecules are indicated as red spheres. **C)** 2mF_o_-DF_c_ electron density map surrounding the JPC0661 molecules calculated with phases from the final refined model which contains the JPC0661 molecules (1.3 σ contour level; 2 Å radius map coverage); **D)** mF_o_-DF_c_ difference electron density map surrounding the JPC0661 molecules calculated with phases from a partially refined model lacking the JPC0661 molecules (3 σ contour level; 2 Å radius map coverage); **E)** Simulated-annealing composite omit map surrounding the JPC0661 molecules (1.3 σ contour level; 2 Å radius map coverage).

**Table 1.**
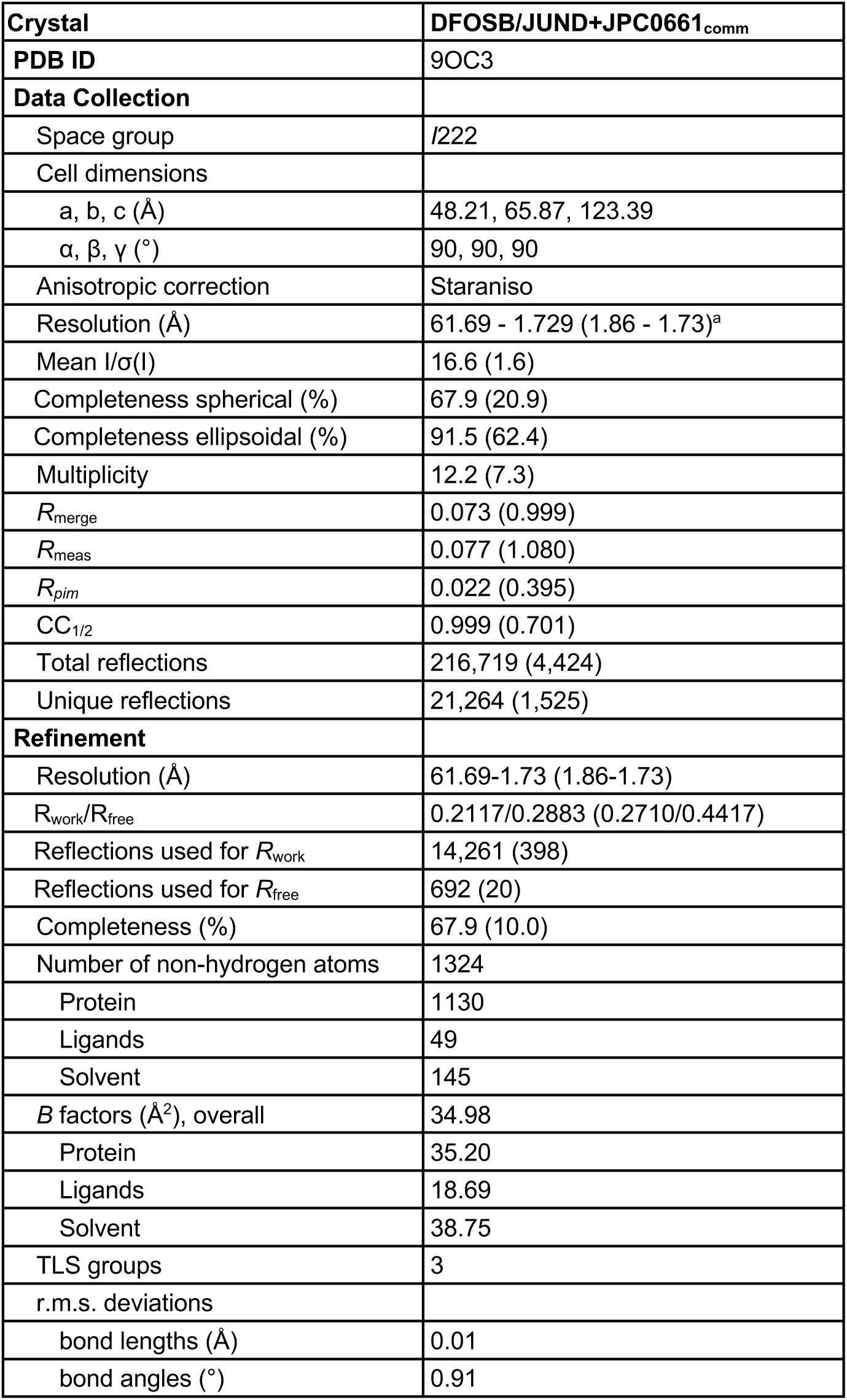
Data collection and structure determination statistics. ^a^Numbers in parentheses are for the highest resolution shell. Data processing statistics are from autoPROC (55) employing anisotropic scaling via STARANISO (56).

Following refinement of the molecular replacement solution, pronounced mFo–DFc difference electron density revealed two JPC0661 molecules (Lig1 and Lig2) bound adjacently within a surface groove of the ΔFOSB bZIP subunit (**Fig. 4A–D**). Subsequent comprehensive refinement, together with a simulated annealing composite omit map, corroborated the well-defined electron density corresponding to both ligands (**Fig. 4E**). Previous work has shown that the DNA-binding site in ΔFOSB/JUND locates to the V-shaped gap formed by the ΔFOSB and JUND bZIP α-helices and in between the putative redox switch residues ΔFOSB Cys172 and JUND Cys285 (13) (PDB ID: 5VPE). The JPC0661-binding site is located outside of the DNA-binding cleft in ΔFOSB; no JPC0661-binding site is found in the corresponding region in JUND. Thus, JPC0661 appears to be an AP1 subunit-selective compound that binds to ΔFOSB.

Extensive interactions with nearby residues hold the two JPC0661 molecules in place (**Fig. 5** and **Supplementary Table S1**). The JPC0661 compound consists of a central phenyl ring with three substituents, a sulfonic acid group, a phenoxy group, and an amino-pyrazolone group (**Fig. 5A**). The two JPC0661 molecules form an extensive network of polar and non-polar interactions with ΔFOSB, including hydrophobic and aromatic stacking interactions, salt bridges, hydrogen bonds and water-mediated interactions (**Fig. 5B**). The two JPC0661 molecules slide side-by-side into a hydrophobic groove on the ΔFOSB bZIP with each phenoxy group buried into the protein and each sulfonic acid group establishing salt bridges located at either end of the groove (**Fig. 5C** and **5D**). The phenoxy group of Lig1 interacts hydrophobically with the side chains of ΔFOSB Arg175, Glu178, and Leu179, and packs against the central phenyl group of Lig2. Its sulfonic acid group forms salt bridges with ΔFOSB Lys171 and Arg175. The central phenyl ring of Lig1 does not contact ΔFOSB but stacks against the amino-pyrazolone group of Lig2 to form π-π interactions while also packing against the edge of Lig2’s central phenyl ring. Similarly, Lig1’s amino-pyrazolone group does not contact ΔFOSB but stacks against Lig2’s amino-pyrazolone ring, forming additional π-π interactions. Lig2’s phenoxy group engages in hydrophobic interactions with ΔFOSB Leu179, Leu183, and Arg182. Its sulfonic acid group forms a salt bridge with ΔFOSB Arg182, which is in turn stabilized by an adjacent salt bridge between ΔFOSB Arg182 and Glu178. Lig2’s central phenyl ring stacks primarily against ΔFOSB Arg182 and fits into a hydrophobic groove formed by ΔFOSB Glu178, Leu179, and Lig1’s phenoxy group. The amino-pyrazolone group of Lig2 stacks against the salt bridge-bonded side chains of ΔFOSB Glu178 and Arg182 on one side, and Lig1’s amino-pyrazolone group on the other, facilitating π-π interactions. Accordingly, Lig1 and Lig2 are packed together by the central phenyl and amino-pyrazolone groups from Lig1 docking against the amino-pyrazolone group of Lig2. Strikingly, a water molecule, WAT411 in chain F (ΔFOSB) in the deposited coordinates (PDB ID: 9OC3), also plays a critical role in mediating the interaction between Lig1 and Lig2 via two close hydrogen bonds between the sulfonic acid group of Lig1 and the amino-pyrazolone ring of Lig2 at 2.7 Å and 2.5 Å, respectively (**Fig. 5D**). Thus, the two JPC0661 molecules fill a hydrophobic groove on the surface of the ΔFOSB bZIP that is located outside of the DNA-binding site.

**Figure 5.**
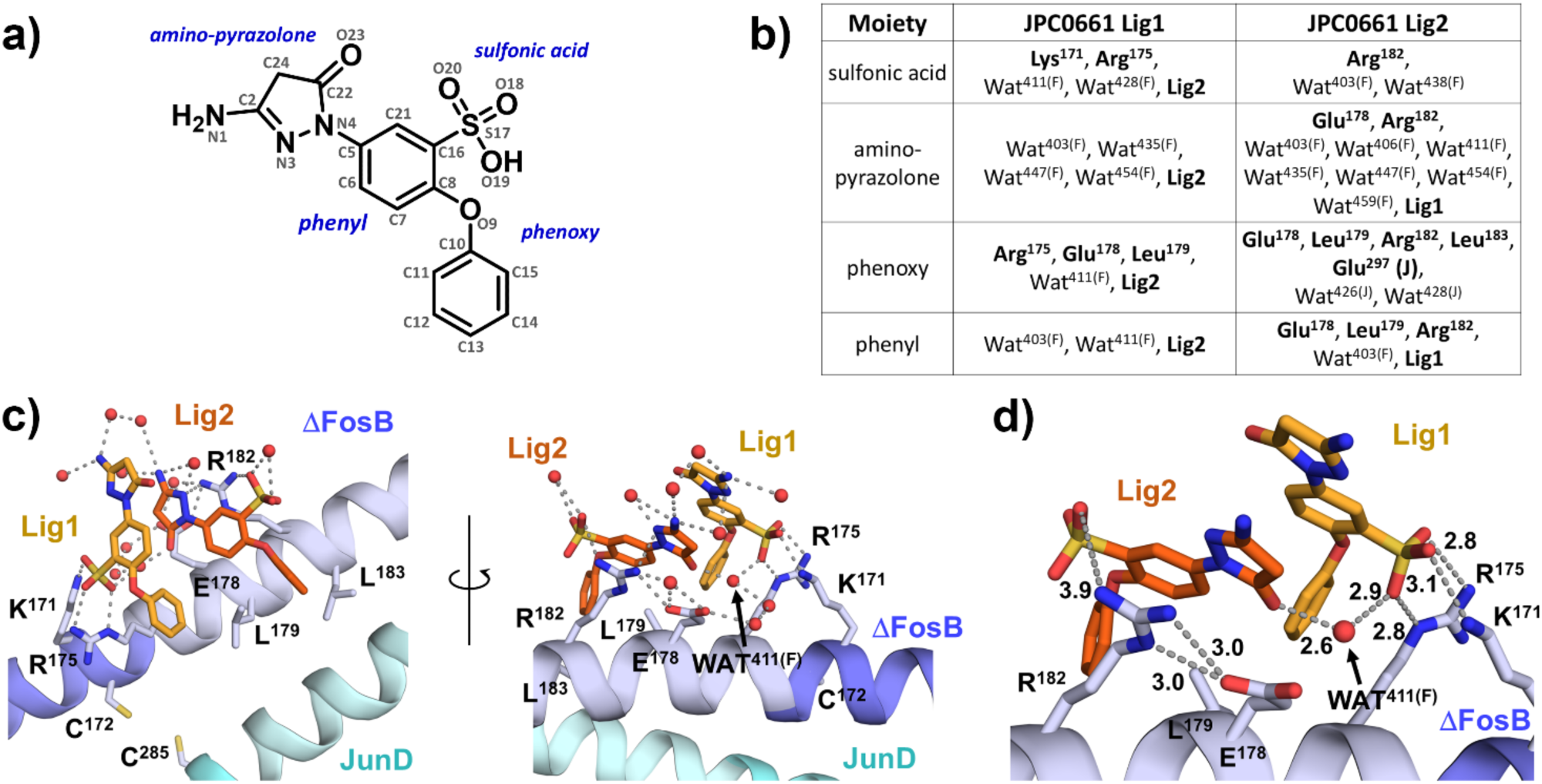
Binding mode of JPC0661 to the ΔFOSB/JUND bZIP. **A)** Structure of JPC0661 with chemical substituents indicated. **B)** Interactions between the ΔFOSB/JUND bZIP and JPC0661 Lig1 and Lig2 within 5 Å. Water numbers refer to those in the deposited coordinates (PDB ID: 9OC3; ΔFOSB has chain ID ‘F’, and JUND has chain ID ‘J’). **C)** Front and back views of the JPC0661-binding site, including key water molecules (red spheres) and their polar interactions indicated by dashed lines. **D)** Zoom-in of the JPC0661-binding site displaying key interactions as discussed in the text. In **C)**, and **D)**, the same color scheme as in Fig. 4A was used. WAT411 (F) refers to water molecule 411 in chain F (ΔFOSB) in the deposited coordinates (PDB ID: 9OC3).

### Characterization of JPC0661 Binding to ΔFOSB

To further validate JPC0661 as an inhibitor of ΔFOSB function and probe the chemical moieties important for compound action, we synthesized a panel of analogs with different chemical substructures and compared these to JPC0661 (**Fig. 6A**). We tested these compounds in FP-DRC assays against ΔFOSB/JUND heterodimers and ΔFOSB homomers and in our cell-based AP1-luciferase reporter assay (**Fig. 6B** and **Supplementary Fig. S1**). YL0324 was designed to test whether the sulfonic acid group is required for compound activity. This compound was poorly active, both in our biochemical and cell-based reporter assays, demonstrating the crucial requirement of the sulfonic acid moiety for compound activity and the specific nature of JPC0661 action (**Fig. 6B** and **Supplementary Fig. S1)**. Three other compounds, YL0325, YL0327, and YL0328, were designed to probe the role of the amino-pyrazolone ring. YL0325, YL0327, and YL0328 were still able to disrupt the binding of ΔFOSB/JUND or ΔFOSB to the AP1 oligonucleotide in the FP-assays (IC_50_ 10-90 μM against ΔFOSB/JUND) (**Fig. 6B, Supplementary Fig. S1A,** and **Supplementary Fig. S1B**). Importantly, in cell-based reporter assays, these compounds inhibited AP1-driven luciferase activity very efficiently, with the most active analog, YL0327, demonstrating an IC_50_ that is reduced by a factor of 20 compared to JPC0661 (IC_50_ 0.05 μM) (**Fig. 6B** and **Supplementary Fig. S1C**).

**Figure 6.**
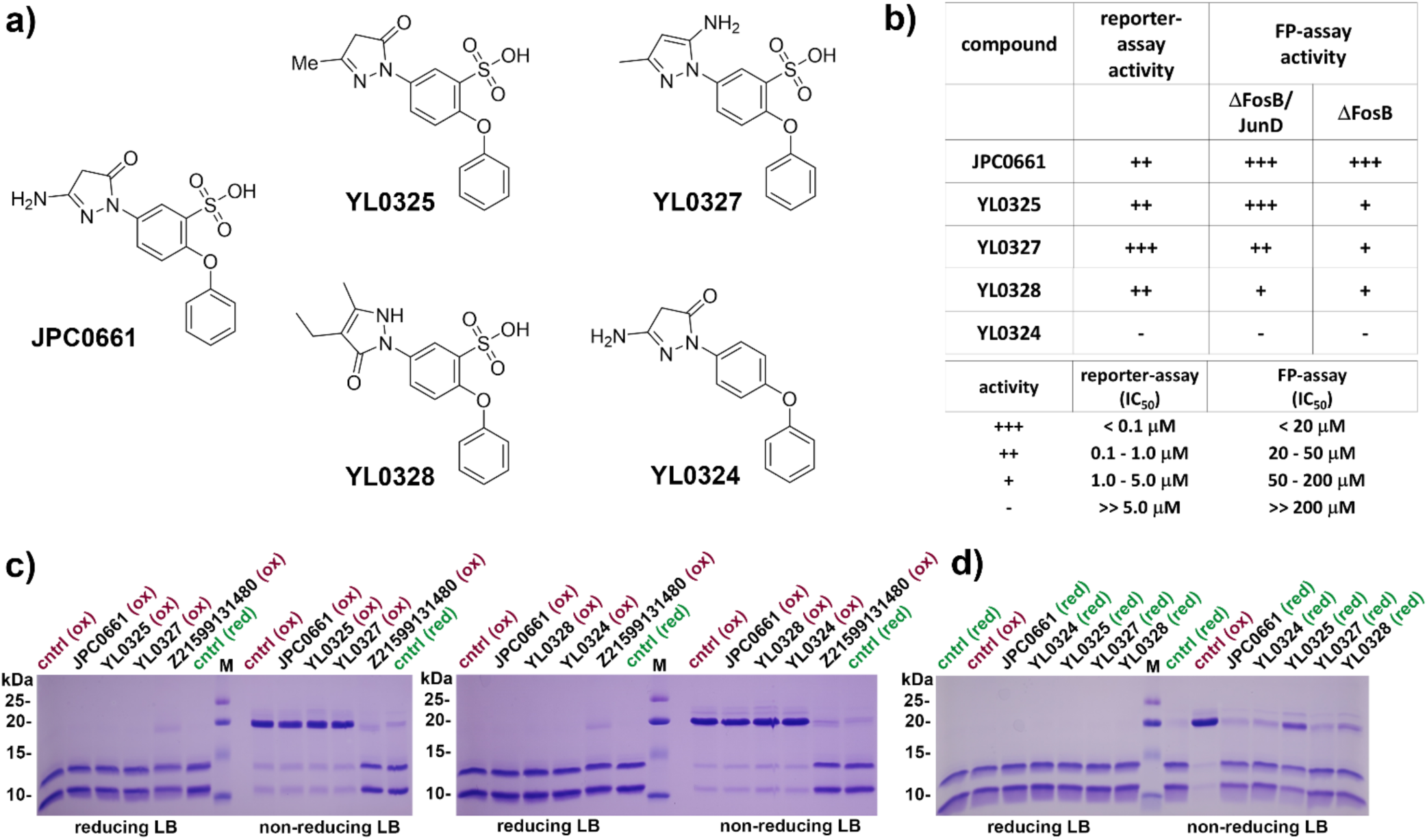
Validation of JPC0661 as an inhibitor of ΔFOSB. **A)** A panel of JPC0661 analogs designed to probe the role of the sulfonic acid group and the amino-pyrazolone group. **B)** Table summarizing the JPC0661 analogs tested in AP1-reporter assays using AP1-luc HEK293 cells, yielding cell-based IC50 values and FP-DRC assays. Lower IC_50_ values are marked with more plus signs (+) and indicate higher activity. The plots from the FP-DRC and cell-based DRC assays for these compounds are shown in **Supplementary Fig. S1**. **C)** ΔFOSB/JUND bZIP incubated with compounds (0.5 mM) with 100 μM diamide (‘ox’, oxidized) or without diamide (‘red’, reduced) and assessed by SDS-PAGE (with or without reducing agent in the loading buffer). **D)** ΔFOSB/JUND bZIP protein incubated with compounds (0.5 mM) with no diamide (‘red) or protein alone (no compound) incubated with diamide (‘ox’) as a control and then assessed by SDS-PAGE (with or without reducing agent in the loading buffer). In **C)** and **D)** ‘cntrl’ denotes the ΔFOSB/JUND bZIP protein in absence of compound.

Next, to assess whether the compounds inhibited the binding of ΔFOSB/JUND heterodimers to DNA by limiting the conformational flexibility of the helical ΔFOSB DNA-binding motif and its ability to swivel and insert into the major groove of DNA (discussed in **Fig. 7A**), we used oxidation-induced closure of the redox switch (ΔFOSB Cys172 and JUND Cys285) as a proxy. When the redox switch closes, the ΔFOSB bZIP helix kinks at a hinge (ΔFOSB Arg 176-Arg 177), enabling ΔFOSB Cys172 to approach JUND Cys285 and form a disulfide bond (19). In presence of the oxidizing agent diamide (100 μM), none of the compounds could prevent the closure of the redox switch when added in excess (0.5 mM) and the oxidized ΔFOSB/JUND bZIP from migrating as a covalently attached heterodimer when assessed by semi-native SDS-PAGE (in the absence of reducing agent) (**Fig. 6C**). Furthermore, under these experimental conditions, the covalent modifier Z21599131480, which attaches to ΔFOSB Cys172 (19) blocked redox switch closure while the panel of compounds did not, suggesting that these compounds also do not covalently modify ΔFOSB Cys172 (**Fig. 6C**).

**Figure 7.**
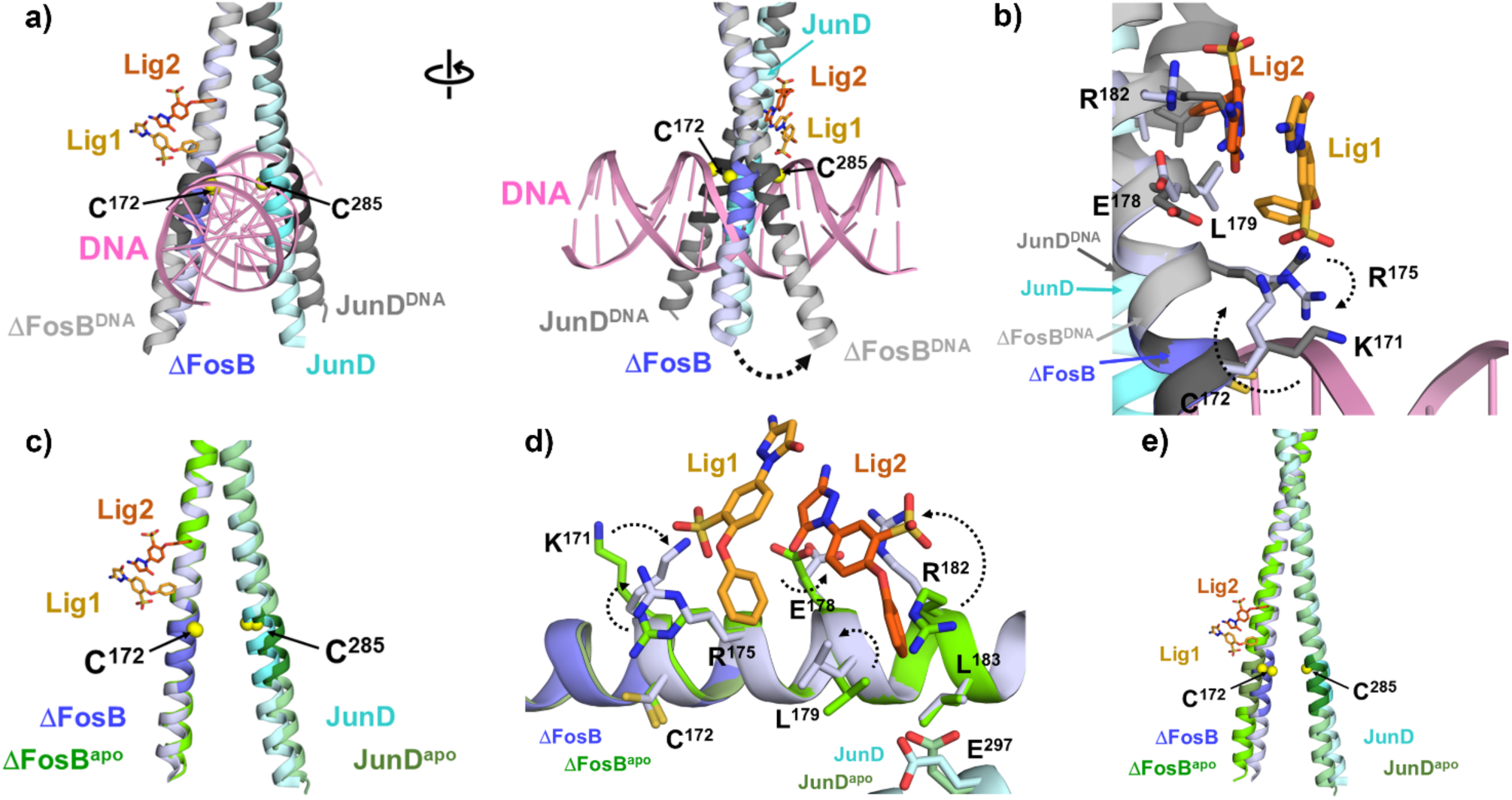
Molecular mechanism of JPC0661 action. **A)** Comparison of the ΔFOSB/JUND bZIP in complex with JPC0661_comm_ (this study) and an AP1-site containing oligonucleotide (PDB ID: 5VPE). The main-chain atoms of ΔFOSB Glu178-Gln189 were used for the superposition. ΔFOSB/JUND bZIP+DNA is shown in grey, with the DNA-binding motifs in darker grey. The redox switch residues ΔFOSB Cys172 and JUND Cys285 are indicated by yellow spheres. **B)** Zoom-in of the superposition shown in **A)**, comparing the binding sites for JPC0661 and DNA. Key residues are displayed, and dashed lines indicate major structural differences. **C)** Comparison of the 3D structure of ΔFOSB/JUND bZIP in complex with JPC0661_comm_ (this study) and the unliganded form of ΔFOSB/JUND bZIP in its reduced state (PDB ID: 7UCC; green color scheme). The main chain atoms of ΔFOSB Glu178-Gln189 were used for the superposition. **D)** Zoom-in of the superposition shown in **C)**, comparing the binding site for JPC0661 and this region in the apo-form of the ΔFOSB/JUND bZIP. Dashed lines indicate structural rearrangements in key side chains as a result of JPC0661-binding, suggesting an induced fit. **E)** Comparison of the 3D structure of ΔFOSB/JUND bZIP in complex with JPC0661_comm_ (this study) and in the unliganded form of ΔFOSB/JUND bZIP in its reduced state (PDB ID: 7UCC; green color scheme). The leucine zipper residues JUND Lys301-Leu317 were used for the superposition to highlight differences in the curvature of the helices. In **A)**–**E)**, the color scheme for ΔFOSB/JUND bZIP+JPC0661_comm_ is as described in Fig. 4A.

We next tested whether our panel of compounds could conversely promote redox-switch closure by oxidizing ΔFOSB/JUND heterodimers, thereby inhibiting DNA-binding. To this end, we assessed the ability of these compounds to directly oxidize ΔFOSB/JUND bZIP heterodimers, again under semi-native SDS-PAGE conditions (**Fig. 6d**). The compounds JPC0661, YL0327 and YL0328 did not trigger oxidation-induced closure of the redox switch in the ΔFOSB/JUND bZIP, while YL0325 was mildly oxidizing. YL0324 also did not impact the protein as would be expected for an inactive compound. Furthermore, these results suggest that TCEP does not diminish the inhibitory action of our compounds by reversing their oxidizing effects; instead, it appears to act through a different mechanism, for example, by binding directly to ΔFOSB and blocking the groove in that accommodates the compounds.

From our analogs of JPC0661, we conclude that the sulfonic acid moiety in JPC0661 is essential for compound activity, while the amino-pyrazolone ring can be modified to yield compounds with strong activity in cell-based assays. Furthermore, the compounds inhibit ΔFOSB/JUND binding to DNA using a mechanism that does not involve oxidizing the protein.

### Mechanism of Action of JPC0661

Several possible mechanisms can be envisioned to explain how JPC0661 disrupts the complex between ΔFOSB/JUND and DNA. Importantly, JPC0661 does not compete with DNA directly because its binding site is located outside of the V-shaped DNA interaction cleft at the fulcrum of the bZIP forceps. To assess how residues that comprise the JPC0661-interaction site in ΔFOSB could impact DNA binding, we compared the crystal structures of ΔFOSB/JUND bZIP bound to an AP1 oligonucleotide (13) with our structure of ΔFOSB/JUND bZIP bound to JPC0661_comm_ in the region ΔFOSB Ala168-Gln184 and JUND Ala281-Thr298 (**Fig. 7A**). In these two structures, the bZIP forceps harboring the DNA-binding motifs are splayed apart to different extents, with helices from the DNA-bound form scissored more widely open to allow insertion into the major groove of DNA. In the DNA-bound form, ΔFOSB Lys171 and ΔFOSB Arg175 interact solely with the DNA backbone (not with nucleotide bases) (13). By contrast, in the JPC0661-bound form, ΔFOSB Lys171 and ΔFOSB Arg175 interact with the sulfonic acid group of Lig1 (**Fig. 5C** and **5D**). This suggests that JPC0661 induces a rearrangement of ΔFOSB Lys171 and Arg175 upon binding, which could weaken the interaction of the ΔFOSB/JUND bZIP with DNA (**Fig. 7B**). Alternatively, interaction of Lig2 with ΔFOSB Arg182 could be key because this residue, for example, may normally be needed to stabilize interactions with JUND or at least this region near the bZIP dimer interface (**Fig. 7D**).

To assess if the presence of JPC0661 causes a change in the conformation of the ΔFOSB bZIP helix, which might impact its DNA-binding abilities or rearrange its side chains, we compared the crystal structure of ΔFOSB/JUND bZIP+JPC0661_comm_ with the unliganded ΔFOSB/JUND bZIP in the reduced state (19) (**Fig. 7C**). Near the binding groove, the overall architecture and curvature of the ΔFOSB bZIP helix are very similar in the compound-bound and unliganded forms (superimposing ΔFOSB Glu178-Gln189). By contrast, key ΔFOSB residues that form the compound-binding groove (Lys171, Arg175, Glu178, Leu179, and Arg182) adopt dramatically different conformations when JPC0661 is bound (**Fig. 7D**). Thus, this novel compound-binding groove appears malleable, suggesting that JPC0661 induces side-chain rearrangements to optimize its interactions with ΔFOSB via an ‘induced fit’.

To further explore if JPC0661 binding prevents flexibility in the ΔFOSB bZIP α-helix, a requirement to insert into the major groove when binding to DNA (**Fig. 7A**), we again compared the structure of ΔFOSB/JUND bZIP+JPC0661_comm_ with that of compound-free ΔFOSB/JUND bZIP in the reduced form. For this comparison, we superimposed the central leucine zipper residues from JUND (Lys301-Leu317) to accentuate any differences in the curvature of both helices in the ΔFOSB/JUND bZIP (**Fig. 7E**). Though the forceps of the ΔFOSB/JUND bZIP splay apart more when compound is present, this difference is relatively small and similar to the variability seen for ΔFOSB and JUND bZIP helices between various crystal forms (13, 14, 19). These structural assessments agree with our in-solution studies demonstrating that JPC0661 does not alter the ability of diamide to induce disulfide bond formation between ΔFOSB Cys172 and JUND Cys285 (closing of the redox switch) nor does it oxidize the protein itself (**Fig. 6C** and **6D**). Our results suggest that JPC0661 disrupts the complex between ΔFOSB/JUND and DNA by binding to a groove outside of the DNA-binding cleft and works as an allosteric effector to decrease DNA binding of ΔFOSB/JUND by competing for specific residues of ΔFOSB that otherwise play a role in regulating DNA-binding.

### Utility of the Druggable Groove to Target Other bZIP Transcription Factors

To assess the broader conservation and potential utility of this novel groove for targeting other members of the bZIP transcription factor family that play key roles in neuroplasticity, we mapped the residues that form the JPC0661-binding site (‘ΔFOSB/JUND+cmpd’) onto the crystal structures of the ΔFOSB/JUND bZIP in different states (**Fig. 8A**). In the ligand-free, reduced form of the protein (‘ΔFOSB/JUND’), the compound-binding groove is not fully formed. This reflects JPC0661 binding via an induced fit that triggers ΔFOSB Leu179 to adopt a different rotamer, generating a deeper, hydrophobic groove that accommodates Lig1 and Lig2. Similarly, ΔFOSB Lys171 and ΔFOSB Arg182 adopt different rotamers to engage Lig1 and Lig2 fully (refer to **Fig. 7D**). In the DNA-bound state (‘ΔFOSB/JUND+DNA’), the side chains of ΔFOSB Lys171 and ΔFOSB Arg175 rearrange to interact with the phosphate backbone, thereby filling a large part of the compound-binding groove. In the oxidized form of ΔFOSB/JUND (with the redox switch “closed”), the compound binding site would not be expected to exist because the ΔFOSB helix unravels and kinks to enable ΔFOSB Cys172 and JUND Cys285 to covalently bind (not shown (13, 19)). Also, the apo-form of the ΔFOSB homomer (‘ΔFOSB/ΔFOSB’) does not show a preformed binding site with a deep cleft near Leu179, suggesting that JPC0661 also must rearrange residues in the homomeric form to form a binding site, just like in ΔFOSB/JUND heterodimers.

**Figure 8.**
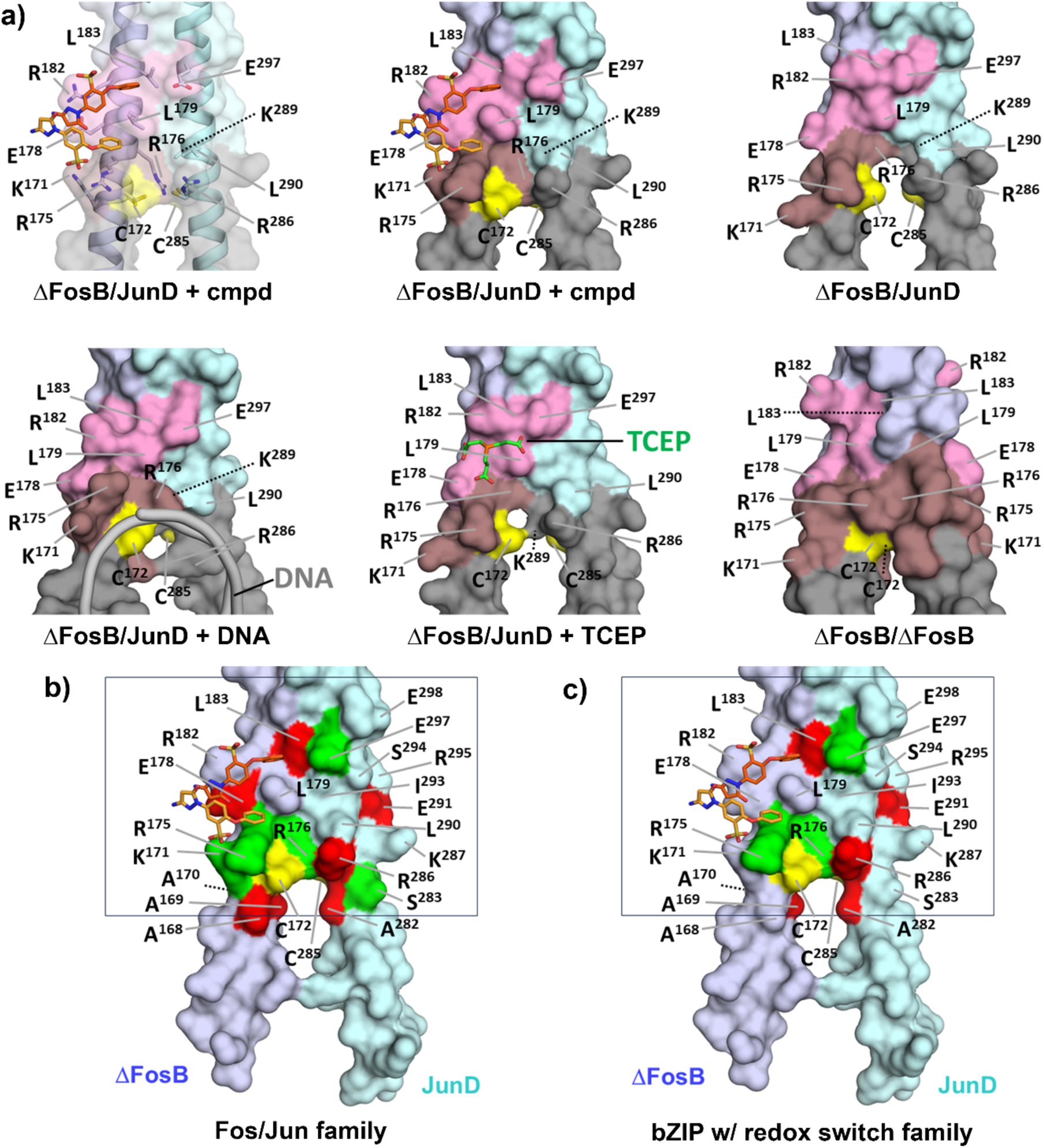
Targeting ΔFOSB. **A)** Comparison of the JPC0661-binding region in different states of the ΔFOSB bZIP for which 3D structures have been determined. ΔFOSB/JUND+cmpd (this study), ΔFOSB/JUND (PDB ID: 7UCC), ΔFOSB/JUND+DNA (PDB ID: 5VPE), ΔFOSB/JUND+TCEP (this study), and ΔFOSB/ΔFOSB (PDB ID: 6UCI). For ΔFOSB/JUND+cmpd, two representations are shown, one with a transparent surface revealing key side chains, and one with a solid surface highlighting the shape of the groove. The protein solvent-accessible surfaces are shown using the following color scheme: residues in ΔFOSB (Glu178, Leu179, Arg182, Leu183) and JUND (Glu297) which are part of the JPC0661-binding region but not part of the DNA-binding site are shown in pink; residues in ΔFOSB (Lys171, Arg175, Arg176) and JUND which are part of the JPC0661-binding region and also the DNA-binding site are shown in brown; residues in ΔFOSB and JUND (e.g., Arg286) which are part of the DNA-binding site are shown in grey; residues in ΔFOSB and JUND which are not part of the JPC0661-binding region or the DNA-binding site are shown in lilac (ΔFOSB) or cyan (JUND; e.g., Lys289, Leu290); the cysteine residues ΔFOSB Cys172 and JUND Cys285 are shown in yellow; Lig1 and Lig2 are shown with orange carbon atoms and TCEP with green carbon atoms. **B)** Sequence conservation of human FOS and JUN family members mapped onto the molecular surface of the ΔFOSB/JUND+cmpd (this study) in the compound-binding area (rectangle). Color scheme: red, conserved; green, semi-conserved; yellow, redox switch. The region outside the binding region is shown in lilac (ΔFOSB) or cyan (JUND). **C)** Sequence conservation of 29 human bZIP-containing proteins mapped onto the molecular surface of the ΔFOSB/JUND+cmpd (this study) in the compound-binding area. In **B)** and **C),** the semi-conserved residues were defined as the following groups: (Arg, Lys), (Gln, Glu), (Ala, Ser, Thr), and (Val, Leu).

We next mapped the amino acid sequences of different AP1 transcription factor family members onto the compound-binding groove (**Fig. 8B** and **8C**). Important differences are seen between human FOS and JUN family members (eight sequences) (**Fig. 8B** and **Supplementary Fig. S3**). Notably, the critical residues ΔFOSB Leu179 and ΔFOSB Arg182, which shape the binding groove and provide key hydrophobic interactions and/or engage the sulfonic acid group of Lig2, are not conserved in human FOS/JUN. Leu179, which is present in all human FOS proteins (FOSB, ΔFOSB, FOS, FOSL1, and FOSL2) is the much larger, positively charged arginine residue in JUN family members (JUN, JUNB, JUND, and JDP2) (**Supplementary Fig. S3**). Arg182 is present in some human FOS and JUN proteins (FOSB, ΔFOSB, JUNB, JUN, and JUND) but not in others where this position is highly divergent (Thr in FOS, Phe in FOSL1 and JDP2, and Lys in FOSL2) (**Supplementary Fig. S3**). JPC0661 does not appear to bind to JUND in our crystal structure, though we cannot rule out interference by the crystal packing. Comparing a group of human bZIP transcription factor family members more broadly, including ATFs, CREBs, FOS, JUN, and MAFs (29 sequences), revealed very low sequence identity in this compound-binding groove (**Fig. 8C** and **Supplementary Fig. S4**). The most conserved residue is Arg175, which is found either as an arginine or lysine residue, preserving the positive charge on the side chain. AP1 transcription factors may thus share a common regulatory mechanism outside the DNA-binding site that triggers DNA release when this positively charged residue is lured away from the DNA backbone. Beyond Leu183 (a leucine-zipper residue whose side chain points inwards, away from the protein surface), no other residue in the compound-binding groove is conserved among the 29 TFs (**Fig. 8C** and **Supplementary Fig. S4**). Residues at key positions carry very different side chains, e.g., Lys171 (E, R, K, S, N, A), Glu178 (E, V, L, D, V, C, T, Q), Leu179 (L, Y, W, K, C, R, Q, T, N, I), and Arg182 (R, C, S, N, Y, T, F, K, V, I, H, E, Q, G) (**Fig.** 8C **and Supplementary Fig. S4).**

Taken together, our results suggest that AP1 transcription factors contain a region outside of the DNA-binding site, which can be manipulated with pharmacological agents to prevent DNA binding or promote DNA release. Importantly, this druggable surface displays key amino acid differences between family members, suggesting that compounds can be designed to target both closely related as well as more distantly related human AP1 transcription factors.

## Discussion

Here, we demonstrate that the binding of the AP1 transcription factor ΔFOSB to DNA can be regulated *in vitro* and in the brain *in vivo* through a novel groove that is located between the DNA-binding cleft and the leucine zipper dimerization interface, and that it can be targeted with small molecules. Compound binding to just the single ΔFOSB subunit of the multimeric protein complex is sufficient to decrease the amount of AP1 transcription factor bound to DNA by about 60% *in vivo*. This groove displays key differences between ΔFOSB and FOS, and it has very limited sequence conservation among 29 bZIP-domain-containing transcription factors, suggesting that it can be exploited to engineer AP1-subunit selective compounds. Accordingly, our studies thus reveal a novel molecular mechanism to modulate AP1-transcription-factor binding to and/or release from DNA, reveal a potential hotspot for endogenous cofactors of ΔFOSB, and hold promise that an entirely novel class of small molecules can be leveraged to target AP1 family transcription factors under normal and diseased states.

### Are bZIP Transcription Factors Undruggable?

AP1 transcription factors have been considered challenging drug targets (21), (22). Compounds targeting them must be highly subunit-selective, because this family is composed of many different members that each encode a subunit with unique biological functions that are determined by the cellular environment and dimerization partners, e.g., the same transcription factor can exert oncogenic or anti-oncogenic effects depending on the context (22–24). Strategies to target AP1 transcription factors directly have focused on their bZIP domains, for example, by: 1) blocking the DNA-binding cleft; 2) interfering with dimerization of the leucine zipper; 3) blocking the putative redox switch that controls the conformation of the DNA-binding site; or 4) blocking protein:cofactor interactions (19, 23). The DNA-binding cleft, which is buried deep in the fulcrum of the bZIP dimer forceps, is difficult to exploit to develop selective compounds because its surface is highly conserved among the different AP1 subunits (19). Efforts to use peptides to disrupt the dimerization of bZIP leucine zippers have proven challenging, as members of the AP-1 transcription factor family readily form both heterodimers and homodimers. This inherent lack of selectivity complicates the design of peptides capable of selectively disrupting leucine zipper interactions (23, 25). Furthermore, it is generally difficult to optimize cellular and nuclear uptake, bioavailability, and biostability of peptide-based therapeutics (23). The most promising small molecule to date, T5224, which was evaluated in phase-2 clinical trials, targets the FOS/JUN AP1 protein complex directly with an IC_50_ of about 10 μM in cell-based assays; it was designed to block the DNA-binding site, but its actual mechanism and specificity are not known (21–23, 26–28). Other compounds such as SP 100030, SPC-839, mormodin I, nordihydroguaiaretic acid, and a series of arylstibonic acid-containing compounds have also been suggested to interact with the basic DNA-binding region of FOS/JUN, but their mechanisms are also unknown and controversial (21, 29–31).

We previously identified several small molecules that disrupt the binding of ΔFOSB/JUND heterodimers and ΔFOSB homomers to an AP1-site containing oligonucleotide in FP assays in the low to mid-micromolar range (32, 33), but their binding sites and mechanisms of action remain unknown. Likewise, we identified and validated thiol-reactive compounds that covalently modify the redox switch residue ΔFOSB Cys172 and disrupted DNA binding in biochemical studies, but which enhanced the expression of a ΔFOSB-induced AP-1-driven luciferase reporter in cell-based studies through an unknown molecular mechanism of action (19). Thus, despite extensive efforts, targeting AP1 transcription factors has remained a formidable challenge.

### Formulating a Novel Strategy to Target ΔFOSB

To develop compounds that decrease DNA binding (or promote DNA release) of ΔFOSB *in vivo*, we considered the following: most transcription factors are thought to bind and release genomic DNA very rapidly, docking onto DNA for several seconds at most so that target gene transcription occurs in stochastic bursts (34–37). The majority of AP1 transcription factors attach to DNA as pre-formed dimers via their bZIP domains, with the basic region inserting into the major groove of DNA while the leucine zipper residues mediate dimerization (38). FOS, as an exception, may also bind to DNA as a monomer first and assemble into stable dimers on genomic DNA later (39). AP1 consensus sites can be extended by flanking DNA sequences that alter the binding site recognized and/or the 3D shape of the DNA (40, 41). Importantly, a given transcription factor can exhibit a broad range of binding affinities to DNA in cells rather than a single binding affinity (42). Also, the strong binding of a transcription factor to nucleosome-free DNA (to plasmids used in cell-based reporter assays, for instance) does not necessarily translate to strong binding to the same DNA consensus site *in vivo* when it is integrated into chromatin and is part of multi-protein transcriptional complexes (37).

AP1 transcription factors like ΔFOSB and JUND are thought to preferentially bind to entry-exit regions at the edge of nucleosomes; there, the wraps of DNA transition from one nucleosome to the next (37, 43). Nucleosomes likely play an active role in mediating the binding and release of ΔFOSB/JUND heterodimers or ΔFOSB homomers to genomic DNA, controlling the accessibility to DNA-binding sites and/or whether just one protein subunit or both bind to DNA (44). Indeed, at select DNA-binding sites *in vivo* ΔFOSB recruits chromatin remodeling factors such as BRG1, a nucleosome remodeler that promotes nucleosome removal, and CBP/P300, a histone acetyltransferase that acetylates lysines on histones and loosens histone:DNA binding, thereby facilitating ΔFOSB’s ability to bind to DNA and regulate the expression of target genes (18, 45, 46).

Complicating matters even further, AP1 transcription factors like ΔFOSB are thought to form multi-protein transcriptional complexes that incorporate a portfolio of different cofactors and even other transcription factors, which work together to modulate DNA-binding properties and function. As such, ΔFOSB can activate or repress gene expression depending on the cofactors bound (1, 35, 40, 41). As an example, ΔFOSB can recruit HDAC1, which represses target gene expression (6, 7, 47).

Taken together, these considerations suggested to us that AP1-sites bound by ΔFOSB might not all be equally sensitive to a compound targeting ΔFOSB and that gauging mRNA levels from only a select few target genes would likely not be a robust read-out for compound action. Furthermore, because most studies have focused primarily on delineating how AP1 transcription factors bind to DNA, very little is known about how they dissociate from DNA, i.e., how ΔFOSB releases from genomic DNA *in vivo*. Additionally, nucleosomes and co-factors likely promote or prevent binding to and/or the release of AP1 transcription factors like ΔFOSB from AP1 sites in genomic DNA (48), further confounding any straightforward strategy to assess compound action. We thus anticipated that ΔFOSB would likely display a diverse spectrum of DNA-binding and release events, especially given its unusual stability and high levels of protein accumulation, which our compounds would need to target.

These uncertainties and complexities surrounding ΔFOSB’s molecular actions *in vivo,* sketched above, fundamentally shaped our approach to target ΔFOSB with small molecules. We realized that we would not be able to predict the binding sites of ΔFOSB to the genome *in vivo* in advance, nor flag which binding sites would be susceptible to pharmacological agents. We also recognized that we would likely not be able to translate compound activity observed *in vitro* to accurately predict activity *in vivo*, since both our biophysical assays with purified ΔFOSB/JUND protein and short AP1-site-containing oligonucleotides in solution and our cell-based assays in AP1-HEK293 cells lacked the native 3D chromatin constraints and components of large ΔFOSB-containing transcriptional multi-protein complexes bound to the many genomic DNA sites *in vivo* in the brain (most of which fall outside of traditional promotors). We thus opted to screen compound libraries using full-length ΔFOSB/JUND heterodimers and ΔFOSB homomers to increase the chances of finding novel compound-binding sites. We also proceeded to test our compounds in a cell-based reporter assay using an AP1-reporter plasmid despite their modest activity in biochemical FP-DRC assays (IC_50_ of about 0.9 μM for JPC0661). Once activity in cell-based reporter assays was seen, we proceeded to validate compound action in the brain *in vivo,* focusing on a direct biological readout (DNA binding/release) using CUT&RUN-sequencing. In tandem, we validated compound binding and the mechanism of action by determining the crystal structure of the ΔFOSB/JUND+JPC0661_comm_ co-complex and confirmed these results through a series of compound analogs and mutants. This pragmatic and unbiased approach proved to be successful.

### *In-vivo* Administration of JPC0661 Broadly Decreases ΔFOSB Bound to Genomic DNA

We used CUT&RUN-sequencing to map the impact of JPC0661 on the binding of endogenous ΔFOSB to its target genomic regions in the brain *in vivo.* We leveraged previous findings of ours that demonstrated induction of ΔFOSB in the dorsal hippocampus of APP transgenic mice. Our CUT&RUN-sequencing data confirmed that ΔFOSB binds overwhelmingly to AP1 consensus sites in the hippocampus and is distributed across promoter regions, intergenic regions, and gene bodies, as we found previously for the nucleus accumbens (18). Infusion of JPC0661 into the dorsal hippocampus of APP mice caused an approximately 60% reduction in ΔFOSB binding peaks compared to vehicle-infused mice, decreasing sites broadly across all genomic regions, including promoter, intergenic, and intronic regions. A similar reduction was seen in male and female mice. The reduced ΔFOSB binding observed by CUT&RUN-sequencing in the brain following JPC0661 treatment, as well as the actions of JPC0661 as an inhibitor of ΔFOSB in our cell-free and cell-based experimental systems, together validate JPC0661 as an inhibitor of ΔFOSB *in vivo*. It is important to note that we purposefully selected CUT&RUN as our *in vivo* validation method to monitor the impact of JPC0661 on the binding of ΔFOSB to genomic DNA. The CUT&RUN technique uses a ΔFOSB-specific antibody to assess the amount of ΔFOSB bound to genomic DNA in the presence of compound or vehicle (using IgG as a measure for non-specific binding). Thus, we were able to selectively measure the effect of the compound on ΔFOSB regardless of any possible off-target effects of JPC0661. Furthermore, we have previously shown that our ΔFOSB-specific antibody reveals virtually no signal when using brain tissue from *fosb*-null mice in CUT&RUN studies, indicating recognition of ΔFOSB is highly selective (18). It will be interesting in future studies to understand which of the many hundreds of significant ΔFOSB peaks involve ΔFOSB heterodimers with JUND or another AP1 protein or instead represent ΔFOSB homomers. Likewise, it will be important to determine whether specific ΔFOSB peaks can be rendered resistant or particularly susceptible to JPC0661-induced release by co-factors that co-assemble with ΔFOSB into multi-protein transcriptional complexes. Gene ontology analyses of our CUT&RUN-sequencing data indicated that JPC0661 treatment reduces ΔFOSB-binding to genes involved in proteasomal regulatory processes yet leaves binding to genes associated with neuronal development and circadian rhythm preferentially intact. These observations suggest the possibility of compound-induced, selective transcriptional reprogramming, a possibility that now requires further investigation. Future studies are also needed to determine the exact impact of JPC0661-induced reductions in ΔFOSB binding on the downstream transcription of the affected genes and consequent effects on brain function and behavior.

### 2-Phenoxybenzenesulfonic Acid Compounds Target ΔFOSB

Our studies reveal that it is possible to target ΔFOSB *in vivo* pharmacologically, while also revealing mechanistic molecular insights into ΔFOSB action through a panel of compounds (**Fig. 6**). Our crystal structure indicates that the compounds anchor into a hydrophobic surface on ΔFOSB by burying large, bulky hydrophobic substituents. ΔFOSB Arg175, which is engaged by JPC0661, is shown to interact with the phosphate backbone of DNA in ΔFOSB/JUND+DNA crystal structures. ΔFOSB Arg175 is highly conserved and found only as arginine or lysine in 29 human bZIP transcription factors (**Fig. 8** and **Supplementary Fig. S4**). ΔFOSB Lys171 is also engaged by JPC0661 and likewise interacts with the DNA phosphate backbone; it is highly conserved only within the FOS/JUN family members (**Fig. 8** and **Supplementary Fig. S3**). Thus, ΔFOSB Lys171 and ΔFOSB Arg175, are strategically poised to modulate the DNA-binding/release properties of ΔFOSB-containing transcription factors even if they do not interact directly with the DNA bases of the consensus site. Other functionally important arginine residues that only contact the DNA backbone have also been identified in related AP1 transcription factors. For instance, Arg261 in JUN is a critical residue as indicated by the charge-reversal mutation Arg261Asp that prevents DNA binding (though JUN Arg261Ala does not), while Arg261Asp and Arg261Ala each prevent JUN from binding to the co-activator STAT3 (49). ΔFOSB Arg164 and JUND Arg277 are the structural equivalents of JUN Arg261, and they also contact only the DNA backbone, albeit on the opposite, wider side of the V-shaped DNA-binding cleft compared to ΔFOSB Arg175. It is striking that JPC0661 also engages ΔFOSB Arg182.

Our compounds may exploit multiple mechanisms to regulate ΔFOSB. For instance, the two JPC0661 molecules stacked side-by-side may work as a ‘door stop’ to sterically hinder the bZIP forceps from splaying open and the ΔFOSB bZIP helix from bending and binding in the major groove of DNA. Indeed, it is yet unclear whether both JPC0661 Lig1 and Lig2 are needed to inhibit ΔFOSB binding to DNA or promote its release, and studies are ongoing to resolve this issue using a portfolio of biophysical approaches. Interestingly, the compound-binding groove may overlap with protein surfaces that are normally exploited by endogenous cofactors of ΔFOSB, which co-assemble into different multi-protein transcriptional complexes. Indeed, the IC_50_ values of our compounds, as determined by the cell-based luciferase assay, are dramatically lower compared to those from the biochemical FP-DRC assay, which uses purified proteins (e.g., about 100 times lower using YL0325 action on ΔFOSB/JUND). This suggests that the compound-binding groove may be more rigid (lowering the entropic cost) or adopt a (slightly) different conformation in the cellular context compared to purified proteins in solution. In support of this idea, the NFAT1 cofactor binds to a stretch of FOS bZIP that corresponds to this novel ΔFOSB druggable groove, including the JPC0661-binding residues ΔFOSB Glu178, Asp181, and Thr182, based on the crystal structure of the FOS:JUN:NFAT1 complex bound to the DNA site (PDB ID: 1S9K) (**Supplementary Fig. S5**). Another cofactor, CRTC, is proposed to align as a single helix alongside each of the two helical subunits of the CREB bZIP homodimer, generating a four-helical bundle (50), and its binding site also overlaps with the compound-binding groove we have identified. CRTC is a coactivator of CREB, and upon binding, it slows down the dissociation of CREB so that the latter stays bound longer to DNA, thereby increasing CREB-dependent transcription (50). Thus, the druggable groove that we have identified in ΔFOSB may coincide with different cofactor binding sites, offering a potential strategy to regulate the activating vs. repressing effects of ΔFOSB on gene transcription. Key steps moving forward will require further studies to determine the exact mechanism of action of JPC0661, including resolving whether both Lig1 and Lig2 are equally required for compound action, whether other compounds can bind to this groove in ΔFOSB, and the exact mechanism by which these compounds work to alter the binding/release of ΔFOSB to DNA.

### The Potential to Target AP1 Transcription Factors in the Future

Being able to target ΔFOSB with small molecules opens the door to developing pharmaceutical agents to treat ΔFOSB-related diseases. ΔFOSB is an attractive drug target because the protein accumulates to high protein levels in the brain following chronic insults, while under normal conditions it is present at very low levels, suggesting that negative consequences of inhibiting ΔFOSB pharmacologically in a non-pathological context may be limited – a hypothesis which can now be tested *in vivo*. 2-Phenoxybenzenesulfonic acid compounds appear promising leads because of their low molecular weight, limited toxicity in neuronal cells, metabolic stability in mouse liver microsomes, favorable cell-membrane permeability, and activity *in vivo* upon local delivery into the brain. Furthermore, our insights into the structure-activity relationships already indicate a clear mechanism of action and delineate chemical moieties essential for compound activity vs. those that can be replaced, for instance, to develop analogs optimized for bioavailability and/or systemic delivery. A large body of work has demonstrated that genetically inhibiting elevated ΔFOSB in the striatum in response to drugs of abuse reduces their rewarding effects (1, 5) and in response to chronic L-DOPA reduces abnormal involuntary movements (confounding side-effects of L-DOPA treatment in patients diagnosed with PD), while leaving the anti-Parkinsonian action of L-DOPA undiminished (10, 11). In the hippocampus, viral inhibition of accumulated hippocampal ΔFOSB in mouse models of AD can reverse cognitive deficits (6, 7, 51). However, in the above studies, the ΔFOSB function was knocked down by genetic or viral interventions, which are difficult to translate into a therapeutic window and to directly mimic with other modalities (52, 53). AP1 subunit-selective, pharmacological agents tuned more finely to inhibit ΔFOSB, both in the degree of perturbation and duration of perturbation, will be better suited to assess the therapeutic impact of ΔFOSB inhibition and delineate a restorative window. Results of the present study, therefore, provide an important step forward in the generation of an entirely novel class of small-molecule therapeutic agents that selectively target AP1 transcription factors with a defined subunit composition. This work thereby sets the stage for further development of such an approach for a range of clinical applications in which these transcription factors have been implicated.

## Experimental Procedures

### Constructs

The following constructs were used for overexpression in Sf9 insect cells: a) full-length mouse N(His)_6_-ΔFOSB (a splice form of FOSB, UniProt ID: P13346) that contained an N-terminal tag MGHHHHHHAG followed by mouse ΔFOSB residues Phe2-Glu237 in the pfastbac1 vector; and b) full-length mouse N(His)_6_-JUND (UniProt ID: J04509) that contained an N-terminal tag MGHHHHHH followed by mouse JUND residues Glu2-Tyr341 in the pfastbac1 vector.

The following constructs were used for overexpression in *E. coli*: a) mouse/human N(His)_6_-ΔFOSB bZIP (from FOSB, UniProt ID: P13346) containing an N-terminal (His)_6_-tag and a tobacco etch virus (TEV) protease cleavage site (MGHHHHHHENLYFQS) followed by mouse ΔFOSB residues Glu153-His219 in the pET21a-NESG vector; and b) mouse/human N(His)_6_-JUND bZIP (from JUND, UniProt ID: J04509) containing an N-terminal (His)_6_-tag and a TEV protease cleavage site (MGHHHHHHENLYFQS) followed by mouse JUND residues Gln260-Val326 in the pET21a-NESG vector. Note that human and mouse amino acid sequences for the ΔFOSB bZIP and JUND bZIP domains are identical. However, the human JUND bZIP (Gln266-Val332) numbering scheme is shifted by 6 residues compared with the mouse JUND bZIP domain (Gln260-Val326). In this work, the human amino acid sequence numbering is used. All constructs were verified by sequencing.

### Reagents and Biological Resources

#### Chembridge and Maybridge libraries

The Chembridge Library NT638 DIVERSet1 (Chembridge; San Diego, CA), with 30,080 compounds, and Maybridge Library HitFinder V9 (Thermo Fisher, Waltham, MA), with 14,400 compounds, for a total of 44,480 compounds, were plated in 384-well plates with a concentration of 10 mM (in DMSO). Additionally, JFD00458 (JPC0661_comm_) was purchased from Maybridge, Thermo Fisher Scientific, Waltham, Massachusetts, US.

#### Oligonucleotides

The 19-mer cdk5 oligonucleotide (‘cdk5 oligo’ or ‘AP1 oligo’) contains the forward and reverse strands of 5′-CGTCGG<u>TGACTCA</u>AAACAC-3′ (AP1 site underscored) derived from the AP1 site in the cyclin-dependent kinase 5 promoter. The 19-mer scr oligonucleotide (‘scr oligo’) contains the forward and reverse strands of 5′-GTATGCGATACGTCTTTCG-3’ and is composed of the same set of nucleotides as the cdk5 oligo but in a scrambled sequence. The TMR-cdk5 and TMR-scr oligos were generated by annealing equivalent amounts (in moles) of the complementary strands each labeled with TAMRA (Sigma Aldrich) at the 5′-end and heating them to 95 °C for 2.5 min, followed by slow cooling to room temperature (roughly 1 min/°C) and storage at −20 °C as 50 μM stocks in annealing buffer (10 mM Tris pH 8, 50 mM NaCl). For electrophoretic mobility shift assays (EMSAs), the biotinylated forward strand of the cdk5 oligo and the unlabeled reverse strand cdk5 oligo (Sigma Aldrich) were annealed as above, yielding oligonucleotide ‘BIO-cdk5’.

#### Cell lines

Human neural progenitor cells, AP1 luciferase reporter human embryonic kidney 293 recombinant cell line (AP1-luc HEK293): BPS Bioscience, San Diego, CA; catalog number 60405; Neuro2A cell line: ATCC; Manassas, Virginia; catalog number CCL-131.

#### Luciferase reagents

Luciferase assay system: Promega; Madison, WI; catalog number E1501; Passive lysis buffer: Promega; Madison, WI; catalog number E1941; CellTiter-Glo: Promega; Madison, WI; catalog number G7571.

### Purification of Wild-Type ΔFOSB/JUND Heterodimers and ΔFOSB Homomers

Full-length mouse ΔFOSB/JUND heterodimers and ΔFOSB homomers were expressed in Sf9 cells using the Bac-to-Bac system (Invitrogen) as described before (12, 19, 32). Briefly, plaque-purified, amplified baculoviruses were prepared in large quantities (titer ∼1×10^8^ pfu/mL) for infection of large-scale Sf9 insect-cell cultures. For overexpression of the wild-type N(His)_6_-ΔFOSB/N(His)_6_-JUND heterodimer (‘ΔFOSB/JUND’), insect cells at a density of about 1.5×10^6^ cells/mL were coinfected with the appropriate baculoviruses using a multiplicity of infection (MOI) of 1.0-1.5 for N(His)_6_-ΔFOSB and 1.0-3.0 for N(His)_6_-JUND. For overexpression of the N(His)_6_-ΔFOSB homomer (‘ΔFOSB’), insect cells at a density of about 1.5×10^6^ cells/mL were infected with the appropriate baculovirus using an MOI of 1.0-1.5. Typically, 6 L of insect cells in SF900 medium were infected, grown for 72-84 h at 28 °C and 145 rpm, centrifuged (15 min, 3000 rpm at 4 °C), and then the cell pellets were resuspended in PBS, flash-frozen in liquid nitrogen, and stored at −80 °C until further use.

The proteins were purified as follows. Cells were resuspended in lysis buffer (25 mM Tris pH 8.0, 0.2% (v/v) Triton X-100, 1 mM TCEP) using 300 mL per 6 L cell pellet, and protease inhibitors added (0.5 mM PMSF, 1 μg/mL pepstatin, 10 μg/mL leupeptin and two tablets of cOmplete™ EDTA-free Protease Inhibitor Cocktail (Roche)). The mixture was incubated on ice for 30 min and lysed by sonication. Subsequently, 300 mM NaCl, 5 mM MgCl_2_, and 50 μg/mL DNase were added to the lysate, the mixture was incubated for 1 h on ice, and then 0.5 M NaBr and 20 mM imidazole added, followed by centrifugation at 18,000 rpm for 30 minutes at 4 °C. For purification of the N(His)_6_-ΔFOSB homomer, 0.5 M NaBr was added.

For Ni-affinity chromatography, Ni-NTA beads (Thermo Scientific) equilibrated in Ni-Buffer A (20 mM Tris pH 8.0, 0.5 M NaCl; 10 mL beads as a 50% slurry) were added to the supernatant, incubated for 3 h, transferred to an empty column and the protein eluted using a gradient with Ni-Buffer B (20 mM Tris pH 8.0, 0.5 M NaCl, 0.5 M imidazole) The sample was then diluted with 25 mM Tris pH 9, 1 M NaCl to a protein concentration of 0.075 mg/mL, dialyzed to equilibrium against 25 mM Tris pH 9, 300 mM NaCl, 1 mM DTT, 0.5 mM PMSF, and then 3 h against 25 mM Tris pH 9, 75 mM NaCl, 1 mM DTT, 0.5 mM PMSF before anion exchange chromatography. The protein sample was loaded onto a Mono Q 5/50 GL column (Cytiva) equilibrated with Mono Q Buffer A (25 mM Tris pH 9, 75 mM NaCl, 1 mM DTT) and eluted with a gradient (0-100%) using Mono Q Buffer B (25 mM Tris pH 9, 1 M NaCl, 1 mM DTT). The eluted protein was then concentrated and further purified by size exclusion chromatography using a HiLoad 16/600 Superdex 75 pg column (GE Healthcare) equilibrated in buffer (20 mM Tris pH 8.0, 1 M NaCl). For ΔFOSB/JUND, fractions were analyzed by SDS-PAGE during purification to confirm the presence of both proteins. Protein purity was assessed by SDS-PAGE. The purified ΔFOSB/JUND and ΔFOSB proteins were stored as flash-frozen aliquots (3–5 mg/mL in 20 mM Tris pH 8.0, 1 M NaCl).

### Preparation of ΔFOSB/JUND bZIP Domains

ΔFOSB and JUND bZIP domains were expressed in *E. coli* as described previously (13, 14, 19). Typically, 6 L of *E. coli* Rosetta 2 (DE3) cells (Invitrogen) transformed with the respective vectors were grown in LB medium at 37 °C to an OD_600_ of ∼0.5, cooled to 16 °C, and then induced with 0.5 mM IPTG overnight. Cells were centrifuged for 30 min at 4 °C, 4000 rpm, and the pellets were resuspended in 10 mL PBS per 1 L cell pellet, flash-frozen in liquid nitrogen, and stored at −80 °C until further use. For protein purification, thawed cell pellets were resuspended in lysis buffer (20 mM Tris pH 8.0, 250 mM NaCl, 1 mM TCEP, 0.5 mM PMSF), 1.1 mg/mL lysozyme was added, and the mixture was incubated on ice for 30 min and then lysed by sonication. Subsequently, 5 mM MgCl_2_, 1 M NaCl, and 30 μg/mL DNase I were added, and the mixture was further incubated on ice for 30 min. In preparation for Ni-NTA affinity chromatography, 0.5 M NaBr and 20 mM imidazole were added, and the mixture was centrifuged at 18,000 rpm for 30 minutes at 4 °C. For Ni-affinity chromatography, Ni-NTA beads (Thermo Scientific) equilibrated in Ni-Buffer A (20 mM Tris pH 8.0, 1 M NaCl, 500 mM NaBr; 10 mL beads as a 50% slurry) were added to the supernatant, incubated for 3 h, transferred to an empty column, and the protein eluted using a gradient with Ni-Buffer B (20 mM Tris pH 8.0, 1 M NaCl, 500 mM NaBr, 500 mM imidazole). The (His_6_)-tag was then removed by adding TEV protease (using 40 μg protease/mg target protein) to the recombinant protein (in digest buffer containing 1 mM DTT, 50 mM Tris HCl pH 8.0, 0.5 M NaCl, and 1% (v/v) glycerol)) followed by incubation at room temperature for 1 h. A second dose of TEV protease was then added (40 μg protease/mg target protein), and the sample was incubated for an additional 2 h and then centrifuged for 10 min at 4000 rpm. To remove the TEV protease, 1 M NaCl and 0.5 mL Ni-NTA beads (50% slurry) equilibrated in digest buffer were added to the sample. The mixture was incubated at room temperature for 1 h and then centrifuged for 10 min at 900 rpm, and the supernatant containing the cleaved target protein was collected. The supernatant was then passed over a disposable Bio-Spin® column (Bio-Rad, catalog number 7326008), and the concentration of the cleaved protein was determined using the Bio-Rad Protein Assay (Assay Kit I (5000002)). At this point, to make the ΔFOSB/JUND bZIP heterodimer, both proteins were combined at a 1:1 molar ratio and exhaustively dialyzed against 20 mM HEPES pH 7.0, 500 mM NaCl at room temperature. ΔFOSB bZIP homomer preparations were dialyzed against 20 mM HEPES pH 7.0, 500 mM NaCl at room temperature as well. As a final step, ΔFOSB/JUND bZIP and ΔFOSB bZIP preparations were concentrated to a maximum of 5 mg/mL and further purified using size exclusion chromatography at 4 °C using a Superdex HiLoad 16/600 75 pg column (Cytiva) equilibrated with GF Buffer (20 mM HEPES pH 7.0, 500 mM NaCl). Protein purity was assessed by SDS-PAGE. Proteins were concentrated to ∼8 mg/mL, flash-frozen in liquid nitrogen, and stored at −80 °C until further use.

### High-Throughput Screening of Compound Libraries

High-throughput screening (HTS) was carried out using an FP assay to evaluate two chemical libraries, Chembridge NT638 Diversity Set1 (30,080 compounds) and Maybridge (14,400 compounds). The assay monitors the binding of full-length N(His)_6_-ΔFOSB/N(His)_6_-JUND heterodimers (ΔFOSB/JUND) and N(His)_6_-ΔFOSB homomers (ΔFOSB) to TMR-cdk5, as described previously (12, 19, 32). Use of the full-length proteins, which contain intrinsically disordered regions, increases the signal in fluorescence polarization experiments, due to their larger size compared to only the bZIP domains, thus facilitating HTS. For the primary screen, compounds were transferred from library stock plates to 384-well assay plates (Corning #3676) using the Echo 550 Acoustic Liquid Transfer Machine (Beckman, Fullerton, CA) at a final concentration of 50 μM. TMR-cdk5 oligonucleotide was added to all wells at a final concentration of 25 nM. Subsequently, either ΔFOSB/JUND (280 nM monomer concentration; in 20 mM HEPES pH 7.5, 150 mM NaCl) or ΔFOSB (300-320 nM monomer concentration; in 20 mM HEPES pH 7.5, 50 mM NaCl) was added to wells containing compounds and TMR-cdk5. Wells containing 25 nM TMR-cdk5 alone were used as positive controls, representing 100% inhibition (no protein-DNA-binding), and wells containing 25 nM TMR-cdk5 plus protein (either 280 nM ΔFOSB/JUND or 320 nM ΔFOSB) without any compound were used as negative controls, representing 0% inhibition (full protein-DNA-binding). Buffer was used as the ‘no-protein’ control. All wells, including controls, received DMSO at a consistent final concentration of 0.5% (v/v). We used compound C1 (Chembridge 6572652) as a reference for calculating quality control and assay performance metrics (19, 32). The total reaction volume in each well was 20 μL. Plates were incubated at room temperature for 15 minutes before measuring FP signals using a BioTek Synergy Neo2 plate reader (excitation: 530 nm; emission: 590 nm).

We calculated assay quality-control metrics throughout the screening campaign following the procedures outlined in the Assay Guidance Manual (54). During the screening of ΔFOSB/JUND heterodimers, the FP signal window was 64.0 ± 6.2 mP, with minimal technical noise (mean coefficient of variance: 1.66% for negative controls and 2.30% for positive controls), and a robust Z’ score of 0.71 ± 0.07. For the screening of ΔFOSB homomers, similarly, the FP signal window was 63.80 ± 2.26 mP, with mean coefficients of variance of 1.49% and 2.08% for negative and positive controls, respectively, and a Z’ score of 0.73 ± 0.04.

Hits were identified using a rule-based classifier. The first-pass criterion removed compounds that interact with the TMR-cdk5 oligo directly (in the absence of protein) by retaining only those compounds for which the absolute value for FP (anisotropy) |FA| was ≤0.1 in the presence of DNA (i.e., equal or less than 10% difference from the positive control, TMR-cdk5, alone). The fraction affected was defined as

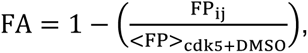

whereby FP_ij_ is the FP value for each assay well position, and <FP>_cdk5+DMSO_ is the median FP of the wells containing buffer including 0.5% DMSO and the TMR-cdk5 oligo. The second-pass criterion selected compounds with a high ability to disrupt the binding of protein to DNA and was defined as a Score of ≥ 0.75 and ≤ 1.25. The Score was calculated by first normalizing the FP signal from each well by subtracting the median value of the on-plate negative control (cdk5 oligo + protein in buffer + DMSO, i.e., no effect from compound) and then scaling it to the difference between the median FP signal of the on-plate positive (cdk5 oligo alone, i.e., 100% inhibition) and negative controls (cdk5 oligo + protein, i.e., 0% inhibition) using the following formula:

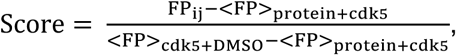

whereby FP_ij_ is the normalized and scaled FP for each assay well position; <FP>_cdk5+DMSO_ is the median FP of the wells containing the TMR-cdk5 oligo; and <FP>_protein+cdk5_ is the median FP of wells containing ΔFOSB/JUND or ΔFOSB and TMR-cdk5 oligo. Median values were used instead of mean values to facilitate identifying outliers in the empirically collected data.

### Fluorescence Polarization-Based Assay of the Effect of Compounds on ΔFOSB/JUND Binding to DNA

To confirm compound effects observed in HTS, candidate hits identified in our initial screen were purchased from vendors (see above) or synthesized in-house (JPC0661) and tested in a 10-point dose-response assay to validate their hit status and obtain estimates of their IC_50_ values, as we have done before (19, 32). A serial dilution of compound (0-200 μM or 0-250 μM) was dispensed into 384-well microtiter plates (Corning #3676) using an Echo 550 Acoustic Liquid Transfer Machine (Beckman, Indianapolis, IN) manually. Subsequently, either ΔFOSB/JUND (280 nM monomer concentration) or ΔFOSB (300-320 nM monomer concentration) was added to the wells containing compounds and TMR-cdk5. DRC assays were carried out in quadruplicates for ΔFOSB/JUND (in buffer 20 mM HEPES pH 7.5, 150 mM NaCl), in quadruplicates for ΔFOSB (in buffer 20 mM HEPES, pH 7.5, 50 mM NaCl), or in duplicates without protein. An end concentration of 2% (v/v) DMSO was maintained in all wells, which we separately ascertained did not impact the assay. Samples were incubated for 15 minutes at room temperature, and the FP signal was measured using a Synergy Neo2 plate reader (BioTek, excitation 530 nm, emission 590 nm) or Pherastar plate reader (BMG LabTech, excitation 540 nm, emission 590 nm). Each 384-well plate also included 16 positive control wells (TMR-cdk5 alone representing ‘100% inhibition’) and 16 negative control wells (protein+TMR-cdk5 representing ‘0% inhibition’) in the appropriate FP buffer. The concentration series of compounds with TMR-cdk5 but without protein served to identify compounds interfering with the assay or binding to TMR-cdk5 directly. The averaged data were fit in PRISM 6 (GraphPad) using the “Log-inhibitor Vs. Response (3 parameters)” fit.

### Electrophoretic Mobility Gel Shift Assay

The ability of compounds to disrupt full-length ΔFOSB/JUND-binding to DNA was evaluated using electrophoretic mobility shift assays (EMSAs) with the Thermo Scientific LightShift EMSA Optimization and Control Kit. Full-length ΔFOSB/JUND (50 or 100 nM) was incubated with 5 nM biotin-labeled oligonucleotide (BIO-cdk5) and varying concentrations of the compounds (0, 1, 10, 20, or 50 μM) in a reaction buffer containing 10 mM Tris (pH 8.0), 100 mM NaCl, 30 mM KCl, 1 mM EDTA, 10 μg/mL BSA, 2.5% (v/v) glycerol, and 0.1% (v/v) NP-40. Controls included 5 nM BIO-cdk5 alone (unshifted, 0%-binding) and ΔFOSB/JUND + BIO-cdk5 (fully shifted, 100%-binding) without compound. Samples were incubated at room temperature for 15 minutes, mixed with bromophenol loading dye (3% (w/v) Ficoll, 0.02% (w/v) bromophenol, 0.25×TBE (44.5 mM Tris, 44.5 mM boric acid, 1 mM EDTA, pH 8.3)), and resolved on pre-run, 5% native polyacrylamide gels on ice at 100 V in 0.5×TBE buffer. Following electrophoresis, the samples were electroblotted at 50 V for 40 minutes at 4 °C onto positively charged nylon membranes (Invitrogen) in 0.5×TBE. Membranes were UV-crosslinked for 10–12 minutes to fix the protein-DNA complexes to the membrane. Detection of the BIO-cdk5 oligo was performed using the Thermo Scientific Pierce Chemiluminescent Nucleic Acid Detection Module. Streptavidin-horseradish peroxidase conjugate (1:300 dilution) was combined with a luminol/enhancer solution and peroxide solution for chemiluminescence development. Membranes were briefly exposed to X-ray film (Kodak BioMax Light), and the film was developed for visualization.

### Crystallization and X-Ray Data Collection

ΔFOSB/JUND bZIP crystals were grown at 293 K using the hanging-drop vapor-diffusion method. Protein solution (2–4 μL of ΔFOSB/JUND bZIP, 8–10 mg/mL in 20 mM HEPES, pH 7.0, 250 mM NaCl, 1 mM TCEP) was mixed with an equal volume of reservoir solution (100 mM Tris, pH 8.5, 35% (v/v) isopropanol or 100 mM Tris, pH 7.0, 40% (v/v) ethanol). Crystals grew to 0.1–0.3 mm in 7 days. For ligand soaking, crystals were incubated with 10-20 mM JPC0661_comm_ for 1 h, cryo-protected in reservoir solution containing JPC0661_comm_, and flash-cooled in liquid nitrogen. X-ray diffraction data were collected at beamline 5.01 (Advanced Light Source, Berkeley, CA) at a wavelength of 0.9774 Å. Crystals belonged to space group I222 with unit cell parameters of **a** ∼48 Å, **b** ∼65–69 Å, and **c** ∼122–123 Å, containing one heterodimer per asymmetric unit. Diffraction extended to ∼1.72 Å resolution. The crystals displayed anisotropy and fiber-like diffraction patterns affected data quality and completeness.

### Structure Determination, Experimental Phasing, and Structural Analysis

Data were processed with autoPROC (55) and included anisotropy correction carried out with STARANISO (56). The crystal structure of ΔFOSB/JUND^cmpd^ was determined by molecular replacement using the structure of the reduced ΔFOSB/JUND bZIP (PDB ID: 7UCC) as the search model in Phaser (57) as implemented in Phenix (58). Model refinement, interspersed with manual rebuilding using COOT (59), was carried out in Phenix as well and included Translation-Libration-Screw rotation (TLS) parameterization. The models were validated with MolProbity (60) with Ramachandran statistics of 99.23% favored, 0.77% allowed, and 0% disallowed, and a clashscore of the final model of 3.28. There is one rotamer outlier. The refined structure exhibits relatively high R_work_ and R_free_ values, most likely due to the severe anisotropic diffraction, which is common for these crystals. For detailed statistics for data collection and refinement (see **Table 1**). Coordinates and structure factors for βFOSB/JUND^cmpd^ were deposited with the Protein Data Bank (PDB ID: 9OC3).

### Toxicity Assays

Generation of an expandable neural progenitor cell line (NPC; 8330-8 RC1) through an induced pluripotent stem cell (iPSC) intermediate from human fibroblasts (GM8330, Coriell Institute for Medical Research, Camden, NJ) has been previously described (61). NPCs were cultured in NPC medium composed of Dulbecco’s modified Eagle’s medium (DMEM, 70%, Gibco catalog number 11995) with nutrient supplements Ham’s F-12 (30%, Mediatech catalog number 10-080-CV) and B27 (2%, Gibco catalog number 17504-044), and 1% penicillin-streptomycin (Gibco), supplemented with epidermal growth factor (EGF, 20 ng/mL, Sigma), basic fibroblast growth factor (bFGF, 20 ng/mL, Reprocell), and heparin (5 μg/mL, Sigma) and allowed to proliferate at 37 °C with 5% CO_2_. NPCs were seeded at 6,000 cells/well in a volume of 50 μL on 384-well plates in NPC medium that were previously coated sequentially with 20 μg/mL poly-ornithine (Sigma, P3655) in water for 3 h at 37°C with 5% CO_2_ and then 5 μg/mL laminin (Sigma, L2020) diluted in PBS overnight at 37°C with 5% CO_2_. After 24 h of proliferation, compounds were pin-transferred to inner wells to indicated concentrations using a CyBio vario pin tool, with all wells DMSO-normalized to 0.13%. After the indicated time points (24 h in triplicate or 72 h in duplicate), plates were removed from the incubator and equilibrated to room temperature for 15 min, followed by dispensing of 25 µL of CTGlo 2.0 reagent (Promega G9242). Plates were allowed to nutate at room temperature for 15 minutes before reading luminescence using a PerkinElmer EnVision 2103 Multilabel Reader.

### Mouse Liver Microsomal Stability Assays

Compound stability was tested in mouse liver microsomes by Cyprotex. Briefly, microsomes (0.3 mg/mL) were incubated with compound at a final concentration of 1 μM (using a 10 mM stock in 100% DMSO) at 37 °C in the presence of 1 mM NADPH (used as a co-factor for compound-metabolizing enzymes) in potassium phosphate/MgCl_2_ containing buffer with a DMSO end concentration of 0.25% (v/v). Samples were removed at the time points 0, 5, 15, 30, and 45 min, and the reaction was terminated by the addition of acetonitrile. Following centrifugation, an internal standard was added to the supernatant, and the samples were analyzed by LC-TOF-MS. The disappearance of the test compound as a function of time was plotted, and intrinsic clearance (CL_int_ in μL/min/mg protein), standard error of the intrinsic clearance (SE CL_int_), and half-life (t½, in min) were calculated.

### Cell-Based Reporter and Toxicity Assays in AP1-luc HEK293 Cells

The experimental approach for our cell-based reporter and toxicity assays followed published methods (19). The AP1-luciferase reporter human embryonic kidney (HEK293) cell line was obtained from BPS Bioscience (USA) and maintained in growth medium 1B (BPS Bioscience) supplemented with 10% fetal bovine serum (FBS, Invitrogen) and 1% penicillin/streptomycin (Hyclone) under standard incubation conditions of 37 °C with 5% CO_2_. The AP1 reporter luciferase assay was used to investigate the impact of test compounds on the transcriptional activity of the AP1-response-element-driven luciferase reporter gene.

Cells were seeded in quadruplicate across two 48-well microplates at a density of 6.0×10⁴ cells per well. Once stabilized and having reached about 70% confluence, the culture medium was replaced with assay medium 1B containing 0.5% FBS, and the cells underwent a 24-h serum starvation period. Serial dilutions of test compounds (ranging from 0.003 to 100 μM) were then applied. After a 2-h pre-incubation, cells were either stimulated with 20% FBS (Invitrogen) (‘serum-stimulated’), maintained in serum-free conditions (‘non-serum-stimulated’) in growth medium 1B, or treated with 0.5% DMSO (a concentration equivalent to that present in the 100 μM compound condition) (control) for 24 h. Following treatment, cells were lysed, and luminescence detection was performed using the luciferase assay system (Promega) per the manufacturer’s protocol with a BioTek Cytation 3 reader. Statistical analyses were performed using GraphPad Prism v.10.0 (GraphPad Software). Each compound was tested in two independent experiments, and data were normalized to the luciferase signals of blank wells from the corresponding experiments. Nonlinear regression was used to calculate IC_50_ values, which are reported with their 95% confidence interval (CI). Additionally, cell viability assays were conducted under conditions identical to the AP1 reporter assay to evaluate compound-induced toxicity. After serum stimulation, viability was determined using the CellTiter-Glo luminescent assay (Promega) according to the manufacturer’s protocol, and luminescence was measured using a BioTek Cytation 3 reader. Statistical analyses were performed in GraphPad Prism v.10.0 (GraphPad Software), and data are presented as means ± standard error of the mean (SEM).

### Chemical Synthesis of JPC0661 and Analogs

The detailed synthetic routes and experimental procedures of JPC0661 and its analogs YL0325, YL0327, YL0328, and YL0324 are provided in the supporting materials (**Supplementary Fig. S6**).

### *In-vivo* JPC0661 Compound Administration

*In vivo* studies were performed in 2–3-month-old heterozygous transgenic male and female mice expressing human amyloid precursor protein (APP) carrying Swedish (K670N, M671L) and Indiana (V717F) mutations linked to familial Alzheimer’s disease (Line J20; MMRRC_034836-JAX; hAPP770 transgene) (62). JPC0661 was delivered unilaterally into the dorsal hippocampus through a cannula connected to an Alzet micro-osmotic pump (model 1003D). Each micro-osmotic pump was filled with 0.2% DMSO/0.9% saline or 50 μM JPC0661 in 0.2% DMSO/0.9% saline per manufacturer’s instructions, connected to a brain infusion cannula via medical-grade polyvinyl chloride catheter tubing, and primed overnight before implantation. One micro-osmotic pump was implanted per mouse, subcutaneously in the intrascapular region, followed by fixation of the cannula base to the skull via Superglue, with the tip of the cannula targeting stereotactic coordinates: dorsal-ventral (D/V), –2.0 mm; anterior-posterior (A/P), –2.1 mm; and medial-lateral (M/L), –1.2 mm. Model 1003D micro-osmotic pumps delivered fluid at a rate of 0.96 μL/h for final drug delivery of 16.7 ng JPC0661/h. After three days of intra-hippocampal infusion of JPC0661 or vehicle, mice were anesthetized with commercial euthanasia solution, followed by transcranial perfusion with ice-cold 0.9% saline solution. Brains were collected and hemisected. The left hemibrain was fixed in 4% paraformaldehyde for immunohistochemistry, while the right hemibrain was flash-frozen on dry ice. The hippocampus from the right hemibrain was isolated, and the dorsal two-thirds sub-dissected and used for CUT&RUN analysis. All procedures were approved by the Institutional Animal Care and Use Committees of Baylor College of Medicine.

Fixed hemibrains were cryoprotected in 30% sucrose in phosphate-buffered saline before sectioning on a freezing sliding microtome. Serial coronal sections (30 μm thickness) were divided into 10 subseries, each containing every tenth section throughout the rostral-caudal extent of the brain. One subseries was stained for ΔFOSB with the avidin-biotin diaminobenzidine (DAB) method, using rabbit anti-ΔFOSB (1:5000; Cell Signaling, D3S8R) as the primary antibody, biotinylated goat anti-rabbit (1:200; Vector, BA-1000) as the secondary antibody, and DAB as the chromogen. All samples were processed, stained, and imaged contemporaneously using the same parameters and imaging settings. Quantification of ΔFOSB was performed by an experimenter blinded to treatment and confirmed ΔFOSB induction in the transgenic mice as observed previously. The mean gray value of the dentate gyrus granule cell layer was measured using ImageJ software, normalized to the mean gray value of the stratum radiatum, and averaged over two consecutive sections for each sample.

### CUT&RUN Sequencing

#### Nuclei Isolation

Nuclei were isolated from the dorsal hippocampus tissue of APP mice after infusion of vehicle or JPC0661. Tissue was homogenized in lysis buffer (320 mM sucrose, 5 mM CaCl_2_, 0.1 mM EDTA, 10 mM Tris-HCl, pH 8.0, 1 mM DTT, 0.1% Triton X-100, 1.5 mM spermidine, and protease inhibitors) using a Dounce homogenizer (30-40 strokes per pestle). Homogenates were filtered through a 40 μm strainer and transferred to ultracentrifuge tubes. A sucrose cushion (1.8 M sucrose, 10 mM Tris-HCl, pH 8.0, 1 mM DTT, 1.5 mM spermidine, and protease inhibitors) was layered beneath, and nuclei were pelleted by ultracentrifugation at 24,000 rpm for 1.25 h at 4 °C.

#### CUT&RUN Assay

CUT&RUN was performed as described previously (18) with slight modifications. Nuclei were resuspended in wash buffer (20 mM HEPES-NaOH, pH 7.5, 150 mM NaCl, 0.5 mM spermidine, 0.1% Triton X-100, 0.1% Tween-20, 0.1% BSA, and protease inhibitors) and incubated with BioMag®Plus Concanavalin A beads (BP531, Bang Laboratories) in binding buffer (20 mM HEPES-KOH, pH 7.9, 10 mM KCl, 1 mM CaCl_2_, and 1 mM MnCl_2_) for 10 min at room temperature. Beads were magnetically separated and incubated overnight at 4 °C with primary anti-ΔFOSB antibody (1:50) or IgG control (1:50). After washes, beads were incubated with pAG-MNase (EpiCypher) for 1 h at 4 °C. Following additional washes in low-salt buffer, beads were incubated in calcium buffer (3.5 mM HEPES-NaOH, pH 7.5, 10 mM CaCl_2_, 0.1% Triton X-100, and 0.1% Tween-20) for 5 min on a pre-chilled 4 °C block, then pAG-MNase digestion was stopped with ED:ET STOP buffer (170 mM NaCl, 20 mM EDTA, 4 mM EGTA, 0.1% Triton X-100, 0.1% Tween-20, 25 μg/mL RNase, and 20 μg/mL glycogen). DNA was eluted at 37 °C for 30 min with shaking, centrifuged at 16,000 RCF for 5 min at 4 °C, and purified using ethanol precipitation. Libraries were generated using the NEB Next® Ultra™ II DNA Library Prep Kit (New England BioLabs) and sequenced on an Illumina HiSeq 4000 with a 2 × 150 bp paired-end configuration at a minimum depth of 20 million reads per sample.

#### Data Analysis

FASTQ files were processed with the NGS-Data-Charmer pipeline (https://github.com/shenlab-sinai/NGS-Data-Charmer). Adapter trimming was performed using Trim-Galore (v0.6.5), followed by secondary trimming with Cutadapt (v2.10). Reads were aligned to the mm10 genome using HISAT2 (v2.2.0), and duplicate reads were removed using the Picard (v3.0) ‘MarkDuplicates’ module. For visualization purposes, de-duplicated reads were converted to bigwig files with the Deeptools (v3.5.0) ‘bamCoverage’ module (--binSize 10 --normalizeUsing RPKM), then visualized as tracks using IGVtools (v2.5.3). Peak calling was performed with MACS2 (v2.2.6) (-f BAMPE -q 0.01 --keep-dup) using the IgG pooled from each APP mouse as background subtraction. Peak files were annotated using ChIPseeker (v1.22.1), followed by heatmap creation in each group. Differential peak correlations were visualized using DiffBind (v3.9) with p < 0.0001, merged, and filtered at an FDR of 10%. Bigwig files were visualized using IGV (v2.12.2). Gene Ontology analysis was conducted using ShinyGO, with significant terms filtered at fold enrichment ≥1.5 and FDR Q <0.05. No differences were seen between male and female mice; hence, all analyses combined the two sexes. Known Motif Enrichment analyses were performed using Homer software (v4.9) (63).

### Impact of compounds on the ΔFOSB/JUND bZIP

To assess the impact of compounds on the ΔFOSB/JUND bZIP, and their potential to prevent closure of the redox switch, we incubated purified protein with compounds in presence and absence of the oxidizing agent, diamide. Briefly, 500 μgr of ΔFOSB/JUND bZIP in 1000 μL 20mM HEPES pH 7.0, 250mM NaCl was incubated with 100 μM TCEP for 60 minutes on ice to ensure the protein was fully reduced. The protein was then dialyzed three times in 20mM HEPES pH 7.0, 250mM NaCl for 15 minutes each against ∼600 mL volume of 20mM HEPES pH 7.0, 250mM NaCl buffer. The protein was subsequently aliquoted into samples of 27 μgr each, 0.5 mM compound end concentration added followed by incubation for 15 min at RT, and then 100 μM diamide (‘ox’) end concentration or an equivalent volume 0.3 μL buffer (20mM HEPES pH 7.0, 250mM NaCl) (‘red’) added to samples followed by incubation for 15 min at room temperature. Samples were then subjected to SDS-PAGE using loading buffer with reducing agent (disrupting disulfide bonds) or without (leaving disulfide bonds intact).

To assess the impact of compounds on the ΔFOSB/JUND bZIP and their potential to promote closure of the redox switch, we carried out the same procedure as above, except that the protein after reduction with TCEP was dialyzed three times in 20 mM HEPES pH 7.0, 250 mM NaCl for 30 minutes each against ∼600 mL volume of 20 mM HEPES pH 7.0, 250 mM NaCl buffer to remove any traces of TCEP, before adding the different compounds in absence of any diamide (‘red’).

## Data Availability

The atomic coordinates and structure factors for the reported crystal structure have been deposited in the Protein Data Bank under accession code **PDB ID 9OC3**. All CUT&RUN sequencing data generated in this study have been deposited in the NCBI Gene Expression Omnibus (GEO) under accession number **GSE304157**.

## Supporting Information

This article contains supporting information.

## Acknowledgements

GR gratefully acknowledges the University of Texas Medical Branch (UTMB) for providing research resources. The Advanced Photon Source (sectors IMCA-CAT and LS-CAT), the Advanced Light Source (ALS), and the National Synchrotron Light Source II (NSLS2) are thanked for access to synchrotron radiation. Drs. Dan Fass, Hubert Lee, Anthony J Pastore, Zachary Rosenthal, and Qichao Bao are thanked for their support with preliminary pilot studies, experimental assistance, and useful discussions.

## Funding and additional information

This work was supported by the National Institutes of Health, National Institute on Drug Abuse [grant numbers R01DA040621, R01DA040621-03S1, R01DA040621-07S1]; the John D. Stobo, M.D. Distinguished Chair Endowment Fund and the Edith & Robert Zinn Chair in Drug Discovery Endowment Fund [to JZ]; the Jeane B. Kempner Award (University of Texas Medical Branch) [to SM]; the Stuart & Suzanne Steele Massachusetts General Hospital Research Scholar Award and internal support through the Precision Therapeutics Unit in the Center for Genomic Medicine at Massachusetts General Hospital [to SJH]; the Cancer Prevention and Research Institute of Texas, Combinatorial Drug Discovery Program [grant number RP150578]; the National Institutes of Health, National Institute of Neurological Disorders and Stroke [grant number R01NS085171 to JC]; and the National Institutes of Health, National Institute on Aging [grant number F30AG085919 to CPS]. Funding for open access charge: National Institutes of Health, National Institute on Drug Abuse [R01DA040621]. The content is solely the responsibility of the authors and does not necessarily represent the official views of the National Institutes of Health.

## Abbreviations

a.a.: amino acid
Aβ: Amyloid β protein
AD: Alzheimer’s disease
AP1: Activator protein 1
APP: amyloid precursor protein
BSA: bovine serum albumin
bZIP: basic leucine zipper
CI: confidence interval
CL: clearance
DMSO: dimethyl sulfoxide
DOPA: L-3,4-dihydroxyphenylalanine
DRC: dose-response curve
DMEM: Dulbecco’s modified Eagle’s medium
DTT: dithiothreitol
D/V: dorsal/ventral
EGF: epidermal growth factor
FBS: fetal bovine serum
FP: fluorescence polarization
GO: gene ontology
HEPES: 4-(2-hydroxyethyl)-1-piperazineethanesulfonic acid
HTS: High-throughput screening
iPSC: induced pluripotent stem cell
IPTG: isopropyl-D-thiogalactoside
LB: Luria-Bertani
LC: liquid chromatography
MOI: multiplicity of infection
MS: mass spectrometry
NTA: nickel-nitrilotriacetic acid complex
NPC: Neural progenitor cells
pAG-MNase: protein A-G micrococcal nuclease
PBS: phosphate-buffered saline
PDB: Protein Data Bank
Pfu: plaque-forming unit
PMSF: phenylmethylsulfonyl fluoride
r.m.s.: root-mean-square
SE: standard error
SEM: standard error of the mean
TAMRA: tetramethylrhodamine
TLS: translation, libration, screw rotation
TCEP: tris(2-carboxyethyl)phosphine
TEV: tobacco etch virus
TOF: time of flight
Tris: tris(hydroxymethyl)aminomethane

