## Supplemental Material for "Discovery of Small Molecules and a Druggable Groove That Regulate DNA Binding and Release of the AP1 Transcription Factor ΔFOSB"

† Joint First Authors

* To whom correspondence should be addressed.

### Present Addresses:

### RUNNING TITLE

Small Molecules Reveal a Novel Druggable Groove in ΔFOSB

a)

| compound | FP-assay activity $IC_{50}$ | |
|---|---|---|
| | ΔFosB/JunD | ΔFosB |
| $JPC0661_{comm}$ | 9.1 μM (95% CI: 6.5 to 12.7 μM)<br>8.6 μM (95% CI: 7.8 to 9.5 μM) | 67.7 μM (95% CI: 60.6 to 75.6 μM)<br>52.1 μM (95% CI: 40.3 to 67.4 μM) |
| $JPC0661_{comm}$ + TCEP | 43.6 μM (95% CI: 31.7 to 59.9 μM)<br>33.3 μM (95% CI: 30.8 to 36.0 μM) | 74.9 μM (95% CI: very wide)<br>84.0 μM (95% CI: 57.1 to 123.7 μM) |
| HTS 10307 | 3.3 μM (95% CI: 2.1 to 5.0 μM)<br>1.5 μM (95% CI: 0.3 to 8.2 μM) | 7.1 μM (95% CI: 5.8 to 8.7 μM)<br>6.7 μM (95% CI: 4.2 to 10.8 μM) |
| HTS 10307 + TCEP | 14.7 μM (95% CI: 11.1 to 19.4 μM)<br>3.7 μM (95% CI: 0.8 to 16.8 μM) | 13.4 μM (95% CI: 10.9 to 16.5 μM)<br>22.4 μM (95% CI: 13.3 to 37.6 μM) |

| compound | FP-assay activity $IC_{50}$ | |
|---|---|---|
| | ΔFosB/JunD | ΔFosB |
| JPC0661 | 13.8 μM (95% CI: 11.0 to 17.2 μM)<br>8.9 μM (95% CI: 5.7 to 13.9 μM) | 9.3 μM (95% CI: 6.8 to 12.8 μM)<br>14.8 μM (95% CI: 10.7 to 20.4 μM) |
| YL0325 | 9.6 μM (95% CI: 0.3 to 321.5 μM)<br>18.2 μM (95% CI: 7.0 to 47.2 μM) | 58.0 μM (95% CI: 42.9 to 78.4 μM)<br>50.5 μM (95% CI: 37.4 to 68.3 μM) |
| YL0327 | 32.9 μM (95% CI: 22.8 to 47.7 μM)<br>52.0 μM (95% CI: 25.2 to 107.2 μM) | 116.2 μM (95% CI: 90.1 to 149.9 μM)<br>113.7 μM (95% CI: 90.2 to 143.3 μM) |
| YL0328 | 60.8 μM (95% CI: 47.1 to 78.6 μM)<br>86.7 μM (95% CI: 55.7 to 134.7 μM) | 97.2 μM (95% CI: 85.9 to 109.9 μM)<br>104.1 μM (95% CI: 72.2 to 150.0 μM) |
| YL0324 | >> 200 μM<br>>> 200 μM | >> 200 μM<br>>> 200 μM |

b)

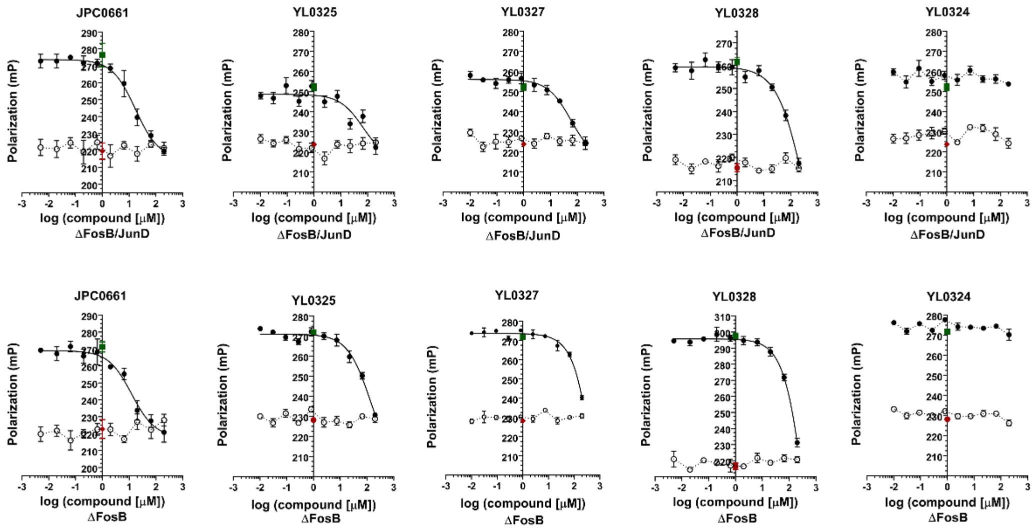


c)

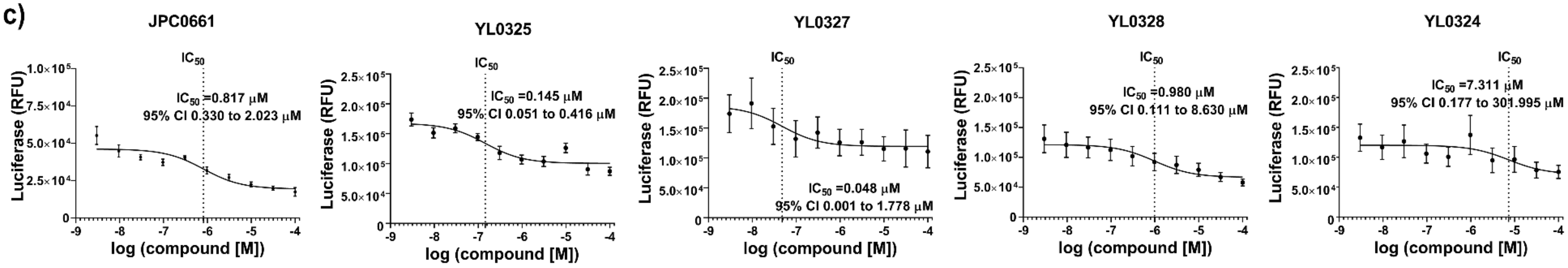


**Figure S1. Validation of JPC0661. A)** Summary table of FP-DRC experiments carried out for $JPC0661_{comm}$, HTS 10307, and a panel of JPC0661 analogs ($IC_{50}$ values and 95% confidence intervals (CI) are shown for each of two independent experiments) testing against both ΔFOSB/JUND heterodimers and ΔFOSB homomers. **B)** Representative FP-DRC assays carried out for JPC0661 and a panel of analogs. Experiments were carried out in independent duplicates. The $IC_{50}$ values are listed in **A)**. **C)** Representative dose-response curves for JPC0661 and a panel of analogs tested in AP1-reporter assays using AP1-luc HEK293 cells. Each compound was tested at least twice in independent experiments (typically n=3–4 wells per experiment), with a total of n=7–8 replicates for YL0325, n=5–6 for YL0327, and n=10–12 for YL0328 and YL0324. Results were normalized to the luciferase signals of blank wells from both experiments (n=6–8). Nonlinear regression using a three-parameter model was applied to fit the luciferase signal and to calculate $IC_{50}$ values which are presented with their 95% confidence interval (CI). Data points represent the mean of the normalized replicates and error bars the SEM. Note that the plot for JPC0661 is identical to **Fig. 2D** and is shown again here to facilitate comparison.

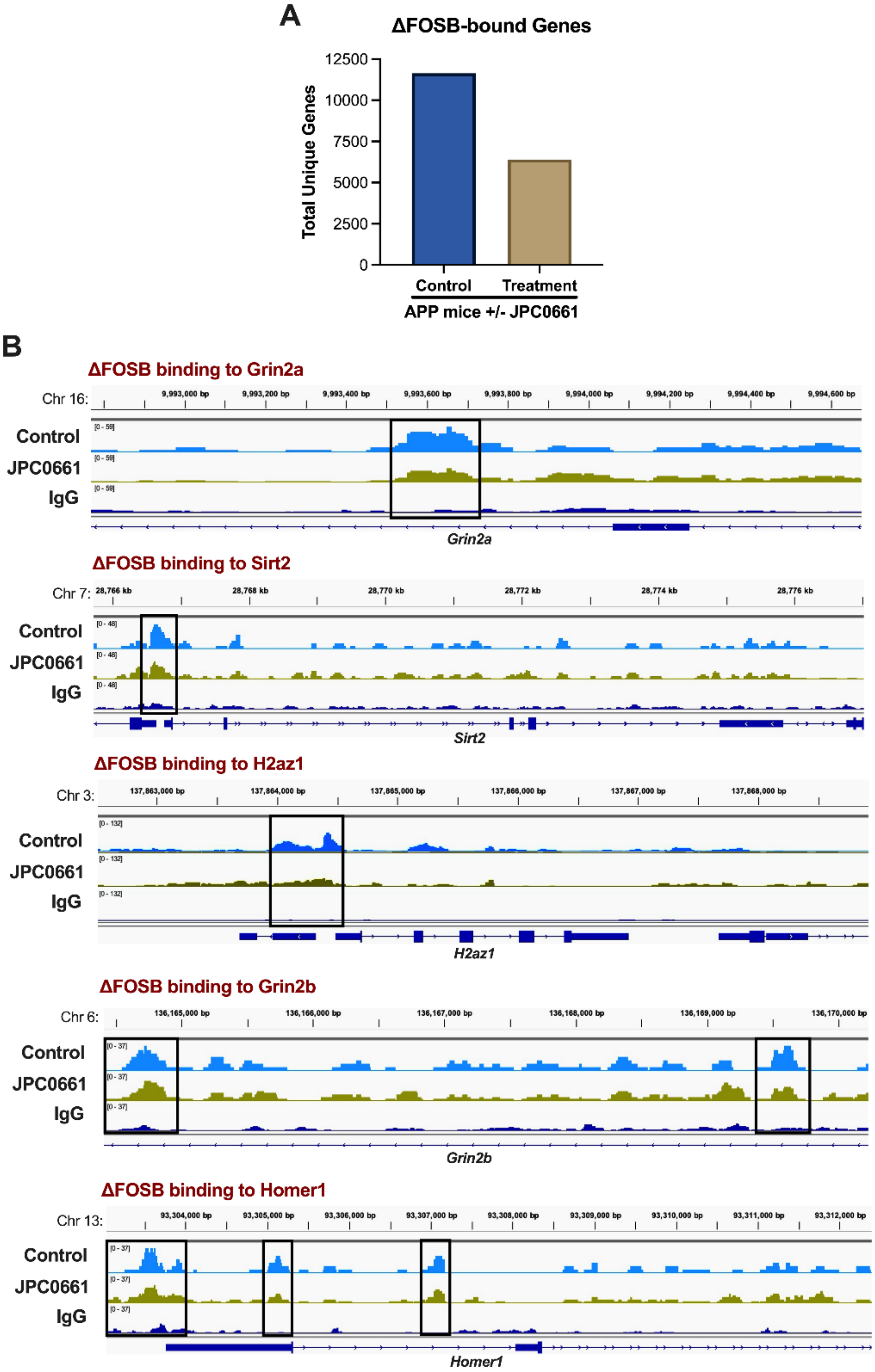
A
ΔFOSB-bound Genes
Total Unique Genes
12500
10000
7500
5000
2500
0
Control
Treatment
APP mice +/- JPC0661
B
ΔFOSB binding to Grin2a
Chr 16:
Control
JPC0661
IgG
Grin2a
ΔFOSB binding to Sirt2
Chr 7:
Control
JPC0661
IgG
Sirt2
ΔFOSB binding to H2az1
Chr 3:
Control
JPC0661
IgG
H2az1
ΔFOSB binding to Grin2b
Chr 6:
Control
JPC0661
IgG
Grin2b
ΔFOSB binding to Homer1
Chr 13:
Control
JPC0661
IgG
Homer1

**Figure S2. Effect of JPC0661 on ΔFOSB binding in dorsal hippocampus *in vivo*. A)** Total number of unique ΔFOSB-bound genes in vehicle-treated *vs.* JPC0661-treated APP mice shows an approximately 50% reduction in ΔFOSB binding after JPC0661 treatment. **B)** Genome browser tracks showing ΔFOSB binding at examples of known *in vivo* target genes for ΔFOSB (*Grin2a, Sirt2, H2az1, Grin2b,* and *Homer1*), demonstrating a reduction in ΔFOSB binding after JPC0661 treatment. Tracks represent CUT&RUN-sequencing of dorsal hippocampus from APP mice treated with vehicle (top, sky blue) or JPC0661 (middle, gold-green) using an anti-ΔFOSB antibody, with IgG control (bottom, dark blue) for non-specific binding and background subtraction.

| Gene | UniProt ID | Gene name | bZIP | Sequence |
|---|---|---|---|---|
| FOS | P01100 | FOS | 137-200 | EEKRRIRRERNKMAAAKCRNRRRELTDTLQAETDQLEDEKSALQTEIANLLKEKEKLEFILAAH |
| FOSB | P53539 | FOSB | 155-218 | EEKRRVRRERNKLAAAKCRNRRRELTDRLQAETDQLEEEKAELESEIAELQKEKERLEFVLVAH |
| FOSL1 | P15407 | FOSL1 | 105-168 | EERRRVRRERNKLAAAKCRNRRK-LTDFLQAETDKLEDEKSGLQREIEELQKQKERLELVLEAH |
| FOSL2 | P15408 | FOSL2 | 124-187 | EEKRRIRRERNKLAAAKCRNRRRELTEKLQAETEELEEEKSGLQKEIAELQKEKEKLEFMLVAH |
| JDP2 | Q8WYK2 | JDP2 | 72-135 | EERRKRRREKNKVAAARCRNKKKERTEFLQRESERLELMNAELKTQIEELKQERQQLILMLNRH |
| JUN | P05412 | JUN | 252-315 | RIKAERKRMRNRIAASKCRKRKLERIARLEEKVKTLKAQNSELASTANMLREQVAQLKQKVMNH |
| JUNB | P17275 | JUNB | 268-331 | RIKVERKRLRNRLAATKCRKRKLERIARLEDKVKTLKAENAGLSSTAGLLREQVAQLKQKVMTH |
| JUND | P17535 | JUND | 268-331 | RIKAERKRLRNRIAASKCRKRKLERISRLEEKVKTLKSQNTELASTASLLREQVAQLKQKVLSH |
| | | | | ^ ^ ^^ ^^^ |

**Figure S3. Multi-sequence alignment of human FOS and JUN family member bZIPs.** Residues in ΔFOSB and JUND interacting with JPC0661 are highlighted in blue (interacting with Lig 1), red (interacting with Lig 2), and olive green (interacting with both Lig 1 and Lig 2), respectively. The positions of these residues within the bZIP regions are highlighted at the bottom of the shown sequences.

| Gene | UniProt ID | Gene name | bZIP | Sequence |
| --- | --- | --- | --- | --- |
| ATF1 | P18846 | ATF1 | 213-271 | QLKREIRLMKNREAARECRRKKKEYVKCLENRVAVLENQNKTLIEELKTLKDLYSNKSV----- |
| ATF2 | P15336 | ATF2 | 352-415 | DEKRRKFLERNRAAASRCRQKRKVWVQSLEKKAEDLSSLNGQLQSEVTLLRNEVAQLKQLLLAH |
| ATF3 | P18847 | ATF3 | 86-149 | DERKKRRRERNKIAAAKCRNKKKEKTECLQKESEKLESVNAELKAQIEELKNEKQHLIYMLNLH |
| ATF7 | P17544 | ATF7 | 332-395 | DERRQRFLERNRAAASRCRQKRKLWVSSLEKKAEELTSQNIQLSNEVTLLRNEVAQLKQLLLAH |
| BACH1 | O14867 | BACH1 | 557-620 | CIHDIRRRSKNRIAAQRCRKRKLDCIQNLESEIEKLQSEKESLLKERDHILSTLGETKQNLTGL |
| BACH2 | Q9BYV9 | BACH2 | 646-709 | FIHDVRRRSKNRIAAQRCRKRKLDCIQNLECEIRKLVCEKEKLLSERNQLKACMGELLDNFSCL |
| CREB1 | P16220 | CREB1 | 269-327 | ARKREVRLMKNREAARECRRKKKEYVKCLENRVAVLENQNKTLIEELKALKDLYCHKSD----- |
| CREBL2 | O60519 | CRBL2 | 23-86 | KIDLKAKLERSRQSARECRARKKLRYQYLEELVSSRERAICALREELEMYKQWCMAMDQGKIPS |
| CREB5 | Q02930 | CREB5 | 375-438 | DERRRKFLERNRAAATRCRQKRKVWVMSLEKKAEELTQTNMQLQNEVSMLKNEVAQLKQLLLTH |
| CREM | Q03060 | CREM | 286-345 | TRKRELRLMKNREAAKECRRRKKEYVKCLESRVAVLEVQNKKLIEELETLKDICSPKTDY---- |
| FOS | P01100 | FOS | 137-200 | EEKRRIRRERNKMAAAKCRNRRRELTDTLQAETDQLEDEKSALQTEIANLLKEKEKLEFILAAH |
| FOSB | P53539 | FOSB | 155-218 | EEKRRVRRERNKLAAAKCRNRRRELTDRLQAETDQLEEEKAELESEIAELQKEKERLEFVLVAH |
| FOSL1 | P15407 | FOSL1 | 105-168 | EERRRVRRERNKLAAAKCRNRRK-LTDFLQAETDKLEDEKSGLQREIEELQKQKERLELVLEAH |
| FOSL2 | P15408 | FOSL2 | 124-187 | EEKRRIRRERNKLAAAKCRNRRRELTEKLQAETEELEEEKSGLQKEIAELQKEKEKLEFMLVAH |
| JDP2 | Q8WYK2 | JDP2 | 72-135 | EERRKRRREKNKVAAARCRNKKKERTEFLQRESERLELMNAELKTQIEELKQERQQLILMLNRH |
| JUN | P05412 | JUN | 252-315 | RIKAERKRMRNRIAASKCRKRKLERIARLEEKVKTLKAQNSELASTANMLREQVAQLKQKVMNH |
| JUNB | P17275 | JUNB | 268-331 | RIKVERKRLRNRLAATKCRKRKLERIARLEDKVKTLKAENAGLSSTAGLLREQVAQLKQKVMTH |
| JUND | P17535 | JUND | 268-331 | RIKAERKRLRNRIAASKCRKRKLERISRLEEKVKTLKSQNTELASTASLLREQVAQLKQKVLSH |
| MAF | O75444 | MAF | 288-351 | RLKQKRRTLKNRGYAQSCRFKRVQQRHVLESEKNQLLQQVDHLKQEISRLVRERDAYKEKYEKL |
| MAFA | Q8NHW3 | MAFA | 254-317 | RLKQKRRTLKNRGYAQSCRFKRVQQRHILESEKCQLQSQVEQLKLEVGRLAKERDLYKEKYEKL |
| MAFB | Q9Y5Q3 | MAFB | 238-301 | RLKQKRRTLKNRGYAQSCRYKRVQQKHHLENEKTQLIQQVEQLKQEVSRLARERDAYKVKCEKL |
| MAFF | Q9ULX9 | MAFF | 51-114 | RLKQRRRTLKNRGYAASCRVKRVCQKEELQKQKSELEREVDKLARENAAMRLELDALRGKCEAL |
| MAFG | O15525 | MAFG | 51-114 | QLKQRRRTLKNRGYAASCRVKRVTQKEELEKQKAELQQEVEKLASENASMKLELDALRSKYEAL |
| MAFK | O60675 | MAFK | 51-114 | RLKQRRRTLKNRGYAASCRIKRVTQKEELERQRVELQQEVEKLARENSSMRLELDALRSKYEAL |
| NFE2 | Q16621 | NFE2 | 266-329 | LVRDIRRRGKNKVAAQNCRKRKLETIVQLERELERLTNERERLLRARGEADRTLEVMRQQLTEL |
| NFE2L1 | Q14494 | NF2L1 | 654-717 | LIRDIRRRGKNKMAAQNCRKRKLDTILNLERDVEDLQRDKARLLREKVEFLRSLRQMKQKVQSL |
| NFE2L2 | Q16236 | NF2L2 | 497-560 | LIRDIRRRGKNKVAAQNCRKRKLENIVELEQDLDHLKDEKEKLLKEKGENDKSLHLLKKQLSTL |
| NFE2L3 | Q9Y4A8 | NF2L3 | 578-641 | LIRDIRRRGKNKVAAQNCRKRKLDIILNLEDDVCNLQAKKETLKREQAQCNKAINIMKQKLHDL |
| NRL | P54845 | NRL | 159-222 | RLKQRRRTLKNRGYAQACRSKRLQQRRGLEAERARLAAQLDALRAEVARLARERDLYKARCDRL |
| | | | | ^ ^ ^^ ^^^ |

**Figure S4. Multi-sequence alignment of all 29 human bZIP sequences containing cysteines in the same location as the redox-switch cysteines Cys172 in ΔFOSB and Cys285 in JUND, respectively.** Residues in ΔFOSB and JUND interacting with JPC0661 are highlighted in blue (interacting with Lig 1), red (interacting with Lig 2), and olive green (interacting with both Lig 1 and Lig 2), respectively. The positions of these residues within the bZIP regions are highlighted at the bottom of the shown sequences. Note, CREBL2 (CAMP Responsive Element Binding Protein Like 2) shows limited sequence conservation of the leucine zipper, but it was nevertheless included in order to evaluate its similarity in the compound binding region.

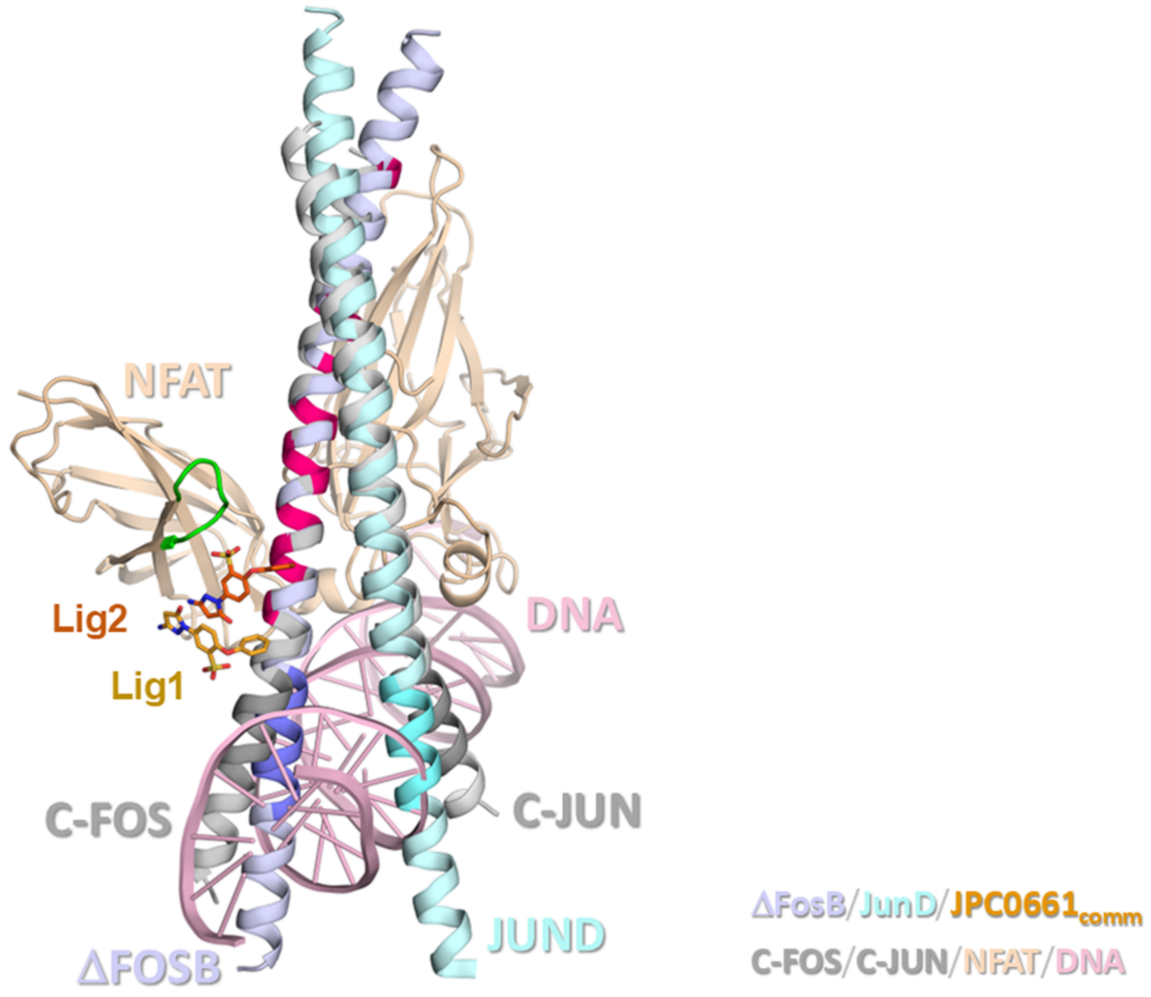


**Figure S5. Comparison of the ΔFOSB/JUND bZIP in complex with JPC0661$_{comm}$ (this study) and the complex of FOS/JUN/NFAT1/DNA (PDB ID: 1S9K).** ΔFOSB is shown in lilac and JUND in light cyan; the DNA-binding motifs are shown in dark lilac (ΔFOSB N165-Arg173) and dark cyan (JUND Asn278-Arg286), respectively. JPC0661$_{comm}$ molecules (Lig1 and Lig2) are shown with carbons in light and dark orange, respectively, oxygen in red, nitrogen in blue, and sulfur in yellow. In the FOS/JUN/NFAT1/DNA complex, FOS and JUN are shown in grey, NFAT1 in beige, and DNA in pink. The main-chain atoms of ΔFOSB Glu178-Gln189 and the corresponding residues in FOS were used to guide the superposition. Residues in FOS that contact NFAT1 within 5.0 Å are: Glu160, Asp163, Thr164, Gln166, Ala167, Thr169, Asp170, Gln171, Glu173, Asp174, Ser177, Gln180, Thr181, and Glu191. These residues correspond to ΔFOSB residues: Glu178, Asp181, Arg182, Gln184, Ala185, Thr187, Asp188, Gln189, Glu191, Glu192, Ala195, Glu198, Ser199, Glu209 (shown in hot pink).

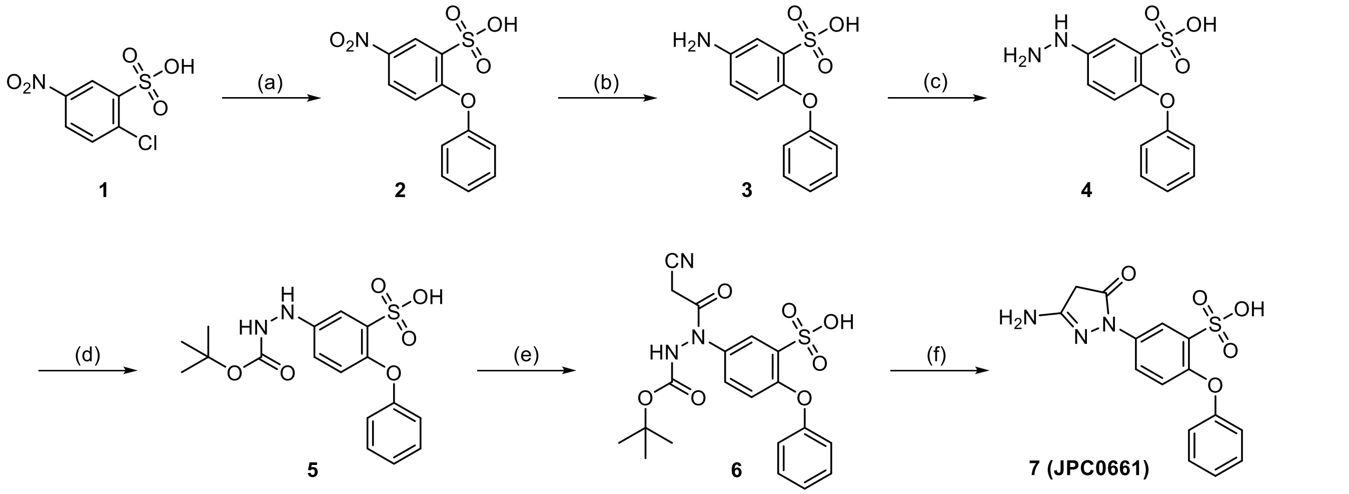


**Figure S6. Synthetic scheme for JPC0661. (a)** Phenol (1.2 equiv.), $K_2CO_3$ (2.0 equiv.), DMF (0.5 M), 100 °C, 24 h; 77% yield. **(b)** Pd/C (10 mol %), $H_2$, MeOH (0.5 M), 25 °C, 24 h; 82% yield. **(c)** $NaNO_2$ (1.1 equiv.), $SnCl_2 \cdot 2H_2O$ (2.0 equiv.), 12% w/w aq. HCl (excess), 0 ~ 25 °C, 12 h; 98% yield. **(d)** $Boc_2O$ (1.2 equiv.), TEA (1.5 equiv.), DMF:THF (v/v 1:1, 0.4 M), room temperature, 12 h; 65% yield. **(e)** 2-cyanoacetic acid (1.5 equiv.), PPAA (1.2 equiv.), TEA (3.0 equiv.), DCM (0.2 M), room temperature, 2 h; quantitative yield. **(f)** HCl/EtOAc (2 equiv., 0.2 M), room temperature, 2 h; 76% yield.

**The synthetic routes and experimental procedures of JPC0661 and the source of its analogs YL0324, YL0325, YL0327, and YL0328**

All chemicals and solvents were obtained from commercial suppliers and used without further purification unless specified. Analytical TLC was performed on silica gel 60F 254 plates (Merck) with UV detection at 254 nm. Preparative column chromatography was carried out on silica gel 60 (70−230 mesh, flash). NMR spectra were recorded on a Bruker spectrometer (400 or 800 MHz for 1H NMR and 201 MHz for 13C NMR) in DMSO-d6. Chemical shifts (δ) are reported in ppm, with TMS (δ = 0 ppm) or residual solvent signals (DMSO: δ = 2.5 ppm) as internal references for 1H NMR, and DMSO (δ = 39.51 ppm) for 13C NMR. Coupling constants are reported in Hz. The purity of the final compounds was determined by HPLC on a Shimadzu system (model: CBM-20ALC-20ADSPD-20AUV/vis) using a Waters μBondapak C18 column (300 × 3.9 mm), with a flow rate of 0.5 mL/min, UV detection at 270 and 254 nm. A linear gradient from 10% acetonitrile (ACN) in water containing 0.1% trifluoroacetic acid (TFA) to 100% ACN (0.1% TFA) over 20 min, followed by 30 min with the final solvent, was used. The compounds 5-Amino-2-(4-phenoxyphenyl)-2,4-dihydro-3H-pyrazol-3-one (YL0324; CAS# 3845-86-1), 5-(3-Methyl-5-oxo-4,5-dihydro-1H-pyrazol-1-yl)-2-phenoxybenzene sulfonic acid (YL0325; CAS# 69053-03-8), 5-(5-Amino-3-methyl-1H-pyrazol-1-yl)-2-phenoxybenzene sulfonic acid (YL0327; CAS# 296793-70-9), and 5-(4-Ethyl-5-hydroxy-3-methyl-pyrazol-1-yl)-2-phenoxy-benzenesulfonic acid (YL0328; CAS# 321143-17-3) were obtained from ChemBridge (United States). The identity and purity of the target molecules were validated by NMR and HPLC analysis before their use in the studies.

*5-Nitro-2-phenoxybenzenesulfonic acid (2).* An oven-dried 300 mL single-neck round-bottom flask equipped with a magnetic stir bar was charged with 2-chloro-5-nitro-benzenesulfonic acid (1) (10.0 g, 42.0 mmol, 1.0 equiv.), phenol (4.75 g, 50.4 mmol, 1.2 equiv.), and $K_2CO_3$ (11.6 g, 84.0 mmol, 2.0 equiv.) in DMF (84 mL, 0.5 M) solvent. The reaction mixture was stirred at 100 °C for 24 h. Upon completion, the reaction mixture was quenched with 50 mL of brine solution. The reaction mixture was extracted with ethyl acetate (100 mL × 3 times), washed with water (100 mL × 3 times), and dried over anhydrous sodium sulfate. The solvent was removed under reduced pressure, and the crude product was purified by column chromatography on silica gel (70-230 mesh size) using a methanol/dichloromethane eluent (5:95 to 20:80 v/v) to obtain the desired product, 5-nitro-2-phenoxybenzenesulfonic acid (2) as a yellow solid (9.5 g, 77% yield). $^1$H NMR (400 MHz, DMSO-$d_6$) δ 8.62 (d, *J* = 3.00 Hz, 1H), 8.17 (dd, *J* = 9.01, 3.00 Hz, 1H), 7.41 - 7.56 (m, 2H), 7.21 - 7.32 (m, 1H), 7.05 - 7.18 (m, 2H), 6.88 (d, *J* = 9.01Hz, 1H), 3.19 (s, 1H). MS (ESI) m/z $[M - H]^+$ Calculated for $C_{12}H_8NO_6S$ 294.01; found 294.00.

*5-Amino-2-phenoxybenzenesulfonic acid (3).* An oven-dried 200 mL single-neck round-bottom flask equipped with a magnetic stir bar was charged with 5-nitro-2-phenoxybenzenesulfonic acid (2) (8.8 g,

30.0 mmol, 1.0 equiv.) in MeOH (60 mL, 0.5 M), and Pd/C (10 wt.% on activated carbon) (320 mg, 10 mol%) was added at room temperature. The flask was evacuated and refilled with a hydrogen balloon. The reaction mixture was stirred at room temperature for 24 h. After completion of the reaction (monitored by TLC), the reaction mixture was filtered through a bed of celite. The filtrate was concentrated *in vacuo*, and the crude product was purified by column chromatography on silica gel (70-230 mesh size) using 2% triethylamine (TEA) added methanol/dichloromethane as eluent (10:90 to 30:70 *v/v*) to obtain the desired product 5-amino-2-phenoxybenzenesulfonic acid (3) as a dark-yellow solid (6.5 g, 82% yield). $^1$H NMR (400 MHz, DMSO-$d_6$) δ 7.23 (t, *J* = 7.88 Hz, 2H), 7.10 (d, *J* = 2.63 Hz, 1H), 6.93 (t, *J* = 7.32 Hz, 1H), 6.82 (d, *J* = 7.88 Hz, 2H), 6.48 - 6.57 (m, 2H), 4.99 (s, 2H). MS (ESI) m/z $[M - H]^+$ Calculated for $C_{12}H_{10}NO_4S$ 264.03; found 264.02.

*5-Hydrazineyl-2-phenoxybenzenesulfonic acid (4).* An oven-dried 300 mL single-neck round bottom flask equipped with a magnetic stir bar was charged with 5-amino-2-phenoxy-benzenesulfonic acid (3) (5.6 g, 21.0 mmol, 1.0 equiv.) in 12% w/w dil. HCl (132 mL, 420 mmol, 20 equiv.). The reaction mixture was stirred at 0 °C for 30 minutes. Next, $NaNO_2$ solution (1.6 g, 23.2 0 mmol, 1.1 equiv.) in $H_2O$ (5 mL) was added dropwise at 0 °C; the mixture was stirred at 0 °C for 1 h, followed by a solution of $SnCl_2{\cdot}2H_2O$ (9.4 g, 42 mmol, 2.0 equiv.) in $H_2O$ (20 mL). The reaction mixture was stirred at 25 °C for 12 h and filtered using a Büchner funnel to obtain the desired product 5-hydrazineyl-2-phenoxybenzenesulfonic acid (4) as a light-yellow solid (6.0 g, 98% yield). $^1$H NMR (400 MHz, DMSO-d6) δ 9.84 - 10.10 (m, 3H), 7.82 - 8.31 (m, 1H), 7.55 (d, J = 2.81Hz, 1H), 7.30 (t, J = 7.95 Hz, 2H), 7.02 (t, J = 7.40 Hz, 1H), 6.89 - 6.93 (m, 1H), 6.85 - 6.89 (m, 2H), 6.79 - 6.84 (m, 1H). MS (ESI) m/z [M - H]+ Calculated for $C_{12}H_{12}N_2O_4S$ 279.04; found 279.00.

*5-(2-(tert-Butoxycarbonyl)hydrazineyl)-2-phenoxybenzenesulfonic acid (5).* An oven-dried 200 mL single-neck round-bottom flask equipped with a magnetic stir bar was charged with 5-hydrazineyl-2-phenoxybenzenesulfonic acid (4) (5.6 g, 20.0 mmol, 1.0 equiv.), TEA (3.1 g, 30.0 mmol, 1.5 equiv.) in a 1:1 mixture of *N,N*-dimethylformamide (DMF), and tetrahydrofuran (THF) (50 mL, 0.4 M) solvent. Next, Boc-anhydride ($Boc_2O$) (5.2 g, 24.0 mmol, 1.2 equiv.) was slowly added to the reaction mixture. After completion of the reaction (12 h, monitored by TLC), the reaction mixture was quenched with 50 mL of brine solution. The reaction mass was extracted with ethyl acetate (100 mL × 3) times), washed with water (50 mL × 3), and dried over anhydrous sodium sulfate. The reaction mixture was concentrated under reduced pressure to obtain the desired product 5-(2-(*tert*-butoxycarbonyl)hydrazineyl)-2-phenoxybenzenesulfonic acid (5) as a purple solid directly used in the next step (5.0 g, 65% yield). $^1$H NMR (400 MHz, DMSO-$d_6$) δ 9.11 - 9.20 (m, 1H), 8.79 (br. s, 1H), 7.51 - 7.57 (m, 1H), 7.21 - 7.27 (m, 3

H), 6.95 (t, *J* = 7.32 Hz, 1H), 6.82 (d, *J* = 8.00 Hz, 2H), 6.58 - 6.68 (m, 2H), 1.42 (s, 9 H). MS (ESI) m/z $[M - H]^+$ Calculated for $C_{17}H_{19}N_2O_6S$ 379.09; found 379.10.

*5-(2-(tert-Butoxycarbonyl)-1-(2-cyanoacetyl)hydrazineyl)-2-phenoxybenzenesulfonic acid (6).* An oven-dried 200 mL single-neck round-bottom flask equipped with a magnetic stir bar was charged with 5-(2-(*tert*-butoxycarbonyl)hydrazineyl)-2-phenoxybenzenesulfonic acid (5) (3.8 g, 10.0 mmol, 1.0 equiv.) and 2-cyanoacetic acid (1.3 g, 15.0 mmol, 1.5 equiv.) in dichloromethane (DCM) (50 mL, 0.2 M). Next, TEA (3.0 g, 30.0 mmol, 3.0 equiv.) and polyphosphoric acid anhydride (PPAA, 50% in ethyl acetate) (7.6 g, 12 mmol, 1.2 equiv.) were slowly added to the reaction mixture. The reaction mixture was stirred at room temperature for 2 h. After completion, the reaction mixture was concentrated under reduced pressure to obtain the desired product 5-(2-(*tert*-butoxycarbonyl)-1-(2-cyanoacetyl)hydrazineyl)-2-phenoxybenzenesulfonic acid (6) as a brown oil in a quantitative yield (5.5 g). The product was used directly in the subsequent step without further purification. MS (ESI) m/z $[M - H]^+$ Calculated for $C_{20}H_{20}N_3O_7S$ 446.10; found 446.00.

*5-(3-Amino-5-oxo-4,5-dihydro-1H-pyrazol-1-yl)-2-phenoxybenzenesulfonic acid (7, JPC0661).* An oven-dried 200 mL single-neck round-bottom flask equipped with a magnetic stir bar was charged with 5-(2-(*tert*-butoxycarbonyl)-1-(2-cyanoacetyl)hydrazineyl)-2-phenoxybenzenesulfonic acid (6) (4.5 g, 10.0 mmol, 1.0 equiv.) in 4 M hydrochloric acid solution in ethyl acetate (50 mL, 2.0 equiv.) solvent. The reaction mixture was stirred at room temperature for 2 h. The reaction mixture was filtered and concentrated under reduced pressure, and the crude product was purified by preparative HPLC using a Phenomenex Luna C18 column (250 mm × 100 mm × 10 µm). Water with 0.1% TFA and ACN (1-25% over 20 minutes as the gradient) was used as the mobile phases to obtain the desired product 5-(3-amino-5-oxo-4,5-dihydro-1H-pyrazol-1-yl)-2-phenoxybenzenesulfonic acid (7, JPC0661) as a white solid (2.5 g, 76% yield) with a purity of 97% as judged by HPLC. $^1$H NMR (800 MHz, DMSO-$d_6$) δ 8.17 (d, *J* = 2.8 Hz, 1H), 7.42 (dd, *J* = 8.9, 2.8 Hz, 1H), 7.34 (m, 1H), 7.30 (t, *J* = 7.7 Hz, 2H), 7.04 (t, *J* = 7.4 Hz, 1H), 6.96 (d, *J* = 8.1Hz, 1H), 6.91 (d, *J* = 8.1Hz, 1H), 6.83 (d, *J* = 8.8 Hz, 1H), 6.78 (d, *J* = 8.9 Hz, 1H). $^{13}$C NMR (201 MHz, DMSO-$d_6$) δ 168.8, 160.6, 158.4, 158.2, 157.5, 153.4, 151.7, 150.2, 138.6, 138.5, 133.9, 130.7, 130.5, 130.4, 125.7, 124.5, 124.1, 123.7, 123.3, 121.4, 121.0, 120.8, 120.1, 119.8, 119.5, 119.5. MS (ESI) m/z $[M - H]^+$ Calculated for $C_{15}H_{12}N_3O_5S$ 346.05; found 346.10.

**Table S1: JPC0661$_{comm}$ Molecule 1 (Lig1) Interactions ≤5.0 Å with the protein**

(sorted by Entity > Residue Nr > Distance), PDB ID: 9OC3. Distances were calculated with the program NCONT from the CCP4 package (1)

| AtomName | InteractingwithEntity | ResNr | ResName | AtomName | Dist[Å] |
|---|---|---|---|---|---|
| **SulfonicAcid** | | | | | |
| O20 | ΔFOSB | 171 | LYS | NZ | 2.80 |
| O19 | ΔFOSB | 171 | LYS | NZ | 3.35 |
| O19 | ΔFOSB | 171 | LYS | CE | 3.52 |
| S17 | ΔFOSB | 171 | LYS | NZ | 3.61 |
| O20 | ΔFOSB | 171 | LYS | CE | 3.63 |
| S17 | ΔFOSB | 171 | LYS | CE | 3.97 |
| O18 | ΔFOSB | 171 | LYS | CE | 4.37 |
| O18 | ΔFOSB | 171 | LYS | NZ | 4.39 |
| O19 | ΔFOSB | 171 | LYS | CD | 4.85 |
| O19 | ΔFOSB | 175 | ARG | NE | 2.78 |
| O18 | ΔFOSB | 175 | ARG | NH2 | 3.14 |
| O19 | ΔFOSB | 175 | ARG | CG | 3.46 |
| O19 | ΔFOSB | 175 | ARG | CD | 3.55 |
| O19 | ΔFOSB | 175 | ARG | CZ | 3.67 |
| O19 | ΔFOSB | 175 | ARG | NH2 | 3.68 |
| O18 | ΔFOSB | 175 | ARG | NE | 3.69 |
| S17 | ΔFOSB | 175 | ARG | NE | 3.76 |
| S17 | ΔFOSB | 175 | ARG | NH2 | 3.86 |
| O18 | ΔFOSB | 175 | ARG | CZ | 3.87 |
| S17 | ΔFOSB | 175 | ARG | CZ | 4.31 |
| O20 | ΔFOSB | 175 | ARG | NH2 | 4.60 |
| O19 | ΔFOSB | 175 | ARG | CB | 4.72 |
| O20 | ΔFOSB | 175 | ARG | NE | 4.78 |
| S17 | ΔFOSB | 175 | ARG | CD | 4.80 |
| S17 | ΔFOSB | 175 | ARG | CG | 4.89 |
| O19 | ΔFOSB | 175 | ARG | NH1 | 4.93 |
| O18 | ΔFOSB | 175 | ARG | CD | 4.97 |
| O19 | ΔFOSB | 411 | HOH | O | 2.80 |
| S17 | ΔFOSB | 411 | HOH | O | 3.72 |
| O20 | ΔFOSB | 411 | HOH | O | 3.88 |
| O19 | ΔFOSB | 428 | HOH | O | 4.06 |
| O20 | ΔFOSB | 428 | HOH | O | 4.48 |
| S17 | ΔFOSB | 428 | HOH | O | 4.95 |
| O19 | Ligand2 | 302 | A1C | O23 | 4.55 |
| **Aminopyrazolone** | | | | | |
| O23 | ΔFOSB | 403 | HOH | O | 3.22 |
| C22 | ΔFOSB | 403 | HOH | O | 3.70 |
| C24 | ΔFOSB | 403 | HOH | O | 4.18 |
| N4 | ΔFOSB | 403 | HOH | O | 4.61 |
| N1 | ΔFOSB | 435 | HOH | O | 2.91 |
| C24 | ΔFOSB | 435 | HOH | O | 3.68 |
| C2 | ΔFOSB | 435 | HOH | O | 3.71 |
| N3 | ΔFOSB | 435 | HOH | O | 4.98 |
| N1 | ΔFOSB | 447 | HOH | O | 3.36 |

| | | | | | |
|---|---|---|---|---|---|
| C2 | ΔFOSB | 447 | HOH | O | 3.83 |
| N3 | ΔFOSB | 447 | HOH | O | 4.06 |
| C24 | ΔFOSB | 447 | HOH | O | 4.80 |
| N1 | ΔFOSB | 454 | HOH | O | 3.82 |
| C24 | ΔFOSB | 454 | HOH | O | 4.29 |
| C2 | ΔFOSB | 454 | HOH | O | 4.32 |
| C2 | Ligand2 | 302 | A1C | N1 | 3.47 |
| N4 | Ligand2 | 302 | A1C | C2 | 3.48 |
| C24 | Ligand2 | 302 | A1C | N1 | 3.49 |
| N4 | Ligand2 | 302 | A1C | C24 | 3.59 |
| C22 | Ligand2 | 302 | A1C | C2 | 3.60 |
| C22 | Ligand2 | 302 | A1C | N1 | 3.67 |
| N3 | Ligand2 | 302 | A1C | N1 | 3.70 |
| N3 | Ligand2 | 302 | A1C | C24 | 3.71 |
| N4 | Ligand2 | 302 | A1C | N1 | 3.75 |
| N3 | Ligand2 | 302 | A1C | C2 | 3.76 |
| C22 | Ligand2 | 302 | A1C | N3 | 3.78 |
| O23 | Ligand2 | 302 | A1C | N3 | 3.82 |
| N4 | Ligand2 | 302 | A1C | N3 | 3.92 |
| C24 | Ligand2 | 302 | A1C | C2 | 3.93 |
| C2 | Ligand2 | 302 | A1C | C2 | 3.94 |
| N1 | Ligand2 | 302 | A1C | N1 | 3.94 |
| O23 | Ligand2 | 302 | A1C | C2 | 4.04 |
| N4 | Ligand2 | 302 | A1C | C22 | 4.13 |
| C22 | Ligand2 | 302 | A1C | C24 | 4.22 |
| N4 | Ligand2 | 302 | A1C | N4 | 4.27 |
| C2 | Ligand2 | 302 | A1C | C24 | 4.29 |
| O23 | Ligand2 | 302 | A1C | N1 | 4.30 |
| C24 | Ligand2 | 302 | A1C | N3 | 4.40 |
| O23 | Ligand2 | 302 | A1C | N4 | 4.41 |
| C22 | Ligand2 | 302 | A1C | N4 | 4.45 |
| N3 | Ligand2 | 302 | A1C | N3 | 4.59 |
| N3 | Ligand2 | 302 | A1C | C22 | 4.65 |
| C24 | Ligand2 | 302 | A1C | C24 | 4.69 |
| N1 | Ligand2 | 302 | A1C | C2 | 4.69 |
| C22 | Ligand2 | 302 | A1C | C22 | 4.71 |
| C2 | Ligand2 | 302 | A1C | N3 | 4.76 |
| O23 | Ligand2 | 302 | A1C | C24 | 4.79 |
| N4 | Ligand2 | 302 | A1C | O23 | 4.82 |
| O23 | Ligand2 | 302 | A1C | C22 | 4.96 |
| **Phenoxy** | | | | | |
| C12 | ΔFOSB | 175 | ARG | CD | 3.87 |
| C11 | ΔFOSB | 175 | ARG | CD | 3.94 |
| C11 | ΔFOSB | 175 | ARG | NE | 4.00 |
| C10 | ΔFOSB | 175 | ARG | NE | 4.01 |
| C13 | ΔFOSB | 175 | ARG | CD | 4.02 |
| C10 | ΔFOSB | 175 | ARG | CD | 4.14 |
| C13 | ΔFOSB | 175 | ARG | CB | 4.18 |
| C14 | ΔFOSB | 175 | ARG | CB | 4.18 |
| C14 | ΔFOSB | 175 | ARG | CG | 4.19 |
| C14 | ΔFOSB | 175 | ARG | CD | 4.22 |
| C15 | ΔFOSB | 175 | ARG | CD | 4.27 |
| O9 | ΔFOSB | 175 | ARG | NE | 4.28 |
| C14 | ΔFOSB | 175 | ARG | O | 4.29 |
| C13 | ΔFOSB | 175 | ARG | CG | 4.35 |

| | | | | | |
|---|---|---|---|---|---|
| C15 | ΔFOSB | 175 | ARG | CG | 4.36 |
| C12 | ΔFOSB | 175 | ARG | NE | 4.41 |
| C15 | ΔFOSB | 175 | ARG | NE | 4.44 |
| C13 | ΔFOSB | 175 | ARG | O | 4.53 |
| C14 | ΔFOSB | 175 | ARG | CA | 4.56 |
| C12 | ΔFOSB | 175 | ARG | CG | 4.65 |
| C10 | ΔFOSB | 175 | ARG | CG | 4.66 |
| C12 | ΔFOSB | 175 | ARG | CB | 4.78 |
| C15 | ΔFOSB | 175 | ARG | CB | 4.78 |
| C11 | ΔFOSB | 175 | ARG | CG | 4.80 |
| C13 | ΔFOSB | 175 | ARG | NE | 4.81 |
| C14 | ΔFOSB | 175 | ARG | NE | 4.82 |
| C14 | ΔFOSB | 175 | ARG | C | 4.82 |
| C11 | ΔFOSB | 175 | ARG | CZ | 4.83 |
| O9 | ΔFOSB | 175 | ARG | CD | 4.85 |
| C13 | ΔFOSB | 175 | ARG | CA | 4.88 |
| C10 | ΔFOSB | 175 | ARG | CZ | 4.98 |
| C13 | ΔFOSB | 175 | ARG | C | 4.98 |
| C14 | ΔFOSB | 178 | GLU | CB | 4.91 |
| C13 | ΔFOSB | 179 | LEU | CD2 | 4.09 |
| C14 | ΔFOSB | 179 | LEU | CD2 | 4.14 |
| C13 | ΔFOSB | 179 | LEU | CG | 4.17 |
| C13 | ΔFOSB | 179 | LEU | CD1 | 4.22 |
| C14 | ΔFOSB | 179 | LEU | CG | 4.34 |
| C14 | ΔFOSB | 179 | LEU | CD1 | 4.84 |
| C15 | ΔFOSB | 411 | HOH | O | 3.58 |
| C14 | ΔFOSB | 411 | HOH | O | 4.07 |
| C10 | ΔFOSB | 411 | HOH | O | 4.49 |
| O9 | ΔFOSB | 411 | HOH | O | 4.74 |
| C15 | Ligand2 | 302 | A1C | O23 | 3.35 |
| C14 | Ligand2 | 302 | A1C | O23 | 3.79 |
| C15 | Ligand2 | 302 | A1C | C6 | 4.11 |
| C14 | Ligand2 | 302 | A1C | C6 | 4.12 |
| C15 | Ligand2 | 302 | A1C | C22 | 4.37 |
| C10 | Ligand2 | 302 | A1C | O23 | 4.54 |
| C14 | Ligand2 | 302 | A1C | C7 | 4.55 |
| C15 | Ligand2 | 302 | A1C | C7 | 4.82 |
| O9 | Ligand2 | 302 | A1C | O23 | 4.85 |
| C14 | Ligand2 | 302 | A1C | C22 | 4.91 |

**Centralphenyl**

| | | | | | |
|---|---|---|---|---|---|
| C6 | ΔFOSB | 403 | HOH | O | 4.90 |
| C16 | ΔFOSB | 411 | HOH | O | 3.85 |
| C21 | ΔFOSB | 411 | HOH | O | 4.17 |
| C8 | ΔFOSB | 411 | HOH | O | 4.31 |
| C5 | ΔFOSB | 411 | HOH | O | 4.85 |
| C7 | ΔFOSB | 411 | HOH | O | 4.97 |
| C5 | Ligand2 | 302 | A1C | C24 | 3.64 |
| C6 | Ligand2 | 302 | A1C | C22 | 3.71 |
| C6 | Ligand2 | 302 | A1C | N4 | 3.72 |
| C5 | Ligand2 | 302 | A1C | C22 | 3.74 |
| C5 | Ligand2 | 302 | A1C | C2 | 3.94 |
| C6 | Ligand2 | 302 | A1C | O23 | 3.97 |
| C7 | Ligand2 | 302 | A1C | O23 | 3.97 |
| C21 | Ligand2 | 302 | A1C | C24 | 4.02 |

| | | | | | |
|---|---|---|---|---|---|
| C5 | Ligand2 | 302 | A1C | N4 | 4.06 |
| C6 | Ligand2 | 302 | A1C | N3 | 4.09 |
| C5 | Ligand2 | 302 | A1C | O23 | 4.10 |
| C8 | Ligand2 | 302 | A1C | O23 | 4.10 |
| C6 | Ligand2 | 302 | A1C | C24 | 4.11 |
| C7 | Ligand2 | 302 | A1C | C22 | 4.13 |
| C21 | Ligand2 | 302 | A1C | C22 | 4.16 |
| C5 | Ligand2 | 302 | A1C | N3 | 4.20 |
| C16 | Ligand2 | 302 | A1C | O23 | 4.21 |
| C21 | Ligand2 | 302 | A1C | O23 | 4.21 |
| C6 | Ligand2 | 302 | A1C | C5 | 4.26 |
| C6 | Ligand2 | 302 | A1C | C2 | 4.26 |
| C7 | Ligand2 | 302 | A1C | N4 | 4.28 |
| C7 | Ligand2 | 302 | A1C | C6 | 4.45 |
| C8 | Ligand2 | 302 | A1C | C22 | 4.53 |
| C16 | Ligand2 | 302 | A1C | C22 | 4.54 |
| C7 | Ligand2 | 302 | A1C | C5 | 4.56 |
| C6 | Ligand2 | 302 | A1C | C6 | 4.58 |
| C5 | Ligand2 | 302 | A1C | N1 | 4.59 |
| C21 | Ligand2 | 302 | A1C | C2 | 4.72 |
| C16 | Ligand2 | 302 | A1C | C24 | 4.77 |
| C7 | Ligand2 | 302 | A1C | C24 | 4.84 |
| C21 | Ligand2 | 302 | A1C | N4 | 4.85 |
| C5 | Ligand2 | 302 | A1C | C5 | 4.97 |

**Table S1, cont.: JPC0661$_{comm}$ Molecule 2 (Lig2) Interactions ≤5.0 Å with the protein**

(sorted by Entity > Residue Nr > Distance). Distances were calculated with the program NCONT from the CCP4 package (1).

| AtomName | InteractingwithEntity | ResNr | ResName | AtomName | Dist[Å] |
|---|---|---|---|---|---|
| **SulfonicAcid** | | | | | |
| O18 | ΔFOSB | 182 | ARG | CD | 3.58 |
| O20 | ΔFOSB | 182 | ARG | NH1 | 3.88 |
| O18 | ΔFOSB | 182 | ARG | NH1 | 4.08 |
| S17 | ΔFOSB | 182 | ARG | CD | 4.22 |
| O18 | ΔFOSB | 182 | ARG | CB | 4.27 |
| O18 | ΔFOSB | 182 | ARG | CG | 4.28 |
| S17 | ΔFOSB | 182 | ARG | NH1 | 4.33 |
| O18 | ΔFOSB | 182 | ARG | NE | 4.59 |
| O20 | ΔFOSB | 182 | ARG | CZ | 4.73 |
| O20 | ΔFOSB | 182 | ARG | CD | 4.73 |
| O18 | ΔFOSB | 182 | ARG | CZ | 4.78 |
| S17 | ΔFOSB | 182 | ARG | NE | 4.93 |
| S17 | ΔFOSB | 182 | ARG | CZ | 4.98 |
| O20 | ΔFOSB | 403 | HOH | O | 3.55 |
| S17 | ΔFOSB | 403 | HOH | O | 4.60 |
| O18 | ΔFOSB | 438 | HOH | O | 4.44 |
| **Aminopyrazolone** | | | | | |
| O23 | ΔFOSB | 178 | GLU | CB | 3.50 |
| O23 | ΔFOSB | 178 | GLU | CG | 3.56 |
| C24 | ΔFOSB | 178 | GLU | CD | 3.85 |
| C24 | ΔFOSB | 178 | GLU | OE1 | 3.94 |
| C22 | ΔFOSB | 178 | GLU | OE1 | 3.94 |
| O23 | ΔFOSB | 178 | GLU | CD | 3.96 |
| C22 | ΔFOSB | 178 | GLU | CD | 3.97 |
| C24 | ΔFOSB | 178 | GLU | OE2 | 3.98 |
| C22 | ΔFOSB | 178 | GLU | CG | 4.05 |
| O23 | ΔFOSB | 178 | GLU | OE1 | 4.14 |
| C22 | ΔFOSB | 178 | GLU | CB | 4.15 |
| C24 | ΔFOSB | 178 | GLU | CG | 4.33 |
| N4 | ΔFOSB | 178 | GLU | OE1 | 4.44 |
| C2 | ΔFOSB | 178 | GLU | OE1 | 4.50 |
| C22 | ΔFOSB | 178 | GLU | OE2 | 4.52 |
| O23 | ΔFOSB | 178 | GLU | OE2 | 4.63 |
| C2 | ΔFOSB | 178 | GLU | CD | 4.75 |
| N3 | ΔFOSB | 178 | GLU | OE1 | 4.81 |
| N4 | ΔFOSB | 178 | GLU | CD | 4.82 |
| C24 | ΔFOSB | 178 | GLU | CB | 4.88 |
| C2 | ΔFOSB | 178 | GLU | OE2 | 4.90 |
| N4 | ΔFOSB | 178 | GLU | CB | 4.91 |
| O23 | ΔFOSB | 178 | GLU | CA | 4.95 |
| C2 | ΔFOSB | 182 | ARG | NH2 | 3.42 |
| N3 | ΔFOSB | 182 | ARG | NH2 | 3.44 |
| N1 | ΔFOSB | 182 | ARG | NH2 | 3.57 |
| N3 | ΔFOSB | 182 | ARG | CZ | 3.58 |

| | | | | | |
|---|---|---|---|---|---|
| N4 | ΔFOSB | 182 | ARG | NE | 3.72 |
| N4 | ΔFOSB | 182 | ARG | CZ | 3.84 |
| N4 | ΔFOSB | 182 | ARG | NH2 | 3.91 |
| N3 | ΔFOSB | 182 | ARG | NE | 3.97 |
| C24 | ΔFOSB | 182 | ARG | NH2 | 3.97 |
| C2 | ΔFOSB | 182 | ARG | CZ | 4.02 |
| N3 | ΔFOSB | 182 | ARG | NH1 | 4.05 |
| C22 | ΔFOSB | 182 | ARG | NE | 4.21 |
| C22 | ΔFOSB | 182 | ARG | NH2 | 4.25 |
| N1 | ΔFOSB | 182 | ARG | CZ | 4.42 |
| N4 | ΔFOSB | 182 | ARG | CD | 4.45 |
| C22 | ΔFOSB | 182 | ARG | CZ | 4.45 |
| C2 | ΔFOSB | 182 | ARG | NE | 4.48 |
| N4 | ΔFOSB | 182 | ARG | NH1 | 4.52 |
| C24 | ΔFOSB | 182 | ARG | CZ | 4.59 |
| C2 | ΔFOSB | 182 | ARG | NH1 | 4.68 |
| O23 | ΔFOSB | 182 | ARG | NE | 4.71 |
| C24 | ΔFOSB | 182 | ARG | NE | 4.71 |
| N3 | ΔFOSB | 182 | ARG | CD | 4.86 |
| N1 | ΔFOSB | 182 | ARG | NH1 | 4.94 |
| N3 | ΔFOSB | 403 | HOH | O | 2.43 |
| C2 | ΔFOSB | 403 | HOH | O | 3.29 |
| N1 | ΔFOSB | 403 | HOH | O | 3.50 |
| N4 | ΔFOSB | 403 | HOH | O | 3.52 |
| C24 | ΔFOSB | 403 | HOH | O | 4.62 |
| C22 | ΔFOSB | 403 | HOH | O | 4.66 |
| N1 | ΔFOSB | 406 | HOH | O | 3.80 |
| C2 | ΔFOSB | 406 | HOH | O | 4.03 |
| C24 | ΔFOSB | 406 | HOH | O | 4.04 |
| N3 | ΔFOSB | 406 | HOH | O | 4.78 |
| C22 | ΔFOSB | 406 | HOH | O | 4.90 |
| O23 | ΔFOSB | 411 | HOH | O | 2.62 |
| C22 | ΔFOSB | 411 | HOH | O | 3.48 |
| C24 | ΔFOSB | 411 | HOH | O | 3.81 |
| N4 | ΔFOSB | 411 | HOH | O | 4.80 |
| N1 | ΔFOSB | 435 | HOH | O | 4.86 |
| N1 | ΔFOSB | 447 | HOH | O | 3.15 |
| C24 | ΔFOSB | 447 | HOH | O | 3.90 |
| C2 | ΔFOSB | 447 | HOH | O | 3.95 |
| N1 | ΔFOSB | 454 | HOH | O | 3.50 |
| C2 | ΔFOSB | 454 | HOH | O | 4.80 |
| N1 | ΔFOSB | 459 | HOH | O | 3.60 |
| C2 | ΔFOSB | 459 | HOH | O | 4.39 |
| N3 | ΔFOSB | 459 | HOH | O | 4.45 |
| O23 | Ligand1 | 301 | A1C | C15 | 3.35 |
| N1 | Ligand1 | 301 | A1C | C2 | 3.47 |
| C2 | Ligand1 | 301 | A1C | N4 | 3.48 |
| N1 | Ligand1 | 301 | A1C | C24 | 3.49 |
| C24 | Ligand1 | 301 | A1C | N4 | 3.59 |
| C2 | Ligand1 | 301 | A1C | C22 | 3.60 |
| C24 | Ligand1 | 301 | A1C | C5 | 3.64 |
| N1 | Ligand1 | 301 | A1C | C22 | 3.67 |
| N1 | Ligand1 | 301 | A1C | N3 | 3.70 |
| C24 | Ligand1 | 301 | A1C | N3 | 3.71 |
| C22 | Ligand1 | 301 | A1C | C6 | 3.71 |
| N4 | Ligand1 | 301 | A1C | C6 | 3.72 |

| | | | | | |
|---|---|---|---|---|---|
| C22 | Ligand1 | 301 | A1C | C5 | 3.74 |
| N1 | Ligand1 | 301 | A1C | N4 | 3.75 |
| C2 | Ligand1 | 301 | A1C | N3 | 3.76 |
| N3 | Ligand1 | 301 | A1C | C22 | 3.78 |
| O23 | Ligand1 | 301 | A1C | C14 | 3.79 |
| N3 | Ligand1 | 301 | A1C | O23 | 3.82 |
| N3 | Ligand1 | 301 | A1C | N4 | 3.92 |
| C2 | Ligand1 | 301 | A1C | C24 | 3.93 |
| C2 | Ligand1 | 301 | A1C | C2 | 3.94 |
| N1 | Ligand1 | 301 | A1C | N1 | 3.94 |
| C2 | Ligand1 | 301 | A1C | C5 | 3.94 |
| O23 | Ligand1 | 301 | A1C | C6 | 3.97 |
| O23 | Ligand1 | 301 | A1C | C7 | 3.97 |
| C24 | Ligand1 | 301 | A1C | C21 | 4.02 |
| C2 | Ligand1 | 301 | A1C | O23 | 4.04 |
| N4 | Ligand1 | 301 | A1C | C5 | 4.06 |
| N3 | Ligand1 | 301 | A1C | C6 | 4.09 |
| O23 | Ligand1 | 301 | A1C | C5 | 4.10 |
| O23 | Ligand1 | 301 | A1C | C8 | 4.10 |
| C24 | Ligand1 | 301 | A1C | C6 | 4.11 |
| C22 | Ligand1 | 301 | A1C | C7 | 4.13 |
| C22 | Ligand1 | 301 | A1C | N4 | 4.13 |
| C22 | Ligand1 | 301 | A1C | C21 | 4.16 |
| N3 | Ligand1 | 301 | A1C | C5 | 4.20 |
| O23 | Ligand1 | 301 | A1C | C16 | 4.21 |
| O23 | Ligand1 | 301 | A1C | C21 | 4.21 |
| C24 | Ligand1 | 301 | A1C | C22 | 4.22 |
| C2 | Ligand1 | 301 | A1C | C6 | 4.26 |
| N4 | Ligand1 | 301 | A1C | N4 | 4.27 |
| N4 | Ligand1 | 301 | A1C | C7 | 4.28 |
| C24 | Ligand1 | 301 | A1C | C2 | 4.29 |
| N1 | Ligand1 | 301 | A1C | O23 | 4.30 |
| C22 | Ligand1 | 301 | A1C | C15 | 4.37 |
| N3 | Ligand1 | 301 | A1C | C24 | 4.40 |
| N4 | Ligand1 | 301 | A1C | O23 | 4.41 |
| N4 | Ligand1 | 301 | A1C | C22 | 4.45 |
| C22 | Ligand1 | 301 | A1C | C8 | 4.53 |
| C22 | Ligand1 | 301 | A1C | C16 | 4.54 |
| O23 | Ligand1 | 301 | A1C | C10 | 4.54 |
| O23 | Ligand1 | 301 | A1C | O19 | 4.55 |
| N3 | Ligand1 | 301 | A1C | N3 | 4.59 |
| N1 | Ligand1 | 301 | A1C | C5 | 4.59 |
| C22 | Ligand1 | 301 | A1C | N3 | 4.65 |
| C2 | Ligand1 | 301 | A1C | N1 | 4.69 |
| C24 | Ligand1 | 301 | A1C | C24 | 4.69 |
| C22 | Ligand1 | 301 | A1C | C22 | 4.71 |
| C2 | Ligand1 | 301 | A1C | C21 | 4.72 |
| N3 | Ligand1 | 301 | A1C | C2 | 4.76 |
| C24 | Ligand1 | 301 | A1C | C16 | 4.77 |
| C24 | Ligand1 | 301 | A1C | O23 | 4.79 |
| O23 | Ligand1 | 301 | A1C | N4 | 4.82 |
| C24 | Ligand1 | 301 | A1C | C7 | 4.84 |
| N4 | Ligand1 | 301 | A1C | C21 | 4.85 |
| O23 | Ligand1 | 301 | A1C | O9 | 4.85 |
| C22 | Ligand1 | 301 | A1C | C14 | 4.91 |
| C22 | Ligand1 | 301 | A1C | O23 | 4.96 |

| | | | | | |
|---|---|---|---|---|---|
| **Phenoxy** | | | | | |
| C15 | ΔFOSB | 178 | GLU | O | 4.84 |
| C14 | ΔFOSB | 178 | GLU | O | 5.00 |
| C15 | ΔFOSB | 179 | LEU | CD2 | 3.75 |
| C14 | ΔFOSB | 179 | LEU | CD2 | 3.75 |
| C14 | ΔFOSB | 179 | LEU | O | 3.76 |
| C14 | ΔFOSB | 179 | LEU | CA | 4.00 |
| C10 | ΔFOSB | 179 | LEU | CD2 | 4.11 |
| C13 | ΔFOSB | 179 | LEU | CD2 | 4.11 |
| C14 | ΔFOSB | 179 | LEU | C | 4.34 |
| C14 | ΔFOSB | 179 | LEU | CB | 4.34 |
| C15 | ΔFOSB | 179 | LEU | CA | 4.39 |
| C12 | ΔFOSB | 179 | LEU | CD2 | 4.44 |
| C11 | ΔFOSB | 179 | LEU | CD2 | 4.44 |
| C15 | ΔFOSB | 179 | LEU | O | 4.57 |
| C13 | ΔFOSB | 179 | LEU | O | 4.60 |
| C14 | ΔFOSB | 179 | LEU | CG | 4.70 |
| C13 | ΔFOSB | 179 | LEU | CB | 4.83 |
| O9 | ΔFOSB | 179 | LEU | CD2 | 4.86 |
| C13 | ΔFOSB | 179 | LEU | CA | 4.87 |
| C15 | ΔFOSB | 179 | LEU | CB | 4.88 |
| C15 | ΔFOSB | 179 | LEU | CG | 4.94 |
| C15 | ΔFOSB | 182 | ARG | CB | 3.89 |
| C14 | ΔFOSB | 182 | ARG | CB | 4.11 |
| C15 | ΔFOSB | 182 | ARG | CD | 4.19 |
| C15 | ΔFOSB | 182 | ARG | CG | 4.63 |
| O9 | ΔFOSB | 182 | ARG | CD | 4.73 |
| C14 | ΔFOSB | 182 | ARG | C | 4.89 |
| C10 | ΔFOSB | 182 | ARG | CD | 4.98 |
| C13 | ΔFOSB | 183 | LEU | CG | 3.97 |
| C14 | ΔFOSB | 183 | LEU | CG | 4.00 |
| C13 | ΔFOSB | 183 | LEU | CD2 | 4.03 |
| C13 | ΔFOSB | 183 | LEU | CD1 | 4.28 |
| C14 | ΔFOSB | 183 | LEU | CD2 | 4.40 |
| C14 | ΔFOSB | 183 | LEU | CD1 | 4.51 |
| C14 | ΔFOSB | 183 | LEU | N | 4.64 |
| C13 | JUND | 297 | GLU | OE2 | 4.83 |
| C12 | JUND | 426 | HOH | O | 4.65 |
| C13 | JUND | 426 | HOH | O | 4.81 |
| C12 | JUND | 428 | HOH | O | 3.78 |
| C13 | JUND | 428 | HOH | O | 3.91 |
| **Centralphenyl** | | | | | |
| C6 | ΔFOSB | 178 | GLU | CB | 4.70 |
| C7 | ΔFOSB | 179 | LEU | CD2 | 3.85 |
| C6 | ΔFOSB | 179 | LEU | CD2 | 4.54 |
| C8 | ΔFOSB | 179 | LEU | CD2 | 4.75 |
| C5 | ΔFOSB | 182 | ARG | NE | 3.59 |
| C21 | ΔFOSB | 182 | ARG | CD | 3.60 |
| C21 | ΔFOSB | 182 | ARG | NE | 3.61 |
| C16 | ΔFOSB | 182 | ARG | CD | 3.64 |
| C21 | ΔFOSB | 182 | ARG | CZ | 3.76 |
| C21 | ΔFOSB | 182 | ARG | NH1 | 3.80 |
| C5 | ΔFOSB | 182 | ARG | CD | 3.83 |
| C8 | ΔFOSB | 182 | ARG | CD | 3.94 |

| | | | | | |
|---|---|---|---|---|---|
| C5 | ΔFOSB | 182 | ARG | CZ | 3.97 |
| C6 | ΔFOSB | 182 | ARG | CD | 4.11 |
| C7 | ΔFOSB | 182 | ARG | CD | 4.15 |
| C6 | ΔFOSB | 182 | ARG | NE | 4.15 |
| C16 | ΔFOSB | 182 | ARG | NE | 4.18 |
| C16 | ΔFOSB | 182 | ARG | NH1 | 4.35 |
| C5 | ΔFOSB | 182 | ARG | NH1 | 4.44 |
| C5 | ΔFOSB | 182 | ARG | NH2 | 4.49 |
| C16 | ΔFOSB | 182 | ARG | CZ | 4.52 |
| C21 | ΔFOSB | 182 | ARG | NH2 | 4.52 |
| C7 | ΔFOSB | 182 | ARG | NE | 4.65 |
| C8 | ΔFOSB | 182 | ARG | NE | 4.68 |
| C6 | ΔFOSB | 182 | ARG | CZ | 4.88 |
| C16 | ΔFOSB | 182 | ARG | CG | 4.99 |
| C21 | ΔFOSB | 403 | HOH | O | 3.21 |
| C5 | ΔFOSB | 403 | HOH | O | 3.79 |
| C16 | ΔFOSB | 403 | HOH | O | 4.25 |
| C6 | Ligand1 | 301 | A1C | C15 | 4.11 |
| C6 | Ligand1 | 301 | A1C | C14 | 4.12 |
| C5 | Ligand1 | 301 | A1C | C6 | 4.26 |
| C6 | Ligand1 | 301 | A1C | C7 | 4.45 |
| C7 | Ligand1 | 301 | A1C | C14 | 4.55 |
| C5 | Ligand1 | 301 | A1C | C7 | 4.56 |
| C6 | Ligand1 | 301 | A1C | C6 | 4.58 |
| C7 | Ligand1 | 301 | A1C | C15 | 4.82 |
| C5 | Ligand1 | 301 | A1C | C5 | 4.97 |
